# Cortex-Wide, Cellular-Resolution Volumetric Imaging with a Modular Two-Photon Imaging Platform

**DOI:** 10.1101/2025.07.03.662899

**Authors:** Jiahao Hu, Yanfeng Zhu, Shoupei Liu, Chengyu Li, Min Zhang, Haidong Lv, Huiyun Yu, Xinyang Gu, Jingchuan Wu, Fang Xu, Liang Chen, Ying Mao, Bo Li

## Abstract

Mapping cortex-wide neuronal activity at single-cell resolution has been limited by the physical trade-off between numerical aperture and field-of-view (FOV) in two-photon microscopes. We present Meso2P, a modular two-photon platform that decouples excitation and detection by introducing a lateral paraboloid fluorescence collector. Ray-tracing predicted collection of approximately 51% of the upper-hemisphere emission, and relative radiometric measurements showed a 13.7-fold gain over the 4×/0.28-NA objective path under the tested geometry, while supporting a contiguous 6 × 6 mm² FOV at high speed (2,048 × 2,048 pixels at 7.67 Hz). The modular platform can be upgraded with optional modules for simultaneous multi-plane imaging (1-4 planes at full resolution and speed), volumetric imaging (6 × 6 × 0.5 mm^3^, 2,048 × 2,048 × 28 voxels at 1.1 Hz capturing > 210,000 neurons), and spatially distributed two-photon optogenetic stimulation across multiple brain regions. To handle the resulting large-scale data, we provide an open-source analysis platform that integrates motion correction, denoising, segmentation and spike inference. We demonstrate cortex-wide sensory responses, layer-specific network synchrony during anaesthesia, and in vivo tracking of micro- and nanoplastic distribution. Meso2P therefore provides a reproducible route to high-throughput volumetric imaging across almost the entire cortex with high detection efficiency.

## Introduction

Understanding how spatially distributed cortical circuits cooperate to generate perception and behaviour^1–3^ requires imaging tools that record activity from multiple brain regions simultaneously and with single-cell resolution^4^. Two-photon (2P) microscopy offers intrinsic optical sectioning, micron-scale resolution and compatibility with genetically encoded indicators, and has therefore become the workhorse of cellular neurophysiology^5–12^. Yet conventional designs image less than 1 mm² per frame, obliging researchers to trade breadth for detail and to rely on sequential recordings when probing cortex-wide dynamics^13^.

Large-field-of-view (FOV) mesoscopes have begun to relax this constraint by scaling scan angles and using custom objectives with entrance pupils exceeding 25 mm. These instruments can span several millimetres, but the required long focal lengths inevitably lower their numerical aperture (NA ≈ 0.4-0.5)^13–20^, reducing fluorescence collection efficiency, compromising deep-tissue sensitivity^21^ and introducing field curvature^22^. Add-on volumetric modules—Bessel beams^23^, spatial multiplexing^24^, or light-field optics^17^—restore 3D coverage but inherit the parent system’s limited NA and are often incompatible with 2P optogenetic stimulation. In parallel, the massive data streams (> 100 GB h⁻¹) produced by large-FOV imaging strain existing analysis pipelines, which were designed for thousands rather than hundreds of thousands of neurons^17, 25–27^.

At the heart of these challenges lies a simple physical coupling in single-objective designs: when excitation and detection share the same objective, NA × FOV is approximately proportional to the objective diameter (D ≈ 2 × *f* × NA). As FOV grows, maintaining a high NA requires impractically large optics and scan apertures. Because fluorescence collection efficiency is strongly related to collection NA and collection solid angle, the reduced NA of large-FOV objectives becomes a fundamental limitation for mesoscale 2P imaging, especially for high-throughput and volumetric recordings.

Here, we introduce Meso2P, a modular 2P platform that addresses this collection-efficiency bottleneck by physically separating excitation and detection. Its key advances are: first, by adapting supplementary fluorescence collection^28^ into large-FOV 2P microscopy, we designed a paraboloid lateral collection module that provides high fluorescence collection efficiency across a 6 × 6 mm² cortical window, corresponding to an equivalent collection NA of 0.87, while excitation is delivered through a 4× objective that provides lateral cellular resolution at high speed (2,048 × 2,048 pixels at 7.67 Hz); second, a modular imaging architecture supports simultaneous multi-plane imaging (1-4 planes at full resolution and speed), volumetric imaging (6 × 6 × 0.5 mm^3^, 2,048 × 2,048 × 28 voxels at 1.1 Hz), and 2P optogenetic stimulation; third, an open-source integrated analysis platform (NeuroPixelAI) motion-corrects, segments and infers calcium signals from > 200,000 neurons in 100 GB datasets on a standard workstation. Together, these features support high-throughput imaging, volumetric recording and multi-region photostimulation within a modular mesoscopic platform.

## Results

### Implementation of Meso2P for High-Efficiency, Mesoscale Two-Photon Imaging

Meso2P integrates four plug-and-play modules for a mesoscale, cellular-resolution imaging and perturbation platform (Fig. 1): (1) a custom wide-angle scan relay for micron-resolution imaging across a 6 × 6 mm^2^ FOV; (2) a lateral paraboloid fluorescence collection module that decouples detection from excitation and provides 51% simulated upper-hemisphere collection (solid-angle-equivalent collection NA, 0.87); (3) a multiplexing module that enables simultaneous multi-plane imaging, which can be further upgraded to volumetric imaging by adding a piezo-driven axial scanning stage; (4) a 2P optogenetics module for cross-regional perturbation.

**Fig. 1.**
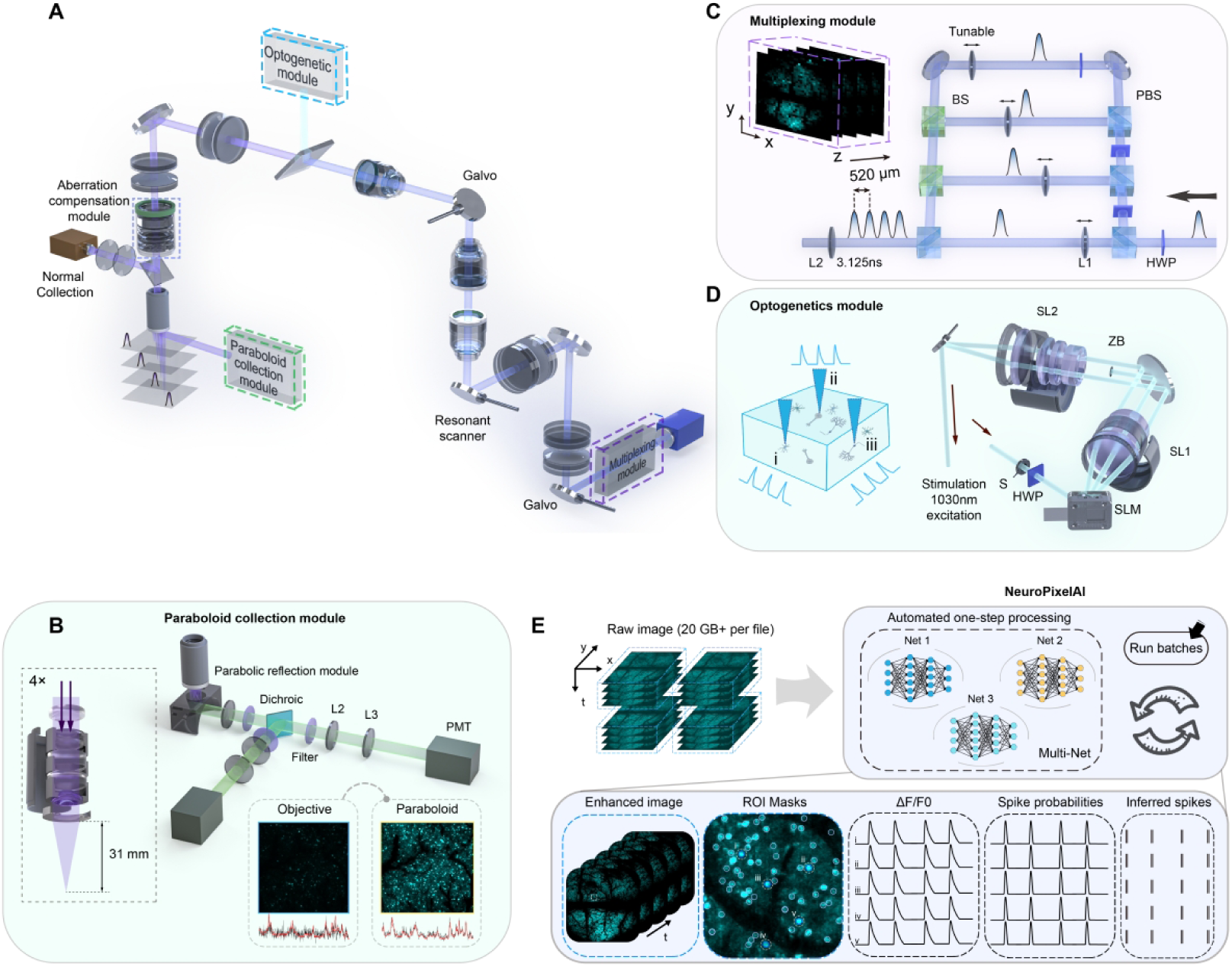
Schematic overview of the Meso2P system. **(A)** Optical schematic of Meso2P. **(B)** Working principle of the lateral paraboloid collection module. **(C)** Multi-plane free imaging enabled by a spatiotemporal multiplexing module combined with piezoelectric z-scanning. **(D)** Optical layout of the SLM-based 2P holographic stimulation module. S, shutter; HWP, half-wave plate; SL1-2, scan lenses; SLM, spatial light modulator; ZB, zero order block. **(E)** NeuroPixelAI, an integrated deep learning-based analysis pipeline for high-throughput segmentation and calcium signal extraction from mesoscale 2P datasets. ***See also* Extended Data Figs. 1-3 and Supplementary Video 1**.

The custom scan relay was designed for near-diffraction-limited, uniform imaging over a near-centimetre-scale field. Galvo-resonant mirrors scanning through ± 13° are relayed by a custom-designed optics chain comprising several scan lenses, a tube lens, an aberration compensation module, and a 4× objective (Fig. 1A, Extended Data Fig. 1A and Supplementary Video 1). Wavefront errors remain < 0.1 λ RMS throughout the scan range, with uniform lateral resolution and brightness across the entire FOV (Extended Data Fig. 1B,C). Unlike widely used mesoscopes that exhibit pronounced field curvature (∼ 66.7 µm/mm)^22^, Meso2P maintains low field curvature (∼ 5.57 µm/mm) as validated by uniform fluorescence from thin ruler imaging (Extended Data Fig. 1D). The resulting configuration achieves lateral and axial resolutions of 1.2-1.8 µm and 21-24 µm, respectively, across a 6 × 6 mm² FOV (Extended Data Fig. 1E). Alternative objectives, such as a commercial 16×, can be integrated to flexibly trade spatial resolution for imaging area (Extended Data Fig. 1F,G).

Decoupling fluorescence collection from excitation relaxes the collection-efficiency constraint imposed by the NA-FOV coupling, significantly enhancing photon collection. Conventional large-FOV 2P systems enlarge objective diameters to increase NA, yet remain fundamentally constrained by the NA × FOV relationship (D ≈ 2 × *f* × NA). Meso2P addresses this collection-efficiency limitation by introducing an aluminium-coated paraboloid fluorescence collection module positioned laterally between the objective and sample (Fig. 1B, Extended Data Fig. 2A). Excitation via the 4× objective at NA 0.28 provides adequate micron-level resolution for large-scale imaging, while fluorescence collection benefits from an equivalent collection NA of 0.87 across a 6 × 6 mm^2^ area (Extended Data Fig. 2B-F), while leaving the 0.28 excitation NA and the associated axial sectioning unchanged. Uniformity in point-spread-function peak intensity and collection efficiency (< 10% variation) ensures consistent high-quality imaging without oversized optics (Extended Data Fig. 2G). The resulting improvement in image quality translates into better segmentation fidelity and higher-quality extracted neuronal signals (Extended Data Fig. 2H). A summary of Meso2P operating specifications is provided in Supplementary Table 1, while descriptive cross-platform comparisons of volumetric and optical/acquisition parameters are provided in Supplementary Tables 2 and 3, respectively. Further details are provided in Supplementary Note 1.

The multiplexing module enables high-throughput, simultaneous multi-plane imaging. Each laser pulse is divided into four temporally separated sub-pulses (∼ 3.125 ns apart), simultaneously exciting multiple focal planes separated by ∼ 130 µm, adjustable to cortical layers of interest (Fig. 1C, Extended Data Fig. 3A,B and Supplementary Note 2). Crucially, the inter-plane spacing is adjustable, allowing precise targeting of anatomically defined cortical layers, such as layers 2/3 and 5, and enabling concurrent imaging of functionally distinct networks at different depths. This flexible multi-plane capability supports simultaneous interrogation of anatomically defined cortical layers across distributed brain regions. Furthermore, the system can be upgraded to volumetric imaging by combining four-plane multiplexing with piezo-driven axial scanning. Piezo-driven axial scanning further divides each plane into seven axial steps (18.5 µm spacing), creating a 6 × 6 × 0.5 mm^3^ volume (3 × 3 × 18.5 µm^3^ voxel size) imaged at 1.1 Hz (Extended Data Fig. 3C,D). Combining efficient single-point excitation with high-efficiency lateral fluorescence collection enables stable imaging at a total laser power of approximately 250 mW, which is lower than the power required by previous systems^24^. While elevated laser power can improve imaging fidelity, it can also increase tissue stress and photodamage risk within the imaged region. We quantitatively compared the extent of photodamage across a laser power range of 0-450 mW via immunofluorescence imaging. We therefore used approximately 250 mW for high-quality image acquisition to limit potential phototoxicity in both single-plane and simultaneous multi-plane volumetric imaging (Extended Data Fig. 4, Supplementary Note 3). Temporal crosstalk is lower when fewer planes are multiplexed; in the four-plane mode, adjacent-channel contamination is measurable and is reduced by an offline demixing process followed by analysis with the NeuroPixelAI pipeline (Extended Data Fig. 5 and Supplementary Note 4). Spatial signal overlap between neighbouring imaging planes is consistent with the measured axial PSF and the 18.5-µm inter-plane spacing (Extended Data Fig. 6).

The integrated 2P optogenetic module supports spatially registered perturbation. A spatial-light modulator generates user-defined holographic stimulation patterns that are scanned by galvanometric mirrors in spiral trajectories (Fig. 1D and Extended Data Fig. 7). Imaging and stimulation beams are co-registered through a dichroic mirror; validation under 4× and 16× objectives is reported in Extended Data Fig. 7 and Supplementary Note 5.

NeuroPixelAI provides automated, scalable processing of large-scale calcium imaging datasets. To efficiently handle extensive neuronal recordings, typically containing tens of thousands of neurons across hundreds of thousands of frames, we implemented NeuroPixelAI as an integrated processing platform that combines previously published algorithms with custom engineering and workflow modules (Fig. 1E). The pipeline automates motion correction, image denoising, neuron segmentation, calcium trace extraction, and spike inference, delivering rapid, robust, and fully automated analysis with minimal user intervention.

### Characterization of Meso2P for high-speed, high-resolution mesoscale imaging

Using *Camk2a-Cre*::Ai162D mice, we first assessed single-plane performance. At 7.67 Hz and 2,048 × 2,048 pixels/frame, the system resolved individual layer 2/3 neurons 250 µm below the dura across the entire 6 × 6 mm² FOV (Fig. 2A). High-resolution mode (4,096 × 4,096 pixels, 3.83 Hz) is available when finer morphology is required. Fluorescence intensity and point-spread function were spatially uniform, enabling automated extraction of 13,586 neurons from 2,000 frames in a representative animal (Fig. 2B,C).

**Fig. 2.**
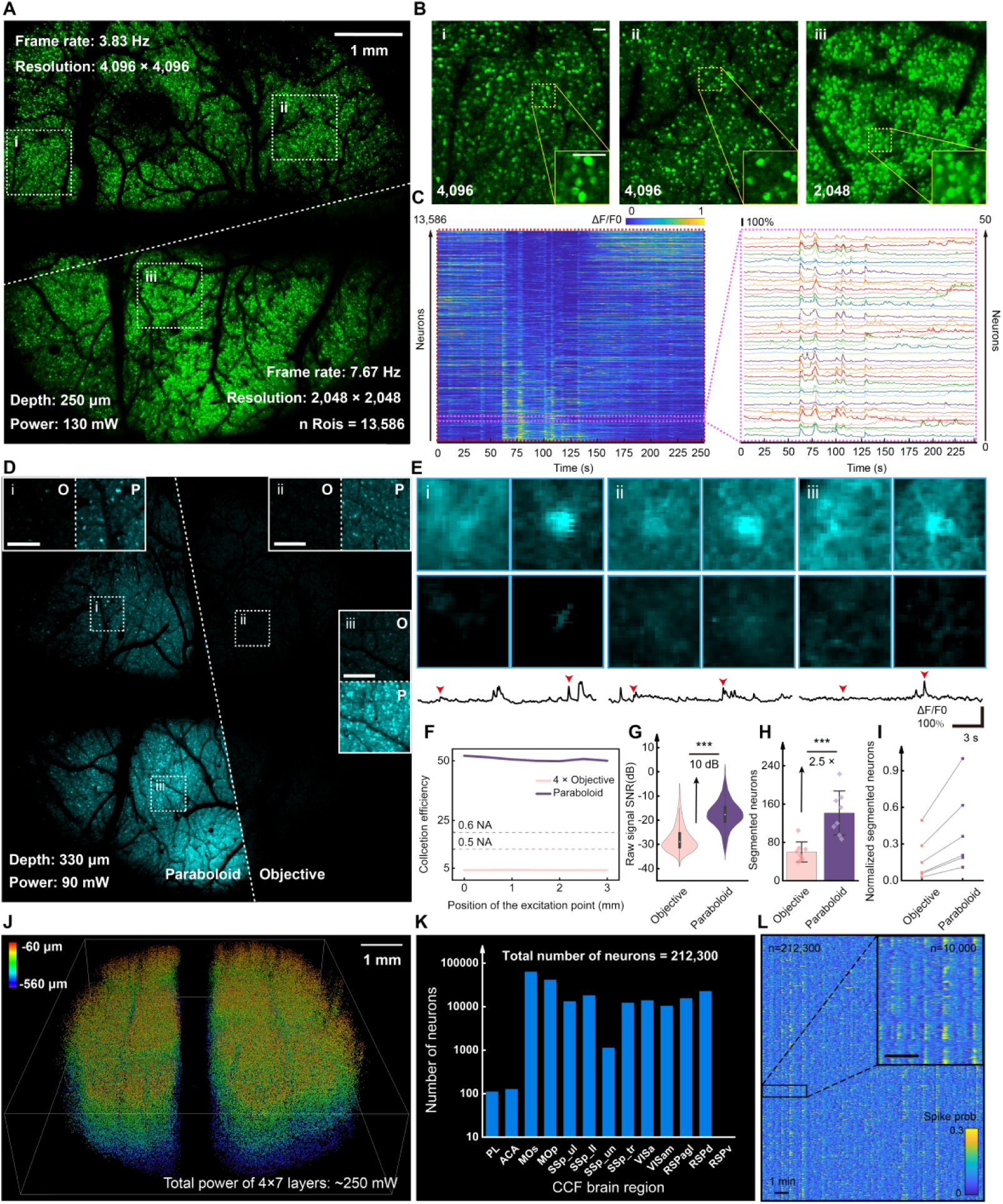
Experimental characterization and performance evaluation of Meso2P. **(A)** In vivo calcium imaging of a transgenic mouse expressing GCaMP6s in excitatory neurons using Meso2P. Top, high-resolution mode (4,096 × 4,096 pixels, 3.83 Hz); Bottom, high-speed mode (2,048 × 2,048 pixels, 7.67 Hz). A total of 13,586 neurons were detected. **(B)** Zoom-in views of the regions marked in (A). **(C)** Heatmaps of calcium activity traces from neurons shown in (A); the zoom-in panel shows traces from 50 example neurons. **(D)** Comparison of image quality between fluorescence signals collected via the parabolic module (left) and direct detection through the objective (right). Three zoom-in views are included. **(E)** Representative neurons and calcium traces obtained using the parabolic module (top) versus the objective (bottom). The red arrow indicates the recording time of the frames. **(F)** Simulated collection efficiency of the paraboloid module and idealized objective-based collection paths with NAs of 0.28, 0.5 and 0.6 under the same upper-hemisphere normalization. **(G)** Violin plot of the signal-to-noise ratios (SNR) from 10,466 neurons imaged with and without the parabolic module. \*\*\**p* < 0.001; Tukey test. **(H)** Quantification of segmented neurons from eight randomly selected 700 μm × 700 μm regions in (D). Data presented as mean ± SD. Significance assessed by Tukey test: \**p* < 0.05,\*\**p* < 0.01,\*\*\**p* < 0.001; n.s., not significant. **(I)** Normalized number of segmented neurons across various segmentation parameters for data with and without parabolic module. **(J)** 3D rendering of 10-minute recording from Meso2P. Neurons are color-coded by depth. **(K)** Histogram of neuron counts extracted across 12 cortical regions. **(L)** Heatmaps of spike probabilities from 212,300 neurons in (J). Zoom-in panel shows spike probabilities of 10,000 neurons. **See also Extended Data** Fig. 3**, Supplementary Video 2.**

Side-collection supported high-quality imaging at this throughput. Zemax ray tracing predicted that the parabolic module collected approximately 51% of the modeled upper-hemisphere emission, corresponding to a solid-angle-equivalent collection NA of 0.87. By comparison, an idealized 0.28-NA objective path collected approximately 4% in the same model, yielding a predicted relative gain of ∼13-fold (Extended Data Fig. 2B,C). For context, under the same idealized upper-hemisphere model, objective-based collection paths with NAs of 0.5 and 0.6^16, 18, 20, 22, 24^ would collect approximately 13.4% and 20.0% of the emission, respectively. The modeled collection fraction of the paraboloid path is therefore approximately 3.8-fold and 2.6-fold higher than these idealized reference values (Fig. 2F). Consistent with this prediction, a separate radiometric measurement yielded a 13.7-fold higher detector signal for the parabolic path than for the 4×/0.28-NA objective path under the tested geometry (Extended Data Fig. 2D,E). Under matched excitation power and acquisition settings, the side-collection configuration was further associated with an approximately 10-dB improvement in the empirically defined trace-based SNR metric and a 2.5-fold increase in neuron segmentation yield (Fig. 2G-I, Extended Data Fig. 2H and Supplementary Video 2).

We next assessed volumetric performance. By capturing 28 axial planes across a 0.5 mm depth at 1.1 Hz, the system achieved high-speed, large-scale volumetric recording (Extended Data Fig. 3A,B). In awake mice, this configuration captured approximately 211,000 neurons spanning over 13 cortical areas (Fig. 2J,K, Extended Data Fig. 3C,D), exceeding the cellular yield reported for one-photon light-field methods imaging similar volumes^17^. Calcium traces maintained high signal-to-noise ratios throughout the volume (Fig. 2L).

Finally, we verified optogenetic compatibility. Under the 4× objective, Meso2P enables spatially distributed photostimulation across multiple brain regions. Under both the 4× and 16× objective configurations, stimulated target sites exhibited stimulation-locked calcium transients, whereas nearby control cells did not show comparable periodic responses (Extended Data Fig. 7, Supplementary Note 5). This proof-of-concept demonstrates spatial co-registration of imaging and perturbation channels within the same optical core.

### NeuroPixelAI enables scalable and automated analysis of mesoscale calcium imaging

NeuroPixelAI is an open-source, GPU-accelerated processing platform that integrates previously published algorithms with custom engineering and workflow modules to convert raw mesoscale imaging data into baseline-corrected activity traces and inferred spike trains via automated motion registration, deep-learning denoising, region-adaptive segmentation, baseline-corrected trace extraction, and spike inference (Fig. 3A,B, Extended Data Fig. 8A and Supplementary Video 3). A streaming phase-correlation implementation with dynamic memory allocation processes a 30-GB file in a single pass on a standard 128-GB workstation, while integrated support for previously published neural-network denoising methods, including DeepCAD-RT^29^ and SRDTrans^30^, allows users to apply these tools within the NeuroPixelAI workflow to suppress shot noise and detector artefacts (Extended Data Fig. 8B,C).

**Fig. 3.**
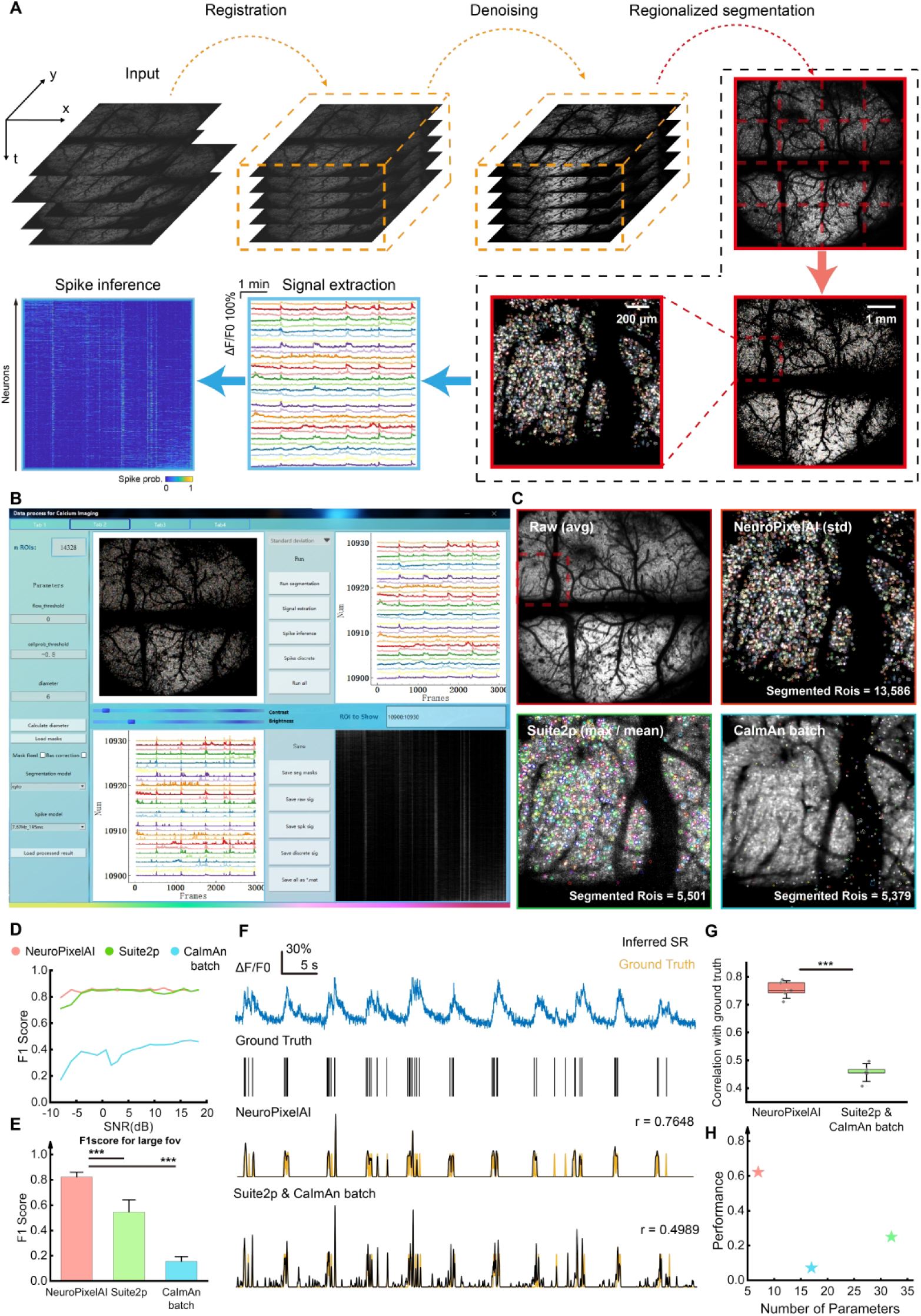
Development and Benchmarking of NeuroPixelAI pipeline. **(A)** Schematic of the NeuroPixelAI processing pipeline, comprising five sequential modules: registration, denoising, regionalized segmentation, signal extraction, and spike inference. **(B)** Representative user interface of NeuroPixelAI. **(C)** Comparative segmentation results on a large FOV dataset. Top left, mean projection of raw fluorescence image. Top right, zoomed-in standard deviation projection with NeuroPixelAI segmentation. Bottom left, Suite2p segmentation (max / mean projection). Bottom right, CaImAn batch segmentation (mean projection). **(D)** F1 score comparison of neuron segmentation performance across the three pipelines on a simulated small FOV (300 μm × 300 μm) dataset generated by NAOMI with Gaussian-Poisson mixed noise. **(E)** F1 score comparison on a large FOV dataset (6 mm × 6 mm). Ten subregions (1.5 mm × 1.5 mm) were randomly sampled for evaluation. Mean ± SD; *\*\*\*p* < 0.001; Tukey test. **(F)** Comparisons of spike inference outputs of NeuroPixelAI, Suite2p and CaImAn batch on the same dataset. **(G)** Correlations between inferred spike trains and electrophysiological ground truth across NeuroPixelAI, Suite2p and CaImAn batch. Mean ± SD; *\*\*\*p* < 0.001; Tukey test. **(H)** NeuroPixelAI achieves higher overall performance (defined as average F1 score × correlation with ground truth) while maintaining a compact set of seven parameters, only 2–4 of which typically require user adjustment under the tested datasets and settings. ***See also* Extended Data** Fig. 8**, Supplementary Video 3.**

Segmentation leverages Cellpose 2.0 within a region-adaptive mask, ensuring reliable soma identification across heterogeneous cortical areas (Extended Data Fig. 8D-F). Cubic-spline correction (Extended Data Fig. 8G) removes baseline drift before Cascade-based spike inference (Extended Data Fig. 8H-J). An iterative, label-free ranking module allows interactive visualization of population activity (Extended Data Fig. 8K).

On our large-FOV datasets and under the tested processing conditions, NeuroPixelAI processes large files directly without manual file splitting. It achieves improved segmentation accuracy (F1-score) on large-FOV data—by ∼ 50% over Suite2p^25^ and ∼ 300% over CaImAn^26^ (Fig. 3C-E), and nearly doubles spike-inference fidelity on GENIE datasets (Fig. 3F,G)^31, 32^. These advantages are achieved with only seven adjustable parameters (typically 2-4 require minor adjustments per dataset), ensuring a lightweight, robust workflow (Fig. 3H). Practically, a one-hour Meso2P dataset (> 200,000 neurons, > 100 GB) can be automatically processed into activity traces and inferred spike trains within a few hours, making large-scale calcium imaging analysis more practical for routine use.

### Cortex-wide neuronal dynamics during whisker stimulation

The mammalian neocortex integrates sensory information through distributed cortical networks, establishing robust yet flexible regional co-fluctuation patterns critical for perception^21, 33–37^. However, it remains poorly understood whether the adaptive reorganization induced by repeated sensory stimuli propagates exclusively within subnetworks or elicits global network-wide reorganization^36–39^. Conventional wide-field approaches capture the spatial extent of such dynamics but are limited by sparse labeling and lack of optical sectioning, restricting depth resolution and the number of neurons sampled^39, 40^. Leveraging Meso2P’s large FOV and optical sectioning, we recorded calcium dynamics from densely labeled layer 2/3 neurons (11,941 ± 3,193 neurons; n = 3 mice) during repeated whisker stimulation (Fig. 4A-C and Supplementary Video 4), enabling us to directly probe functional connectivity changes within large-scale neural networks with high precision.

**Fig. 4.**
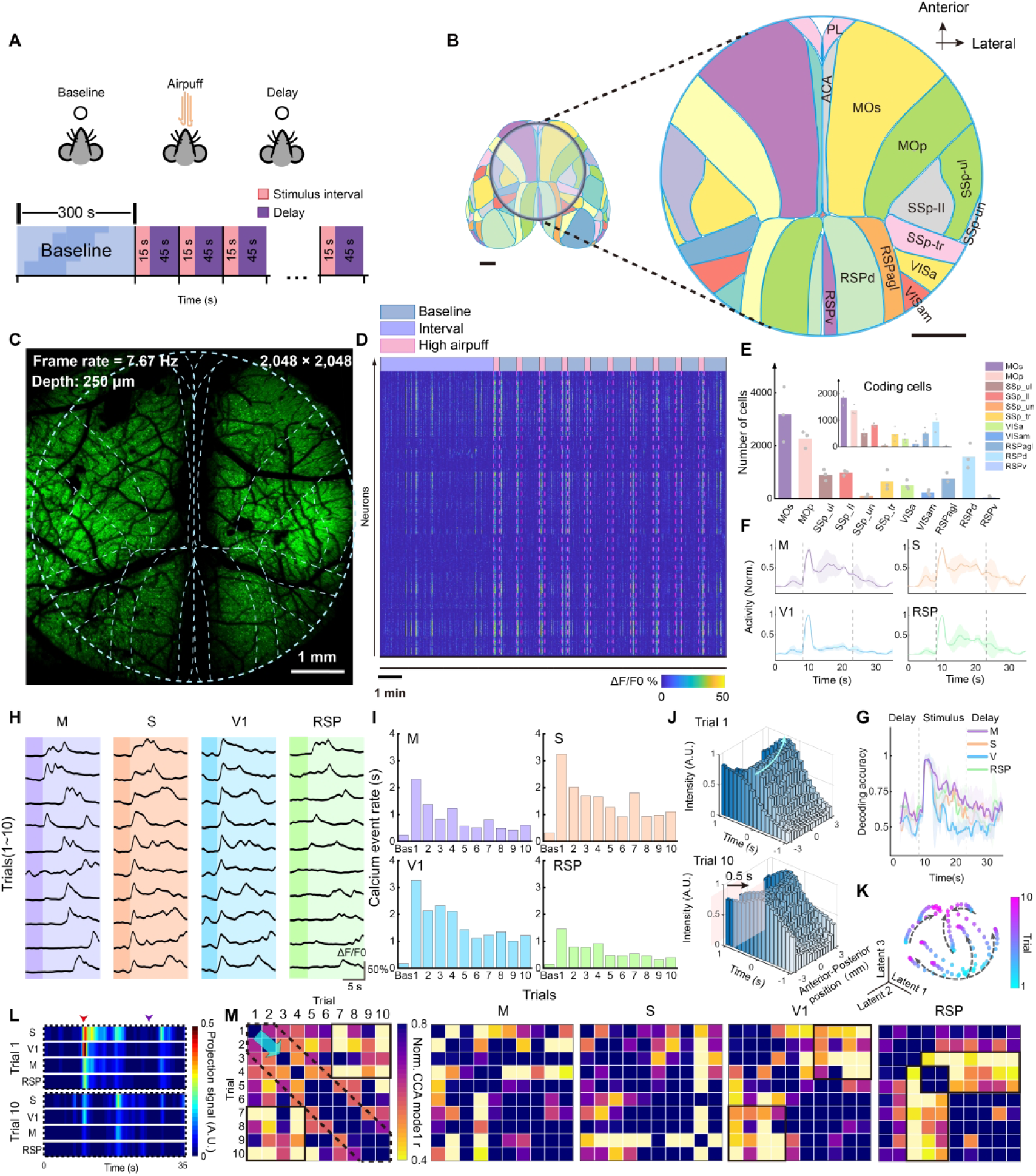
Reorganization of whole-cortex population coding patterns during repetitive whisker stimulation. **(A)** Experimental schematic of the whisker stimulation paradigm. Each session began with a 300 s baseline, followed by ten trials of 15 s air-puff stimulus and 45 s inter-trial intervals. **(B)** Imaging field and anatomical landmarks. Scale bar, 1 mm. Inset: magnified cortical map. PL, prelimbic cortex; ACA, anterior cingulate area. M, motor cortex; S, somatosensory cortex; V1, primary visual cortex; RSP, retrosplenial cortex. The same abbreviations are used in all subsequent figures. **(C)** Standard deviation projection of the calcium imaging video showing > 10,000 neurons, overlaid with cortical area boundaries. **(D)** Heatmaps of calcium signals extracted from neurons in (C). Pink dashed lines indicate stimulus onset in each trial. **(E)** Mean numbers of neurons detected per brain region across mice (11,941 ± 3,193 neurons, mean ± SD; n = 3 mice). Gray dots represent individual animals. Inset: histogram of neurons active across all 10 trials. **(F)** Normalized activity of 4 cortical regions aligned to locomotion onset (vertical dashed line; n = 3 mice). Shaded area represents mean ± SD. **(G)** Mean classification accuracy of stimulus identity across trials using SVM based on regional activity. Dashed lines denote stimulus onset and delay time. Data are presented as mean ± SD (n = 3 mice). **(H)** Trial-by-trial calcium traces from representative neurons across 4 cortical regions. Dark shading, delay period; Light shading, stimulation period. **(I)** Bar plot of average calcium event rate of the 4 cortical regions over baseline and 10 trials. **(J)** Top, bar plots of normalized neuron spike probability at each time point and anterior-posterior bin in trial 1. The blue line indicates the direction of propagation of the peak-signal arrival time. Bottom, same analysis in trial 10. The transparent pink walls mark the peak-signal arrival time in trial 1 and trial 10, respectively. **(K)** 3D CEBRA-Time visualization (on spike probabilities) of neural recording across ten trials in a representative mouse. The gray arrows indicate the direction of change in the trial status. **(L)** The projected activity of each area in the CCA dimensions identified. Top, CCA-projected activity in trial 1. Bottom, CCA-projected activity in trial 10. Signals were projected onto pairwise CCA-derived dimensions across brain regions and averaged (n = 3 mice). A typical change of CCA-projected activity for each area is shown in Extended Data Fig. 9K. **(M)** Each matrix shows normalized correlation coefficients, r, for CCA mode between two of ten trials. A large matrix element value indicates that the source cofluctuated with the target using a similar activity mode; small values imply distinct cofluctuation modes. Results are shown for the largest CCA modes for each source-target pair, averaged over three mice. Solid-outlined boxes highlight areas with lower relevance. Dashed-outlined box indicates stronger correlation between adjacent trials. ***See also* Extended Data** Fig. 9 **and Supplementary Video 4.**

Single air-puff stimuli triggered widespread, cortex-wide activation across multiple areas (Fig. 4D), where neurons displayed diverse stimulus tuning profiles (Fig. 4E, Extended Data Fig. 9A,B) but with significant co-activation across different areas (Fig. 4F). Population decoding via support-vector machines revealed stimulus presence with > 90% accuracy for 1.5-2 s post-stimulus (Fig. 4G), and accuracy scaled positively with neuron count (Extended Data Fig. 9C,D). Consistently, S1 and V1 contributed more significantly and stably to the overall response pattern (Fig. 4H,I), and the stimulus-evoked information was predominantly driven by inter-regional shared fluctuations between S1 and V1 (Extended Data Fig. 9K).

Repeated stimuli induced adaptive reorganization of cortical population activity, decreasing calcium event rates and peak spike probabilities across all brain regions, stabilizing after several trials (Fig. 4I, Extended Data Fig. 9E-G). Posterior cortical areas (AP ≈ –3 to –2.5 mm) exhibited advanced response onsets (∼ 0.5 s) in later trials, indicating a temporal reorganization (Fig. 4J, Extended Data Fig. 9H,I). CEBRA embeddings of neuronal trajectories showed directional shifts converging onto stable attractors with repeated stimulation (Fig. 4K).

Canonical correlation analysis (CCA) revealed region-specific reorganization of neuronal population activity (Extended Data Fig. 9J). Although stimulus information was primarily encoded by locally shared fluctuation patterns between S1 and V1 (Fig. 4L, Extended Data Fig. 9K), global coordination across cortical regions emerged during adaptation to repeated stimulation. This global coordination exhibited regionally heterogeneous temporal engagement: Motor cortex (M1), S1, and V1 maintained sustained involvement, while the retrosplenial cortex (RSP) underwent rapid reconfiguration during mid-adaptation (Fig. 4M). Critically, analysis of neuronal loading weights across regions demonstrated that adaptive reorganization to sensory stimuli involved not merely changes in network response magnitude, but also restructuring of information transfer architecture within the network. This restructuring was evidenced by directional dynamic changes in unit-specific weights, which were no longer driven by specific local subnetworks but instead reflected global coordinated shifts (Extended Data Fig. 9L,M).

Thus, leveraging Meso2P technology, we achieved cortex-wide cellular-resolution recordings at high temporal resolution in densely labeled mice, revealing network-level changes potentially underlying robust perceptual adaptation. Its large FOV design enables high-fidelity, high-throughput spatially extended characterization of functionally distinct cortical regions at mesoscopic scales, facilitating neural network architecture analyses.

### Simultaneous dual-layer imaging reveals laminar dynamics during anesthesia

Conscious processing is thought to rely on activity within the cerebral cortex^41–45^, whereas general anesthesia induces large-scale cortical network dynamics characterized by widespread neuronal synchrony, particularly involving layers 2/3 and 5. Notably, synchrony changes in layer 5 neurons are temporally aligned with the loss and recovery of consciousness^41–45^. Capturing this phenomenon across multiple cortical layers simultaneously, at cellular resolution and mesoscale coverage, remains challenging with existing approaches.

Using Meso2P, we performed simultaneous mesoscale imaging of layers 2/3 and 5 in head-fixed, isoflurane-anesthetized mice (Supplementary Video 5), with imaging planes at ∼ 300 μm and ∼ 560 μm below the cortical surface covering prefrontal and posterior cortices (Fig. 5A-B). Throughout both induction and recovery phases, the system reliably captured large populations of active neurons from both layers with high signal fidelity (Fig. 5C-D). Notably, both layers exhibited prominent large-scale synchrony across millimeter-scale regions during anesthetic induction (Fig. 5E)—consistent with previous studies^45–48^—with this effect being particularly pronounced in layer 5 (Fig. 5F-H). Unexpectedly, leveraging Meso2P’s cross-layer, high-speed and large-scale imaging capability, we observed asynchronous information flow within global synchronization events: individual “synchronous” events contained distinct spatiotemporal origins and propagation pathways, with near-complete global synchrony emerging only during late anesthesia. Granger causality analysis revealed a predominant posterior-to-anterior directional bias during induction (Fig. 5I). To validate this finding, we analyzed the spatiotemporal sequence of individual synchronized events by aligning normalized calcium activity across 16 contiguous cortical windows within a ± 2 s window centered on peak activation (Fig. 5J, Extended Data Fig. 10A,B). Although activity appeared globally synchronized, consistent temporal offsets revealed heterogeneity in initiation sites and propagation trajectories. As anesthesia deepened, this variability progressively diminished, with Layer 5 neurons achieving near-complete synchronization exclusively during late-stage events (Fig. 5J, Extended Data Fig. 10B,C and Supplementary Video 6). Quantitative analysis of spike onset distributions confirmed posterior-to-anterior propagation as the dominant direction (Fig. 5K).

**Fig. 5.**
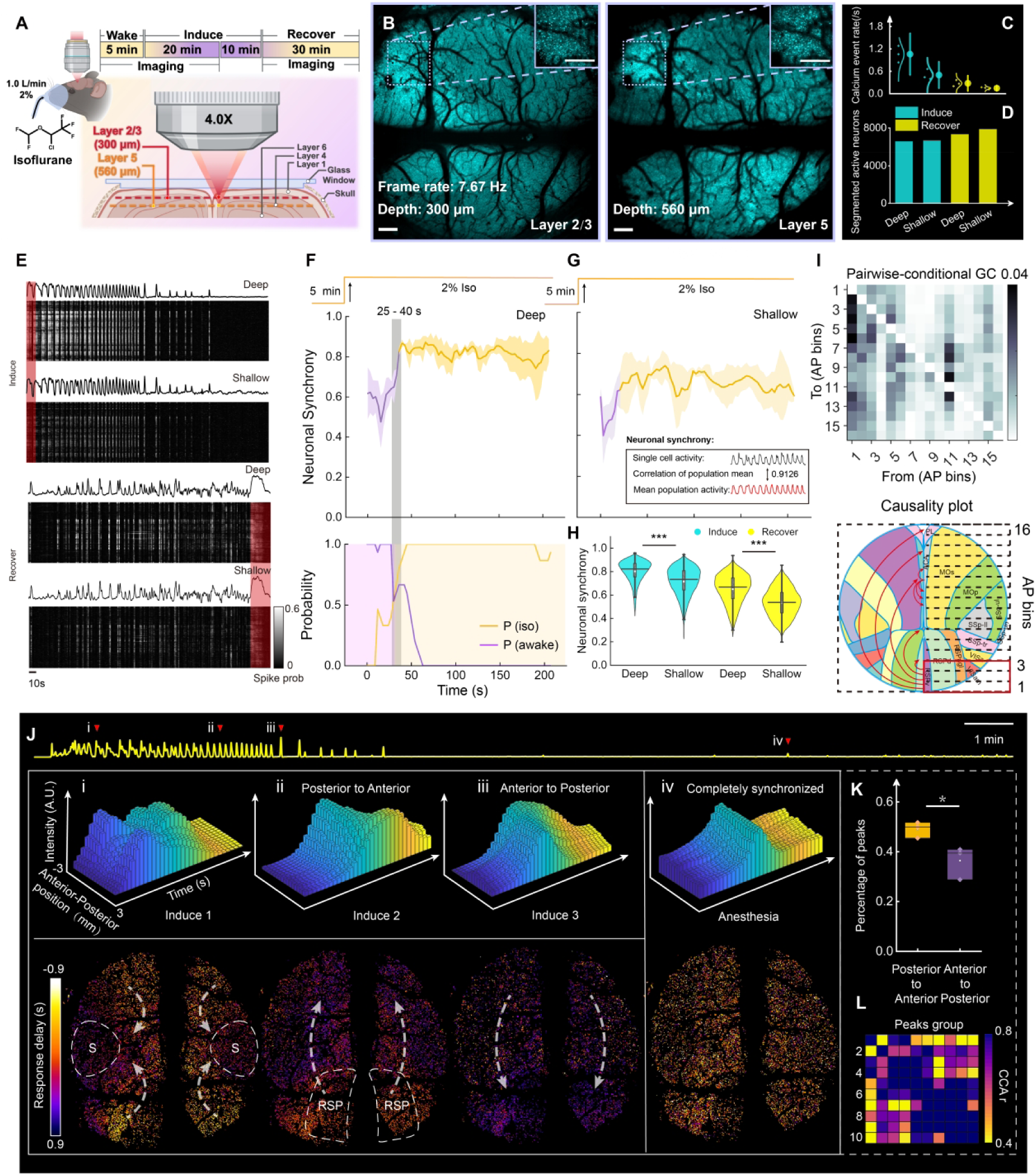
Cross-laminar recording reveals dynamic signal propagation within and across cortical layers during anesthesia induction. **(A)** Experimental schematic showing simultaneous two-plane calcium imaging during general anesthesia, with acquisition depths at 300 μm (cortical layer 2/3) and 560 μm (layer 5). **(B)** Representative two-plane imaging examples acquired simultaneously at dual depths, with zoom-in views showing neurons in the same cortical area at both depths. **(C)** Mean calcium event rates of all tracked neurons across different periods and depths, calculated from the initial 3-minute window per period (n = 3 mice). Error bars represent 95% confidence intervals. **(D)** Bar plot of the average number of segmented active neurons across different periods and depths. **(E)** Spike probabilities in a representative mouse. Black trace, averaged signals. Red bands, state transition periods. **(F-G)** Neuronal synchrony of all tracked neurons. Top, anesthesia protocol. Middle, neuronal synchrony in 15-s rolling window. Bottom, probability of mouse being in various states. Purple shading indicates P(awake) > P(iso); yellow shading indicates P(iso) > P(awake). Gray vertical bar marks crossing of these states. Data shown as mean ± SD across n = 3 mice. **(H)** Violin plot of neuronal synchrony across different periods and depths (n = 21,387 active cells from three mice). *\*\*\*p* < 0.001; Tukey test. **(I)** Top, Granger causality matrix across 16 cortical bins during anesthesia. First column shows causality from the most posterior bin to other bins. Causality with \**p* < 0.05 (Granger F test) was shown after correction for multiple comparisons to control false discovery rate. Bottom, summary map of causality between cortical bins. Red arrows indicate inferred causality direction. **(J)** Schematic of synchronized signal propagation direction in a representative mouse. Top, mean spike probabilities during anesthesia induction. Red arrows indicate synchronized spikes expanded in the middle and bottom panels. Middle, bar plots of normalized neuronal spike probability across anterior-posterior positions, showing peak timing sequences in four synchronized events. Bottom, single-neuron-resolution response delay map. Neurons are color-coded by their peak time relative to population activity. (Yellow: early-peaking source regions; purple: late-peaking target regions). Full replication in mice shown in **Extended Data** Fig. 10C (n = 3 mice). **(K)** Boxplot of signal propagation direction. For each detected synchronized event, peak timing differences were computed between anterior (8 bins) and posterior (8 bins) regions, then averaged to determine information flow (n = 3 mice), *\*p* < 0.05, Tukey test. **(L)** Heatmap of normalized CCA mode correlation matrices between consecutive synchronized event clusters, revealing gradual reorganization of neuronal ensemble activity patterns during anesthesia. ***See also* Extended Data** Fig. 10**, Supplementary Videos 5 and 6.**

CCA analysis further revealed progressive divergence in global synchrony patterns and substantial reconfiguration of neuronal loading weights across trials (Fig. 5L; Extended Data Fig. 10D).

Dual-layer imaging also allowed direct interrogation of interlaminar interactions. Activity predominantly propagated in a bottom-up manner from layer 5 to layer 2/3 (\*\**p* < 0.01), with occasional bidirectional interactions (Extended Data Fig. 10E,F).

In summary, these findings show that anesthesia-induced cortical synchrony emerges primarily in deep layers and propagates in a posterior-to-anterior and bottom-up fashion, accompanied by dynamic reconfiguration of interlaminar interactions—significantly departing from the minimal delay synchronization proposed in prior studies^41–45^. By combining high-speed, large-scale and multilayer imaging, Meso2P supports investigation of information propagation across macroscopic cortical networks and laminar organization during anesthesia and other altered states of consciousness.

### Dynamic monitoring of MNP distribution and vascular occlusion *in vivo*

Micro- and nanoplastics (MNPs) have rapidly become widespread environmental contaminants, increasingly detected in human tissues including the brain^49^.

Conventional analytical methods (e.g., Py-GC-MS and Raman imaging) quantify MNPs but lack the temporal resolution required for dynamic biological studies^50^. Standard 2P microscopy, constrained by its small imaging field, also struggles to capture real-time three-dimensional MNP distribution and cerebrovascular interactions.

We leveraged Meso2P’s large-FOV volumetric imaging to first delineate cortical vasculature of C57BL/6J mice recovered from surgery with vascular dyes (Extended Data Fig. 11A), then dynamically tracked fluorescent 5-μm microplastics (MPs) and 200-nm nanoplastics (NPs), revealing their distinct spatiotemporal accumulation patterns and cerebrovascular consequences (Fig. 6A,B).

**Fig. 6.**
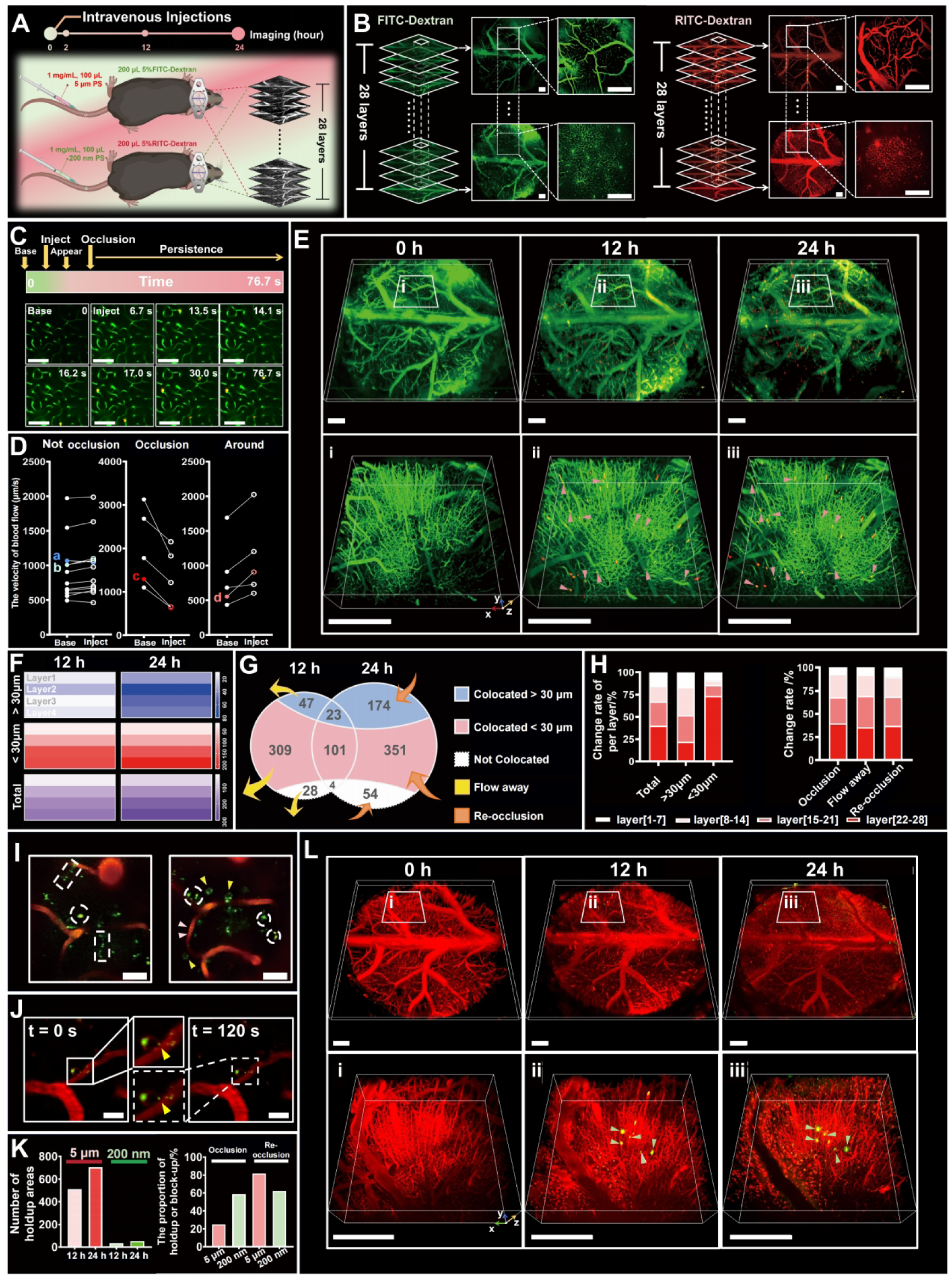
Dynamic monitoring of MNP distribution and vascular occlusion in the living brain. **(A)** 28-layer MNP imaging in head-fixed mice using Meso2P. **(B)** Vascular resolution in 28-layer Meso2P imaging: FITC-Dextran (green) and RITC-Dextran (red) labeled vessels. **(C)** Dynamic appearance and persistence of a vascular occlusion after injection of 5-µm MPs. Vessels (green), MPs (red), co-localization (yellow). **(D)** Comparison of blood flow velocity in MP-treated vessels; data sources a-d shown in **Extended Data** Fig. 11B. **(E)** MP distribution at different time points in volumes of 6 × 6 × 0.45 mm³ (top) and 1.6 × 1.6 × 0.45 mm³ (bottom): vessels (green), MPs (red), co-localization (yellow), co-localized MPs observed at both time points (pink arrows). **(F)** Depth distribution of MPs in vessels of different diameters across cortical layers 1–4 (top to bottom) at two time points. **(G)** Spatiotemporal distribution of MPs in vessels of different diameters at two time points. **(H)** Turnover rates of MPs across layers, and proportions of occlusion, clearance, and re-occlusion between layers. **(I)** Distribution pattern of 200-nm NPs in the brain after injection: NPs adhering to vascular walls (pink arrows), within NP-laden cells (yellow arrows), extravascular distribution (white dashed circles), and occluding vessels (white dashed rectangles); vessels (red), NPs (green). **(J)** NP uptake by NP-laden cells: pre-uptake (white solid line) and post-uptake (white dashed line) position comparison (yellow arrows). **(K)** Comparison of vascular occlusion between MPs and NPs. **(L)** NP distribution at different time points in volumes of 6 × 6 × 0.45 mm³ (top) and 1.6 × 1.6 × 0.45 mm³ (bottom): vessels (red), NPs (green), co-localization (yellow), co-localized NPs at both time points (green arrows). *Scale bars*: 500 µm (B, E, L); 100 µm (C); 20 µm (I, J). ***See also* Extended Data** Fig. 11 **and Supplementary Videos 7-11.**

MPs entered cortical vessels rapidly (∼ 13 s post-injection), causing immediate vascular occlusions (Fig. 6C, Supplementary Video 7). Occluded vessels exhibited reduced blood flow, compensated by increased flow in adjacent vessels (Fig. 6D; Extended Data Fig. 11B). Compared to the 12-hour time point, partial vascular occlusions were resolved by 24 hours, while some persisted for at least 12 hours (Fig. 6E). MPs preferentially accumulated in smaller vessels (< 30 µm diameter), particularly within deeper cortical layers, and either persisted or re-occluded after partial clearance (Fig. 6F-H, Extended Data Fig. 11C, Supplementary Video 8).

In contrast, smaller-sized NPs diffused broadly upon vascular entry, forming diverse aggregates (granular, aggregated, and cellular) within vessels and surrounding tissues (Fig. 6I, Extended Data Fig. 11D, Supplementary Video 7). NPs adhered to vascular walls, occasionally extravasated, and progressively accumulated within putative phagocytes with time-dependent increase (“NP-laden cells”, Fig. 6J, Extended Data Fig. 11E, Supplementary Video 8). NP-induced vascular occlusions, though fewer, were notably persistent (∼ 60% unresolved; Fig. 6K,L). Vessel wall adhesion promoted recurrent clustering, stabilizing occlusions despite NPs’ small size (Extended Data Fig. 11F, Supplementary Video 9).

Thus, Meso2P offers a powerful platform for dynamic, cellular-resolution analysis of complex *in vivo* processes. Its volumetric imaging capacity captures three-dimensional particle distributions and biological responses in real time, enabling new investigations into a wide range of questions that demand large-scale, three-dimensional coverage.

## Discussion

Meso2P addresses a long-standing collection-efficiency bottleneck in large-FOV 2P microscopy by physically separating excitation and fluorescence collection. In conventional single-objective mesoscopes, increasing the FOV usually requires a long focal length and therefore reduces NA, which limits fluorescence collection efficiency. Meso2P builds on the principle of supplementary fluorescence collection^28^ and relieves this limitation by using a 4× excitation objective to provide lateral cellular resolution across a 6 × 6 mm² imaging window, while a compact paraboloid collector recovers fluorescence over a large solid angle, corresponding to an equivalent collection NA of 0.87. This design does not increase the excitation NA. Instead, it exploits a regime relevant to mesoscale calcium imaging: the excitation PSF and sampling are sufficient for soma-level lateral resolution, whereas the detected fluorescence signal is strongly influenced by collection efficiency. In the thick-sample limit, multiphoton excitation theory indicates that increasing focusing NA does not necessarily increase the total generated fluorescence when other conditions are fixed, whereas the detected signal remains strongly dependent on fluorescence collection efficiency^5, 12^. Thus, Meso2P converts high-efficiency fluorescence collection from a constraint imposed by objective geometry into a modular optical design choice for cortex-wide imaging.

The volumetric implementation pushes single-beam point scanning toward its physical limits. Spatiotemporal multiplexing interleaves four 80-MHz beamlets with a nominal temporal spacing of 3.125 ns, which is close to the reported fluorescence lifetime of GCaMP6s (2.6–2.8 ns)^51^. Piezo-driven axial stepping distributes excitation pulses across a 520-µm depth range at 1.1 Hz. The resulting 6 × 6 × 0.5 mm³ imaging volume provides a practical balance among sampling density, tissue heating, and indicator kinetics under the tested conditions. Achieving greater imaging speed would likely necessitate parallel excitation approaches, potentially compromising resolution and signal fidelity in scattering tissues. Crucially, Meso2P’s modular design—comprising lateral collection, volumetric imaging, and multi-region photostimulation modules—allows users to flexibly adapt the instrument according to experimental requirements and to incorporate additional functionalities as imaging technologies evolve.

Handling the data rate of cortex-wide imaging motivated the development of NeuroPixelAI. Under the tested datasets and processing conditions, the pipeline processes full-session files on a single workstation and achieves higher segmentation accuracy and spike-inference fidelity than the compared alternatives. Its open-source code base and modular design invite adoption of new models as they appear and lower the barrier for laboratories that lack extensive computational expertise.

Although we primarily demonstrated Meso2P’s capabilities through neuroscience applications, its optical core is versatile and equally suited to imaging dynamic biological processes such as immune-cell trafficking, tumor-stromal interactions, or embryonic morphogenesis, wherever large-scale coverage with single-cell resolution is required.

Several limitations remain. Although Meso2P achieves lateral single-cell resolution across a large FOV, its 21-24 µm axial point-spread function approaches the diameter of a cortical soma, so applications that demand sub-laminar separation will still benefit from high-NA objectives. The lateral paraboloid collector occupies the space above the cranial window and therefore imposes practical constraints on experimental access. In the current configuration, conventional large-field visual stimulation is not readily compatible with the light-sealed collector geometry, and integration with head-mounted devices is limited. Some tactile stimulation paradigms also require additional mechanical optimization. Nevertheless, the front and lateral sides of the mouse remain accessible, allowing the air-puff, limited whisker-stimulation, anesthesia-related and face/pupil-monitoring experiments demonstrated in this study. The four optical arms exhibit arm-dependent, incompletely compensated group delay dispersion. In future designs, additional dispersive elements could be inserted into the shorter-GDD arms to better match the GDD among the four arms, enabling more uniform compensation when combined with a laser source or external compressor with a larger pre-chirp range. The 3.125-ns inter-beamlet temporal delay represents a compromise between crosstalk and multiplexing depth. Crosstalk could be further reduced by using detectors with faster response times or by reducing the number of multiplexed planes, thereby increasing the inter-beamlet delay to 4-6 ns. NeuroPixelAI analyses each imaging plane independently; a fully three-dimensional model could further improve source separation in densely labeled tissue. Finally, while we confirmed accurate co-registration of imaging and holographic stimulation, we did not explore closed-loop perturbations, leaving ample scope for future circuit manipulation experiments.

## Conclusion

Collectively, Meso2P demonstrates that decoupling fluorescence detection from excitation is a practical strategy for cellular-resolution imaging across a mesoscale FOV. Its modular hardware–software architecture provides a reproducible and flexible framework for next-generation instruments aimed at high-throughput imaging, perturbation and analysis of biological activity across extensive cortical volumes.

## Methodological Details

### Animals

All mouse (*Mus musculus*) experiments were approved by the Institutional Animal Care and Use Committee of the Department of Laboratory Animal Science, Fudan University, Shanghai, China (ethical approval no. 2022JSITBR-013). Mice were housed in a controlled environment (∼ 20 °C, 45% relative humidity) under a 12 h light/dark cycle (lights on 08:00, off 20:00). Unless specified, both male and female mice transgenically expressing GCaMP6s in excitatory neurons (*Camk2a-Cre*, Cyagen Biosciences #C001015 × Ai162D, Jackson Labs #031562; ROSA-LSL-GCaMP6s) underwent cranial window implantation at 8-10 weeks. Briefly, under 1.5% isoflurane (RWD Life Science #R510-22-10) anesthesia (mixed with oxygen), craniotomy was drilled (7 mm). The skull flap was removed while preserving meningeal integrity. After debris clearance, a cranial window (6.5-mm coverslip bonded to a glass ring: 7.5-mm outer diameter, 6-mm inner diameter) and a metal headpiece fixed with dental cement/cyanoacrylate glue were implanted. Mice with major intraoperative hemorrhage were excluded. After 2-week recovery (daily Dexamethasone, Beyotime Biotechnology #ST1254g, 1 mg/kg, i.p. and Ceftriaxone sodium, Shanghai Macklin Biochemical Technology #C832409, 50 mg/kg, i.p., once daily for 3 days), mice were adapted to head fixation for 2 weeks before imaging. Unless specified, head-fixed imaging used a parabolic lateral collection module and custom 4× objective. At the end of the experimental procedures, unless specified, experimental animals were euthanized by CO₂ asphyxiation followed by cervical dislocation. Neuronal imaging signal extraction was performed using NeuroPixelAI.

## System Configuration

### Laser Source

The laser system consisted of two components: an imaging laser source and a photostimulation laser source. For imaging, we employed the Chameleon Discovery NX laser system (Coherent, Inc.), which features a tunable wavelength range from 660 nm to 1,320 nm, a pulse duration of 100 fs, and a repetition rate of 80 MHz. At a wavelength of 920 nm, the system delivers an average output power of 3,200 mW. We first utilized the built-in dispersion compensation module of the laser to introduce an anomalous group delay dispersion (GDD) of -17,358 fs², which partially compensated the material dispersion accumulated in the system. The residual total GDD of the first multiplexed beam path, including the main optical path and the first arm of the spatiotemporal multiplexing module, was approximately 8,082 fs² (Supplementary Note 6 and Supplementary Tables 4 and 5), which broadened the pulse duration at the sample from the ideal 100 fs to 245 fs. As a result, the excitation power required to generate the same fluorescence level was 1.56 times that needed under ideal conditions. In addition, after the spatiotemporal multiplexing module, the four beamlets carried different residual GDD values (Supplementary Table 6); system performance could be further improved by incorporating additional dispersion compensation. We then adjusted the laser output power using a rotatable half-wave plate (AHWP05M-980, Thorlabs) combined with a polarizing beam splitter (PBS255, Thorlabs), equipped with a precision rotation mount (PRM1) and a voice coil motor controller (KVC101, Thorlabs). For photostimulation, we used a Monaco 1035 laser system (Coherent).

### Spatiotemporal Multiplexing Module

To enable four-channel spatiotemporal multiplexing, we constructed a polarization-based system that interleaves four 80-MHz beamlets with nominal temporal offsets of 3.125 ns (Extended Data Fig. 3A,B). The system initially divides the incident light into four beams using a combination of three polarizing beam splitters (PBS623, LBTEK) and half-wave plates (HWP25-905A-M, LBTEK). Each half-wave plate is mounted on a rotatable adjustment frame (CRM-1AS) to control power allocation among the optical paths. Each optical path was then assigned a distinct time delay, resulting in the sequential emission of the four light beams at 3.125 ns intervals. Furthermore, the system incorporated a pair of cemented lenses (AC250-150-B-ML and AC250-300-B-ML, Thorlabs) arranged as a magnification telescope. This arrangement enabled beam enlargement while allowing focal position adjustment using a precision translation stage (A-LSQ-E, Zaber), thereby creating four adjustable focal points on the imaging plane. Finally, the optical paths were combined using two beam splitters (BS1255-B, LBTEK) and a polarizing beam splitter, producing four interleaved beamlets with nominal temporal intervals of 3.125 ns and adjustable spatial separation between their focal planes. Temporal crosstalk between adjacent beamlet channels at this spacing was quantified and reduced by offline demixing (Extended Data Fig. 5 and Supplementary Note 4). The power of each beam within the composite light and the spacing between focal points could be adjusted via the corresponding half-wave plates and by moving the lenses on the translation stage, respectively, to meet varying imaging requirements.

### Excitation Optical Pathway

The excitation optical pathway of Meso2P is shown in Fig. 1A and Extended Data Fig. 1B,D. The optical path begins with a Y-axis scanning galvanometer mirror (6215H, Cambridge Technology) at the system forefront. The beam then passes through a telescope comprising two telecentric flat field scanning lenses (LSM54-850, Thorlabs) before being incident on a resonant mirror (CRS8K, Cambridge Technology) for X-axis scanning. This separated-axis design prevents sampling depth changes from affecting the XY field of view (FOV). After the scanning process, the beam undergoes aperture magnification through a magnification telescope composed of a telecentric flat field scanning lens (LSM54-850, Thorlabs) and two doublet achromatic lenses (AC508-300-B), followed by a custom aberration compensation module before entering the rear aperture of the objective lens.

### Aberration Compensation Module

In Meso2P, most optical modules employ commercial low-aberration components, but cumulative aberrations from optical mismatches still significantly degrade image quality. To correct multiple types of aberrations present in the system and enhance the fluorescence excitation efficiency, we designed a symmetric cemented lens pair with large air gaps, inspired by the Petzval lens structure. This pair of lenses, after design optimization, features an extended focal length as given by:

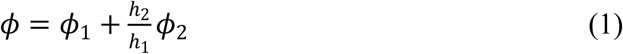

Here, *ϕ*_1_ and *ϕ*_2_ represent the focal powers of the front and rear lens groups, respectively, and *ϕ* denotes the total focal power of the module. This design allows for significant optimization of spherical aberration, field curvature, and astigmatism without affecting the original focal position of the system. The design primarily focuses on optimizing the flatness of the imaging field quality, ensuring that the overall composite aberration RMS across the imaging field remains within 0.1 ± 0.01 λ at the operational wavelength of 920 nm ± 18 nm.

### Parabolic lateral collection module

The parabolic lateral collection module consists of six components: a parabolic reflector module, a first focusing lens (LA1145-A, *f* = 75 mm, Thorlabs), a filter cube module (including custom-sized low-pass filter ET720sp-2p8, high-pass filter T556lpxs-UF3, and band-pass filters for red and green wavelengths ET605/70m and CT512/60bp, Chroma Technology), a second focusing lens, a third focusing lens (48-244-INK, *f* = 50 mm, Edmund Optics), and a photomultiplier module (H13543-300, HAMAMATSU) (Fig. 1B and Extended Data Fig. 2A).

Due to the constraints imposed by the aperture of optical components, the imaging field of view (FOV), and the objective geometry, the parabolic reflector module was designed jointly with the excitation objective. The objective working-distance constraint *WD*_obj_, the parabolic-geometry distance *p*, and the distance *L* behind the focus are subject to the following limitations:

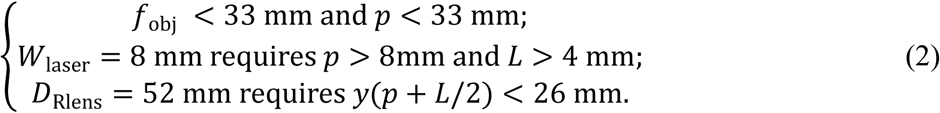

Furthermore, to ensure that the fluorescence collection efficiency is nearly uniform across different points on the imaging plane, the distance before reflection along path 1 should be relatively long, necessitating a maximally large *p*. Additionally, to enable the collection module to gather fluorescence at larger angles, the angle θ along path 2 should be relatively small, which requires to be maximally large while keeping *p* as small as possible. A grid search method was employed to balance these constraints, ultimately selecting the module parameters *p* = 20 and *L* = 8.

Given the absence of commercially available 4× long-working-distance objectives suitable for large-field multiphoton fluorescence imaging with a working distance greater than 30 mm, a custom objective lens was designed and fabricated to accommodate the parabolic lateral collection module (Fig. 1B and Extended Data Fig. 2A). This objective uses a five-group, seven-element design, equipped with an M32 × 0.75 thread, and is mounted on a piezo objective scanner (PFM450E, Thorlabs), allowing for independent axial scanning. The objective has a focal length *f* = 45 mm, a numerical aperture NA = 0.28, and a working distance of 31 mm. Targeted optimizations were performed to minimize spherical aberration, astigmatism, and meridional field curvature, ensuring that the composite wavefront aberration RMS across the entire field at the operational wavelength of 920 nm remains below 0.02 λ. Using Zemax-based optical system simulations, this lens approaches the performance of an ideal objective. Notably, due to the customizable structure of the self-designed objective, adjustments were made during simulation to further enhance image quality, aligning the lens optimally with the system.

### Multiplexed Acquisition of Quadruple Electrical Signals

Due to the frequency limitations of the data acquisition card (vDAQ, MBF Bioscience), the maximum signal acquisition rate of the system is constrained to 80 MHz. To acquire the four temporally interleaved signal channels using an 80-MHz acquisition card, a strategy involving the multiplexed acquisition of quadruple electrical signals was implemented. During the imaging process, the fluorescence signal is first converted into an electrical signal by a photomultiplier tube (PMT, H13543-300, Hamamatsu), which is then amplified by an amplifier. The output from the amplifier (C9663, Hamamatsu) is fed into a four-way power splitter (PD-0/6-4S, TD-MICROWAVE), which divides the signal into four pathways. To achieve temporal displacement among the multiple acquisition points, different cable lengths are used based on the rate of electrical transmission through the conductor. This arrangement allows the four separate channels of the acquisition card to capture the four temporally interleaved signal channels sequentially. The difference in cable lengths can be estimated using the following equations:

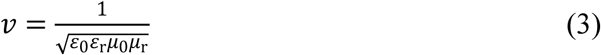

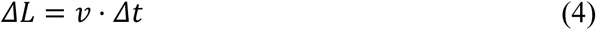

where *ε*_0_ is the permittivity of free space, 8.89 × 10^-12^ F/m, *ε*_r_ is the relative permittivity of the material, *μ*_0_ is the permeability of free space, 4π × 10^-7^ H/m, *μ*_r_ represents the material’s relative permeability, and Δ*t* is 3.125 ns.

### Quantification of parabolic collector collection efficiency enhancement

To quantitatively evaluate the enhancement in fluorescence collection efficiency provided by the parabolic collector module, we first performed ray-tracing simulations using Zemax (Extended Data Fig. 2B). The fluorescence excitation spot was modeled as an isotropic point source with a power of 1 W, and a cone angle of 90° was applied to restrict the emission to the upper hemisphere. The collection path of the parabolic collector module was implemented according to the experimentally built configuration, whereas the objective collection path was represented by a detector path with an aperture defining an NA of 0.28. The radial position of the point source was varied from 0 to 3 mm in steps of 0.5 mm. The simulation results show that the collection efficiency of the parabolic collector module remains stable at 51%, compared with 4% for the NA = 0.28 objective, corresponding to an approximately 13-fold improvement (Extended Data Fig. 2C). For contextual comparison with commonly used mesoscope objectives, idealized objective-based collection paths with NAs of 0.5 and 0.6 were evaluated using the same upper-hemisphere normalization, yielding collection fractions of approximately 13.4% and 20.0%, respectively. To express this collection efficiency in a standardized manner, we used the equivalent relationship between collection efficiency achievable within the upper 2π solid angle and numerical aperture to derive an equivalent collection NA of 0.87 for the parabolic collector module:

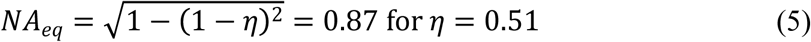

To support this theoretical result experimentally, we next performed relative radiometric measurements for the objective and the parabolic collector module separately (Extended Data Fig. 2D,E). A low-power 3535 LED emitting at 520 nm (maximum current 60 mA, nominal operating voltage 3.0 V) served as the point source, and a microscope CCD camera was used as the detector. An ND 2-400 neutral-density filter was placed in front of the camera to prevent saturation. Both measurements used the same camera, neutral-density filter, acquisition settings and detector-plane geometry. We recorded the absolute integrated pixel values for LED driving currents ranging from 4 mA to 22 mA and then calculated the enhancement factor of the parabolic collector module relative to the objective, yielding a value of approximately 13.7-fold. Considering that, in the Zemax simulation, the objective collection efficiency was evaluated with an idealized equivalent-NA detector, this experimentally measured enhancement is consistent with the 13-fold improvement obtained from the simulation.

### Assessment of neuronal signal quality and segmentation improvement with the parabolic collector module

To quantitatively assess the improvement in neuronal signal quality afforded by the parabolic collector module, we compared the signal-to-noise ratio (SNR) and the number of segmentable neurons between the two acquisition configurations on the same set of samples (Fig. 2G-I). For SNR evaluation, 10,466 neurons distributed across a 6 × 6 mm² field of view were selected from the same sample. Using the denoised neuronal signal (*S*_denoised_) as an empirical reference, we calculated the SNR of the corresponding raw signal (*S*_raw_) acquired with the parabolic collector and with a 4×/0.28-NA objective-based collection condition. SNR was defined as:

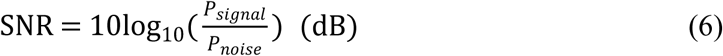

where *P*_signal_=|*S*_denoised_|^2^ and *P*_noise_=|*S*_raw_−*S*_denoised_|^2^. Because this is a denoised-reference, trace-based metric, it is reported as an empirical signal-quality measure and is not converted into an independent estimate of photon number. Under the tested conditions, the parabolic collector module increased neuronal SNR by approximately 10 dB relative to the 4×/0.28-NA objective path (Fig. 2G; \*\**p* < 0.001, Tukey’s test).

To further quantify cell segmentation performance, we randomly selected eight 700 × 700 μm² regions from multiple samples and processed them with NeuroPixelAI using identical parameters for motion correction, image denoising and cell segmentation; the resulting neuron counts were normalized to the maximum number obtained across all regions. Under the same algorithmic pipeline and parameter settings, the parabolic collector module yielded 2.5-fold more segmentable neurons than the 4×/0.28-NA objective path (Fig. 2H,I).

### Denoising Algorithm Testing

To evaluate the performance of various denoising algorithms, we compared their effects on data with a 0 dB signal-to-noise ratio (SNR) across different frame rates (Extended Data Fig. 8B), where SNR is defined as the ratio of signal power to noise power:

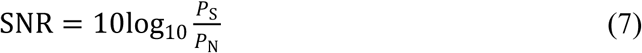

where *P*_S_ represents the signal power, and *P*_N_ denotes the noise power.

We first acquired clean calcium imaging videos at 100 Hz frame rate, 200 μm depth, and 300 × 300 μm² FOV using NAOMI^52^. These datasets were then downsampled to generate frame rates spanning 1-30 Hz, followed by noise injection to simulate experimental conditions. The noise inherent in calcium imaging data predominantly consists of photon shot noise, dark current noise, and readout noise^53^. Among these, readout noise is characterized by a Gaussian distribution and the power of dark current noise is relatively low and stable^54^. The principal component of noise, however, is the unavoidable shot noise, which occurs due to quantum fluctuations during the photon acquisition process and is also known as Poisson noise. The theoretical noise model can thus be described as:

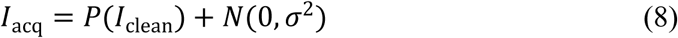

Considering digital resolution (image bit depth), the generation of MPG noise can be reformulated as:

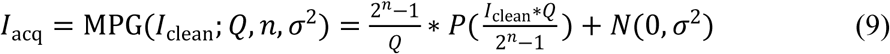

here, *I*_clean_ represents the photon distribution in the pristine data, and *I*_acq_ represents the corresponding noise-contaminated photon distribution. *P* is a function operator of the signal-related Poisson noise, *N* represents signal-independent zero-mean Gaussian noise with a variance characteristic σ^2^, n denotes the bit depth of the captured image (set here as n = 16), MPG(·) is the operator generating MPG noise, used to simplify the mathematical expression and *Q* represents the relative number of photons per pixel captured by the detector. In this work, different *Q* values were used to simulate calcium imaging data with various SNRs. Given that Gaussian noise constitutes a smaller component of the total noise, its mean is set to zero, and its variance is set to σ^2^ = 1000, hence = 31.623.

To calculate SNR in calcium imaging videos, for a noise-free calcium imaging video, we calculate the signal power by squaring the intensity of each pixel in every frame throughout the video and then dividing by the total number of pixels. This can be expressed specifically as:

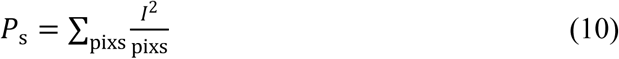

where *I* represents the pixel value of a single pixel in a particular frame, and pixs is the sum of the number of pixels across all frames. After calculating the signal power, similarly, the noise power is determined by computing the squared difference between the pixel values of the noisy video and the noise-free video at each point, summed and then divided by the number of pixels:

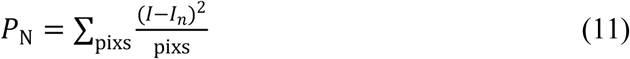

where *I_n_* denotes the pixel value in the noisy video for a single pixel in a particular frame. Finally, the SNR is calculated according to Equation (7).

### Assessment of Cell Segmentation Algorithms

Neuronal spatial segmentation is typically framed as a multi-class task, assigning each pixel to its corresponding neuron. To evaluate algorithm performance, we used standard metrics: Precision, Recall, and the F1 Score (Extended Data Fig. 8D-F). Furthermore, the computational speed of the segmentation algorithms was included as a performance benchmark.

Precision, also known as positive predictive value, represents the proportion of true positives among the samples predicted as positive. It can be described by the following equation:

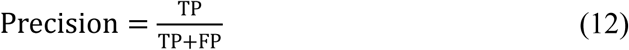

where TP is true positive, FP is false positive.

Recall, also referred to as sensitivity, indicates the proportion of true positives out of all actual positive samples. It can be represented as:

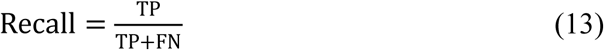

where FN is false negative.

The F1 Score, which emerges from the concepts of Precision and Recall, typically presents a trade-off between these metrics: high Precision often coincides with lower Recall, and vice versa. To provide a balanced consideration of these metrics, the concept of the F1 Score was introduced:

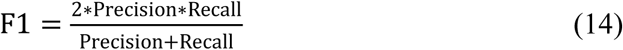

Regarding the processing speed of the algorithms, we have simply evaluated it in frames per second.

The specific method of testing employed cross-validation. In the experimental setup, we used six sets of calcium imaging videos at 3.51 Hz with 6,500 frames (512 × 512 pixels), manually annotating nearly 600 ROIs as ground truth using Cellpose 2.0. For algorithms requiring network training, five sets of data were used for training and the remaining set served as the test set, completing the evaluation across all six video sets in a repetitive manner. For algorithms that did not require training, parameter adjustments were made prior to testing. During each algorithm’s testing phase per video set, we concurrently recorded Precision, Recall, F1 Score, and image processing speed as metrics of evaluation.

### Baseline Correction Algorithm

In certain experiments, we observed a concurrent reduction in neuronal action potential firing and total [Ca^2+^]_i_ fluctuations, manifesting as baseline variations in the recorded data. To accurately extract spikes from individual neurons despite ongoing baseline fluctuations, we incorporated a baseline correction algorithm based on Cubic Spline Approximation (Smoothing) (csaps) into our software. Upon inputting data, the smoothing parameter of csaps is adjusted to smooth = 1e-8. The data fitted using this method serve as the baseline for calculating the change in neuronal fluorescence, Δ*F*/*F*_0_:

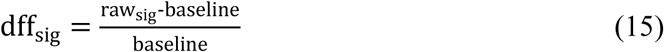

where baseline is the baseline obtained by fitting the original data using the csaps algorithm, raw_sig_ represents the original data, and dff_sig_ denotes the final calculated change in neuronal fluorescence intensity Δ*F*/*F*_0_.

### Spike Inference Algorithm Testing

To evaluate algorithm performance, we computed the Pearson correlation coefficient between electrophysiologically recorded membrane potentials and inferred spike trains from original calcium imaging data processed by each algorithm. Analyses were performed across multiple standardized noise levels (Extended Data Fig. 8H-J).

Specifically, each algorithm processed original calcium signals to generate inferred spike probability time series. These inferred traces were then compared to simultaneously recorded membrane potential signals, which were converted into standardized spike probability estimates using established methods. Correlation coefficients served as the primary metric to: (1) compare algorithm performance across different standardized noise levels, and (2) assess the stability of each algorithm’s performance at a fixed noise level.

The definition of standardized noise level is given by^55^:

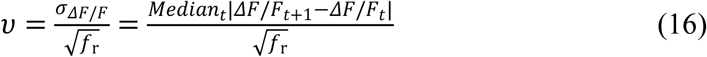

here, based on optical imaging principles with finite granularity noise where the detector collects *N* photons per second, the noise magnitude is 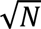. Thus, the baseline noise level is expressed as: 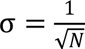 (in percentage units). Considering the frame rate, *f*_r_, for each frame, the formula’s can be replaced entirely by *N*/*f*_r_, resulting in a baseline noise level per frame of: 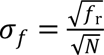. To eliminate the factor of frame rate, the standardized noise level υ is defined as the Equation (16).

To convert the potential signals into spike probabilities, we performed binarization followed by Gaussian kernel convolution. The original spike timing data in Fig. 3F and Extended Data Fig. 8H show discrete signals. After binarization and convolution processing, the resulting spike probabilities are used to evaluate the performance of the algorithms.

### Iterative Correlation Sorting Algorithm

Even in the absence of recorded specific stimulus times during experiments, neuronal signals can be sorted by their inherent temporal characteristics of local firing potentials. Initially, the data from all neurons are aggregated into a single time series (representing the overall temporal signal of all neurons). The 100 largest-amplitude time points from this series as template for cross-correlation-based alignment of all neuronal signals. Subsequently, the average of the top 10 most highly correlated neuronal signals post-alignment is taken as the new template to reorder all neuronal signals. After 30 iterations, the resulting correlation sorting plot appears in the lower-right image frame of software Tab 2.

### Development and Implementation of NeuroPixelAI for Large-Field Image Analysis on a High-Efficiency Software Platform

In order to address the challenges associated with processing extremely large image datasets from wide-field observations and to furnish a platform capable of one-click, high-speed processing, we developed NeuroPixelAI using PyQt5. The workflow is compartmentalized into six distinct stages: motion registration, image denoising, cell segmentation, signal extraction (baseline correction), spike inference, and correlation-ranking clustering. Given the dynamic changes inherent to biological activity, tissue expansion, and mechanical fixation during imaging processes, there often occurs displacement within the imaging field. To counteract this, our initial step involves motion registration. We adopted a phase-correlation-based registration method, similar to that used in Suite2p^25^, which iteratively computes the average of the most correlated frames in a video to serve as a reference image. Subsequent frames are then aligned with this reference image to achieve motion registration. We enhanced this algorithm by incorporating dynamic memory allocation, allowing it to handle larger data structures effectively. Moreover, we reduced the adjustable parameters from twenty to four input parameters, and introduced options for non-rigid registration and bidiphase adjustment, significantly reducing parameter redundancy and simplifying usability. The second step is image denoising. Calcium imaging data, due to the transient, low-amplitude accumulations of intracellular calcium ions and the rapid dynamics involved, coupled with the limitations of electronic detection equipment, are susceptible to various types of noise contamination, such as shot noise, dark current noise, and readout noise^54^. This poses significant challenges for high-performance imaging, such as dendritic imaging and deep tissue imaging. We referred to algorithms such as DeepCAD-RT^29^ and SRDTrans^30^ (Extended Data Fig. 8B,C), which are known for their speed, precision, and parameter-free operation. Upon completion of their training phase, these algorithms can effectively eliminate various types of white noise from multiphoton image data. Consequently, we integrated the denoising function with the prior motion registration into a unified module (Image Enhancement) within NeuroPixelAI and included visualization capabilities. The third step involves cell segmentation. To effectively handle the diverse cellular morphologies and distributions across different brain regions and to facilitate efficient one-click processing, we developed an optimized processing pipeline. Building upon Cellpose 2.0^56^, we implemented parameter optimization and established a region-adaptive framework to achieve high-precision, high-throughput cellular segmentation across large fields of view. The streamlined workflow enables efficient one-click processing within NeuroPixelAI, while retaining manual curation capabilities for user-guided refinement. Algorithm performance was validated against multiple alternatives (CNMF-E^26, 57^, Cellpose^25, 56^, SUNS^58^) using both raw and denoised data (Extended Data Fig. 8D-F). The fourth stage in our methodology is signal extraction. Based on the cellular masks generated during the third stage of cell segmentation, we sum the pixel values within each ROI in every frame to obtain the raw temporal data of individual neurons. Additionally, we offer an optional baseline correction method utilizing the cubic spline interpolation algorithm. This approach is designed to correct baseline fluctuations in the signals, which may arise due to changes in the concentration of optical indicators within cells, photobleaching, or specific biological activities (Extended Data Fig. 8G). This method is particularly advantageous for subsequent spike inference during extended recordings of multiple signals. The final stage involves spike inference. Commonly used optical imaging signals, particularly calcium signals, serve as indirect, typically non-linear, and low-pass filtered representations of more fundamental variables of interest—namely, somatic action potentials (spikes)^55, 59–61^. Historically, many studies have analyzed raw calcium signals directly^39, 62^ or employed various deconvolution algorithms, such as OASIS (Suite2p, CaImAn)^62–65^ or custom-written toolbox^66^, to approximate signals closer to the original action potentials for analysis. However, in practical imaging scenarios, the single-pulse responses of different neurons, and even different temporal segments within the same neuron, vary significantly^66^. Moreover, parameters like pulse width, often preset in deconvolution algorithms, are challenging to estimate accurately prior to analysis. This variability leads to substantial standard deviation fluctuations in the precision of traditional deconvolution algorithms, especially when processing data with low SNRs. We tested various deconvolution algorithms^25, 26, 67^ and the deep learning algorithm Cascade using datasets provided by CRCNS^31, 32^ and to restore calcium signals to electrophysiological signals (Extended Data Fig. 8H-J). Our findings demonstrate that the deep learning-based Cascade algorithm significantly outperforms traditional deconvolution methods when dealing with calcium imaging data of varying SNR levels. We have integrated Cascade’s testing methodology into NeuroPixelAI, along with enhancements such as visualization features for neuronal signals during processing and a signal-correlation iterative sorting algorithm (Extended Data Fig. 8K). These enhancements allow for an intuitive observation of the temporal signals and overall correlation characteristics of neurons post-processing.

### Instrument (Meso2P) Validation

For in-house-bred *Camk2a-Cre*::Ai162D mice, imaging comparisons were conducted on the same cortical plane (250-µm depth) without and with the parabolic lateral collection module. Images were acquired in high-speed (2,048 × 2,048 pixels/frame, 7.67 Hz) or high-resolution mode (4,096 × 4,096 pixels/frame, 3.83 Hz) using 130-mW excitation power. Fluorescence collection efficiency was assessed using the parabolic lateral collection module at 330-µm depth below the cortex with 90-mW imaging power. Twenty-eight-layer volumetric imaging was achieved using parabolic lateral collection, time-division multiplexing, and a piezo-driven z-scanning module. This enabled simultaneous imaging at 2,048 × 2,048-pixel resolution across a 6 × 6 × 0.5 mm^3^ volume (from 60 to 560 µm below the cortical surface) at 1.1 Hz with 250-mW excitation power. For 8-10-week-old *Camk2a-Cre*::Ai162D mice, the red-shifted opsin pAAV-EF1α-DIO-C1V1(t/t)-Ts-mCherry-WPRE (Obio Technology), a widely used optogenetic actuator, was injected before cranial window implantation. After a 4-week recovery period for viral expression, multi-region 2P optogenetic stimulation was conducted with 50-ms light pulses, 10 pulses per site and a 4-s interval between adjacent quadrants for 5 cycles at a depth of 220 µm (4×/0.28-NA objective). The imaging and stimulation powers were 180 mW and 79 mW, respectively. Single-cell 2P optogenetic stimulation used 50-ms light pulses every 16 s (10 cycles; cycles 4-9 as control) at a depth of 220 µm (Nikon 16×/0.8-NA objective). The imaging and stimulation powers were 20 mW and 50 mW, respectively.

### Photodamage Assessment

Uniform 28-layer imaging of 500- µm- thick cortex was performed for 1 h under 0 mW (control), 250 mW, 350 mW and 450 mW excitation. In addition, identical imaging acquisitions were carried out at a depth of 400 µm for parallel comparison (parabolic lateral collection module, 4× objective). Twenty hours post-imaging, each mouse received an intraperitoneal injection of heparinized saline (80 U/mL) prior to anesthesia to optimize transcardial perfusion efficiency. Mice were then deeply anesthetized with 1% pentobarbital sodium, followed by transcardial perfusion with heparinized saline (20 U/mL) and subsequent fixation with 4% paraformaldehyde (PFA). Brains were post-fixed overnight in 4% PFA at 4 °C, cryoprotected sequentially in 20% and 30% sucrose/PBS, and sectioned at 35-µm thickness. Photodamage was assessed by immunofluorescence: sections were blocked in 5% normal donkey serum/PBST (PBS + 0.3% Triton X-100) for 1 h, incubated with primary antibody (HSP70/HSP72 monoclonal antibody, Enzo Life Sciences #ADI-SPA-810, 1:200; CD31 Antibody, R&D Systems #AF3628, 1:200; NeuN Polyclonal antibody, Proteintech #26975-1-AP, 1:200) overnight at 4 °C, and labeled with secondary antibody (Donkey anti-Mouse IgG (H+L) Highly Cross-Adsorbed Secondary Antibody, Alexa Fluor™ Plus 647, Invitrogen #A32787, 1:500; Donkey anti-Rabbit IgG (H+L) Highly Cross-Adsorbed Secondary Antibody, Alexa Fluor™ Plus 488, Invitrogen #32790, 1:500; Donkey anti-Goat IgG (H+L) Cross-Adsorbed Secondary Antibody, Alexa Fluor™ 568, Invitrogen #A11057, 1:500) for 2 h at room temperature, and sealed with Antifade Mounting Medium with DAPI (Beyotime Biotechnology #P0131). Six cortical regions per brain were imaged using SLIDEVIEW™ VS200 universal slide scanner (Evident Scientific, Tokyo, Japan). Photodamage regions were segmented using the Threshold tool in ImageJ software (v1.53t, NIH)^68^, with a fixed threshold applied uniformly across all images. Fluorescence intensity within segmented photodamage regions was quantified and normalized to the 0-mW control group. Statistical comparisons were conducted using one-way ANOVA followed by Tukey’s post hoc test (n = 3 mice per group).

### Optical resolution calibration

To calibrate the optical resolution of our multiphoton imaging system, we computed the 1/e intensity radius of the focal spot based on Gaussian functions fit to integral representation of the electric field near the focus of a diffraction-limited focus^69^:

Lateral radius:

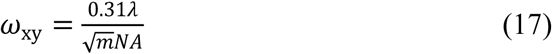

Axial radius:

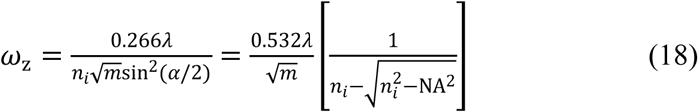

α = arcsin(NA/n) in radians. ω is the 1/e radius. For the 1/e^2^ radius multiply by 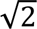. For the FWHM multiply by 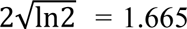. m = order of excitation: m = 1 for conventional excitation, m = 2 for 2 photon excitation, etc.

For an excitation wavelength of 920 nm and an excitation NA of 0.28, these equations yield theoretical lateral and axial FWHM values of 1.07 µm and 19.36 µm, respectively. The corresponding bead-based measurements yielded lateral and axial FWHM values of approximately 1.2–1.8 µm and 21–24 µm, respectively, indicating modest broadening relative to the theoretical estimates.

To quantitatively characterize the actual resolution of the system, fluorescent microspheres (0.2 μm diameter, catalog no. 7-3-0020, BaseLine) were embedded in 1.2% agarose gel at a concentration of 0.585 μl/ml. Using the gel-embedded microspheres, the lateral and axial point spread functions (PSF) were measured at various positions from the center to the periphery of the FOV under two configurations employing 4× and 16× objectives, respectively (Extended Data Fig. 1E,F). The PSF profiles were fitted with a Gaussian function and reported as full width at half maximum (FWHM).

## Quantification and statistical analysis

### Image Processing and Statistical Analysis

All image data were processed using NeuroPixelAI for motion correction, denoising, cell segmentation, signal extraction, and spike inference, with additional baseline correction and iterative sorting applied based on data characteristics. Outputs include but are not limited to: enhanced images, neuron masks and properties, raw pixel values, ΔF/F0 signals, spike probabilities and inferred spikes. Denoising improves visualization and downstream processing stability, but the biological and technical conclusions were validated using raw or non-denoised data comparisons (Extended Data Fig. 12). Statistical analyses used MATLAB® R2023a (v9.14.0, Mathworks). Normally distributed continuous data are presented as mean ± s.d. All statistical details (the size and type of individual samples, n) are specified in figure legends.

### Cascade Model Training

Given variable imaging rates across experiment, we trained multiple spike-inference models using the “Global EXC” dataset^55^ at diverse frame rates and Gaussian smoothing kernel parameters. This dataset covers multiple indicators (e.g., GCaMP6f, GCaMP6s, OGB-1, GCaMP5k, Cal-520, R-CaMP1.07, and jRCaMP) and brain regions (e.g. the visual cortex, somatosensory cortex, hippocampus, several regions of the zebrafish forebrain, and the olfactory bulb), including some inhibitory neurons. The training noise levels ranged from standard noise level υ 2 to 9. The smoothing parameter was empirically set to the inverse of the frame rate multiplied by 1.5; for instance, at 7.67 Hz, the smoothing parameter was 195 ms, and at 30 Hz, it was 50 ms. The causal ground truth smoothing kernel was set to 0 ms.

## Cortex-wide neuronal dynamics during whisker stimulation

### Experiment Paradigms

An air-puff nozzle was positioned in front of head-fixed *Camk2a-Cre*::Ai162D mice. After a 5-min baseline recording, mice received 10 cycles of air-puff stimulation (15-s duration each) with 45-s interstimulus intervals. Air-puff delivery was synchronized with imaging using a programmable solenoid valve controller. Neuronal calcium signals at 300-µm depth within a 6 × 6 mm^2^ cortical area were continuously recorded with the parabolic lateral collection module (2,048 × 2,048 pixels, 7.67 Hz).

### Brain Region Division and Regional Neuron Statistics

Brain regions were delineated and masks were created referencing the Allen Brain Atlas^70^. Imaging areas were proportionally registered based on coronal and horizontal planes, and all neurons were assigned to brain regions based on centroid coordinates processed by NeuroPixelAI (Fig. 4C and Extended Data Fig. 9A).

### Definition of Active (Coding) Cells

Active neurons were identified based on the inferred spikes output from NeuroPixelAI. Neurons that exhibited an average action potential firing rate more than twice the resting state firing rate of mice during ten stimulus trials were considered “active”. Under Gaussian distribution, non-coding neurons should have a 95% probability of firing at less than twice the resting rate in a single experiment (Fig. 4E and Extended Data Fig. 9C,D).

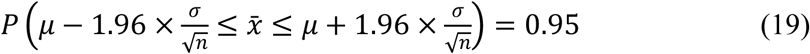

### Decoding Neural Population Activity Using SVM Decoders

To investigate the encoding responses of neural population activity in various brain regions to air puff stimuli, we employed SVM decoders with a linear kernel for population decoding^71^. For each mouse, we utilized the neural activity data collected from 10 s before to 10 s after each trial, comparing it to resting state activity. Decoders were trained on 80% of trials using a sliding 2-s time window and tested for accuracy on the withheld 20% of trials. To avoid bias and overfitting, we repeated decoding 512 times with random trial selections and averaged the measured values.

In tests examining the relationship between decoding accuracy and the number of neurons, we randomly sampled increments of 200 (5%) neurons from a range of 200 to 4,000 neurons for decoding tests. The average decoding accuracy over the 15 s duration of the air puff stimulus was recorded as the decoding accuracy for each group of neurons. Each mouse underwent this random sampling 10 times to mitigate the random errors caused by uneven distribution of neurons across brain regions.

### Global Neural Peak Activity Conduction Bar Graph

In each trial, we observed specific directional inter-regional transmission of neural group signals. To visualize fluctuations, we divided the AP = [-3 mm, 3 mm] area into 16 bins and categorized the neurons within each. Since calcium signaling serves as a low-pass, non-linear proxy for the sequence of cellular action potentials (spikes), and the spike probabilities inferred by the cascade algorithm can be considered a Gaussian kernel convolution of the spike sequence, generally featuring a relatively higher temporal resolution, we averaged the spike probabilities S (*N*_b_ × *N*_t_, where *N*_b_ is the number of regional bins and *N*_t_ is the number of time points) across all neurons in each bin to obtain S_bin_. This was then normalized to the peak values within a 2 s window before and after each stimulation to generate the peak transmission matrix T:

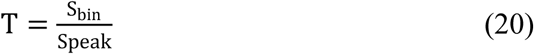

Subsequently, we generated a 3D visualization of matrix T, showing normalized signal fluctuations across regions and the sequence and timing of peak arrivals in response to a single stimulus (Fig. 4J and Extended Data Fig. 9H,I).

### Dimensional Reduction and Visualization of Neural Representations Using CEBRA

To investigate whether global network-wide reorganization of activity patterns occurs during adaptation to repeated sensory stimuli, we employed the CEBRA^72^ algorithm for dimensional reduction and visualization of neural representations. For our study, the CEBRA algorithm was applied to neuron signals that vary over time, specifically in the CEBRA-Time mode. The input consisted of spike probabilities from neurons across 10 trials. During the application of the CEBRA algorithm, we employed a nonlinear convolutional neural network model, the “offset10-model,” for self-supervised training. The batch size for training data was set at 512, with a learning rate of 0.0003 for the optimizer, and a maximum iteration count of 10,000. The multi-dimensional neural signals were reduced to a three-dimensional space, with both the training and testing phases accelerated by high-performance graphics processing units. To enhance the visualization of the dimensional reduction results, we employed pseudocolor coding that changes over time to represent each time point in the reduced data, as shown in Fig. 4K.

## Canonical Correlation Analysis and Principal Component Analysis of Pattern

### Shifts and Neural Activity Weight Changes

To investigate the correlation between activity patterns across different cortical areas and their gradual representational changes over successive experiments, we employed Canonical Correlation Analysis (CCA) to analyze the shifts in shared activity patterns within pairs of experiments and brain regions. For each experiment, the spike probability was captured within a 1.5 s window before and after stimulus presentation, segregated by brain region. For a given pair of trials within a specific brain region, we represented the dynamical features of this area during the two trials with matrices X and Y, sized *N*_t1_ × *N*_1_ and *N*_t2_ × *N*_2_, respectively. Here, *N*_t1_ and *N*_t2_ denote the total number of time points in the group dynamic matrices for the two trials, while *N*_1_ and *N*_2_ represent the number of neurons in each trial. Following the standard methodology of CCA, we identified two sets of loading vectors, {*W_i_*} and {*v_i_*}. Each vector represents the weight distribution of each neural ensemble in a shared activity pattern observed across the two trials. The index i, ranging from 1 to the minimum of the dimensions of X and Y, denotes each mode. We determined these modes such that the projections of the neural activity fluctuations, X and Y onto *W_i_* and *v_i_*, achieved maximum correlation between the two ensembles.

The maximization of the correlation between these projections is expressed as:

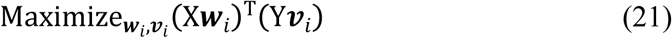

subject to the normalization constraint: 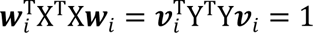. Given this normalization condition, the quantity (X*W_i_*)^T^ ⋅ (Y*v_i_*) equals the correlation coefficient *r_i_* of the activity modes, X*W_i_* and Y*v_i_*, in the two different trials.

In order to explore the origins of gradual shifts in global activity patterns in response to stimuli and the involvement of different brain regions in these changes, we initially normalized the correlation coefficient *r*_1_ for mode 1 extracted between different trials for each brain region, and visualized this data using a correlation heatmap (Fig. 4M). Subsequently, for each trial, we conducted cross-CCA between brain regions. Specifically, within a given independent trial, we performed cross-CCA on the dynamic matrices of the M, S, V, and RSP regions sequentially, obtaining pairwise correlated modes *w*_areaij_ and *v*_areaij_. These modes were then averaged according to brain region:

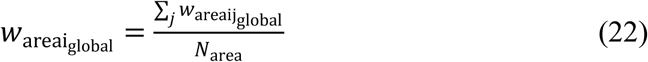

this calculation yielded a single-trial local loading vector *W*_areaiglobal_ for each brain region, where *N*_area_ denotes the number of brain regions. Here, *W*_areaiglobal_ reflects the absolute contribution of all neurons in that region to the global activity pattern during that trial. We posited that as trials progressed, the variation in these patterns would manifest in slow changes in the weight distribution of neuronal responses across different regions (Extended Data Fig. 9M). Accordingly, we mapped the changes in neuronal contribution weights *W*_weightchange_ across trials for different brain regions using the weight change matrix:

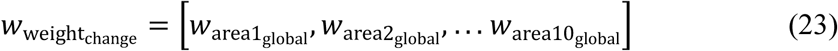

these changes were visualized and further analyzed using PCA to reduce the dimensions of the weight change matrix. The dominant latent component, PC1 (Extended Data Fig. 9L), revealed gradual shifts in the contribution weights of neurons across different brain regions.

Additionally, we focused on the collective contributions of each brain region to the global activity pattern across ten trials. We projected the group activity of different regions onto the global activity pattern and averaged it across neurons, i.e.,

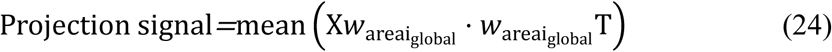

It was observed that, despite all regions participating in the encoding response to air-puff stimuli and undergoing similar changes in neuronal contribution weights through successive adaptations, the activity patterns in regions S and V contributed more significantly to global encoding compared to regions M and RSP (Fig. 4L and Extended Data Fig. 9K).

## Dissecting layer 2/3 and layer 5 activity simultaneously across cortex during anesthesia

### Experiment Paradigm

A facemask was positioned in front of head-fixed *Camk2a-Cre*::Ai162D mice and adjusted to fit snugly (gas flow 1.0 L/min). After a 30-min adaptation, 10-min awake baseline imaging was performed. Isoflurane was then increased to 2% for 20-min anesthesia induction, followed by a 10-min stabilization. Isoflurane was reduced to 0% for the recovery imaging. Simultaneous dual-layer imaging at 300-μm and 560-μm depth (6 × 6 mm^2^ FOV) used the parabolic lateral collection module at 2,048 × 2,048-pixels resolution (7.67 Hz).

### Analysis of Synchrony in Neuronal Population Activity

To investigate changes in synchrony between deep and shallow neuronal activity during various stages of isoflurane anesthesia, we computed the Pearson correlation coefficient, *R*, for individual neurons against the remaining population activity within each window. This was performed using a 15 s time window with a 3 s step size:

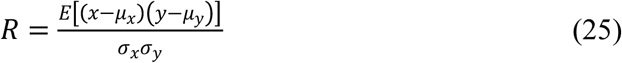

where *x* represents the activity of a single neuron within the window, *y* represents the activity of the excluded neuronal population and μ_x_, μ_y_ denote the means of *x* and *y*, respectively, while σ_x_ and σ_y_ represent their standard deviations. *E* signifies the arithmetic mean. The mean correlation across all neurons was then used to quantify the synchrony of population activity within that window (Fig. 5F,G). Subsequently, we defined the probability of anesthesia-related events based on synchrony, using a 650 ms time window. Specifically, the peak synchrony during wakefulness and the lowest synchrony during anesthesia were used as thresholds to calculate the probability of population activity synchrony crossing these thresholds across multiple experiments.

To compare the synchrony differences between deep and shallow layers under different anesthetic states, we also analyzed the synchrony within the first 180 s following the onset of synchronized activity, typically between 40 s and 220 s after administration and cessation of isoflurane. Both during the induction and the awakening phases, neurons in both layers exhibited synchronized activity, with deeper layers showing higher synchrony than shallow layers, and induction phases showing higher synchrony than anesthesia phases (Fig. 5H, \*\*\**p* < 0.001; Tukey test; neurons = 19,808 ∼ 23,684; n = 3 mice).

### Granger Causality Analysis

Based on experimental observations, we hypothesized that multiple “synchronized activities” during the induction of anesthesia actually represent specific directional flows rather than complete synchrony. To infer causality in activity between anterior and posterior (AP) cortical regions during individual synchronized events (synchronization peaks), we uniformly divided the cortical populations within an AP range of [-3 mm, 3 mm] into 16 segments. We then analyzed Granger causality among these 16 groups of neurons using the MVGC Multivariate Granger Causality Toolbox (https://github.com/lcbarnett/MVGC1)^73^. The input matrix included three dimensions: AP bins, time points, and peaks number, recording activity of different AP bins’ neuronal populations within a 4 s interval (2 s before to 2 s after each peak). This multivariate Granger causality toolkit, based on a vector autoregressive model, calculates Granger causality where the model order (time lags) was estimated by Akaike Information Criteria with the maximum set to 20. Pairwise conditional causality was then calculated, where causality includes directional information: the direction “from” an X module “to” a Y module indicates that activity in X preceded that in Y. Granger causality values that reached \**p* < 0.05 (Granger F test) after applying the false discovery rate for multiple-comparison correction (Fig. 5I).

### Analysis of Global Neural Activity Transmission During Synchronous Peaks

During periods of anesthesia, multiple “synchronous” peaks are not entirely synchronized and exhibit various directions of transmission. It is only during deep anesthesia that these “synchronous” peaks approach complete synchronization. To verify this, similar to the repetitive air-puff whisker stimulation experiment, we initially utilized the spike probability outputs from NeuroPixelAI to perform regional peak detection, resulting in the generation of a spike transmission matrix T_peaks_. Here, peak detection was conducted using a MATLAB-based local thresholding algorithm, with parameters set at a threshold of 0.001, minimum peak height of 0.04, and minimum peak distance of 20. The structure of T_peaks_ is represented as *N*_b_ *N*_t_ ⋅ peaks_num_, where *N*_b_ denotes the number of regional bins, *N*_t_ indicates the number of time points (spanning [-2 s, 2 s] across 31 frames at a frame rate of 7.67 Hz), and peaks_num_ represents the detected number of peaks. A three-dimensional visualization of the signals within T_peaks_ facilitates the generation of a global map of neural peak activity transmission (Fig. 5J and Extended Data Fig. 10C).

To further visualize the dynamic transmission process of “synchronous” signals throughout the anesthesia induction, we rendered a global dynamic transmission map of peak activities at the resolution of individual neurons (Fig. 5J, Extended Data Fig. 10A,C, and Supplementary Video 6). In this representation, yellow signifies the origin regions of the signals, while purple denotes the receiving regions. Specifically, for each “synchronous” event, we normalized the spike probability for each neuron around the event ([-2 s, 2 s]), based on peak values. We then calculated the response delay, which is the difference between the peak positions of individual neurons and the peak position of the group activity. If the response delay is less than zero, the neuron is identified as the source of the event; if greater than zero, the neuron is considered a receiver. Based on the response delays of various neurons, we were able to ascertain the global dynamics of peak transmission within the imaging field area (6 × 6 mm^2^): for each layer in T_peaks_, or each peak, we determined the times *t*_AP_ at which different AP bins reached their peak and calculated the standard deviation across these bins:

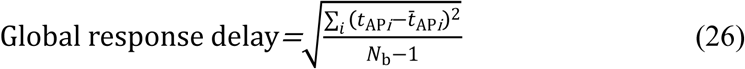

the global response delay reflects the fluidity of each spike’s global response, illustrating the progression from incomplete to almost complete synchronization of multiple “synchronous” peaks during the entire period of anesthesia induction (Extended Data Fig. 10B). To mitigate the impact of spurious spikes detected during peak detection, we applied a sliding window averaging and normalization over 20 spikes.

Through the assessment of global response delays, we hypothesize that during the process of anesthesia induction, there exists a gradual change in neural group activities and a redistribution of neuronal activity weights, associated with an increase in ‘synchronized’ peak events. To investigate this, we categorized the detected peaks into deciles, representing 10% of the peaks per group, and initially computed the normalized CCA mode1 correlation matrices between these groups (Fig. 5L). Subsequently, we demonstrated the matrices of changes in neuronal weight across these ten groups of peaks (Extended Data Fig. 10D). The correlation coefficient matrices were derived through pairwise CCA of each group of peaks. The CCA of the neuronal weight change matrices encompassed two elements: the neural dynamics matrix for the period [-2s, 2s] around a specific group of peaks, and the average neural dynamics matrix for all other peaks except for the group in question. Following each CCA, the weight matrices *w_i_* were normalized (ensuring the sum of weights for all neurons equaled one) and subjected to iterative correlation ranking.

In addition, we analyzed the proportions of forward versus backward transmission peaks (Fig. 5L; \**p* < 0.05) and the transmission from deep to shallow layers versus from shallow to deep layers during the entire anesthesia induction process (Extended Data Fig. 10F; \*\**p* < 0.01). For the anterior-posterior peak transmission, we calculated the difference in response delays between the first 50% of bins and the last 50% of each event to determine whether the anterior or posterior regions reached peak values first. For the deep-to-shallow layer transmission, we calculated the differences in response delays across 16 bins corresponding to different depths (Extended Data Fig. 10E). A negative response delay indicated that, for a given event, the signal was transmitted from deeper to more shallow layers; conversely, a positive response delay suggested transmission from shallow to deeper layers. These findings align with previous studies^74–77^, indicating that during the anesthesia induction phase, global signal propagation predominantly occurs from posterior to anterior regions, with notable instances of reverse transmission, primarily occurring from deeper to more shallow layers.

## Dynamic monitoring of MNP distribution and vascular occlusion in the living brain

### Experiment Paradigm

Prior to imaging, male C57BL/6J mice (Charles River Laboratories #219) under isoflurane anesthetized received 200-μL retro-orbital injections of 5% FITC-Dextran or RITC-Dextran. Twenty-eight-layer structural vascular imaging at 450-µm depth cover 6 × 6 × 0.45 mm^3^ (4× objective) or 1.6 × 1.6 × 0.45 mm^3^ volumes (16× objective), using the parabolic lateral module at 2,048 × 2,048 pixels resolution (7.67 Hz/layer). For real-time MNPs tracking, dye-injected mice received 100-μL tail vein injections of 1 mg/mL 5-µm polystyrene (PS) microplastics (MPs, Baseline #7-1-0500) or 200-nm PS nanoplastics (NPs, Baseline #7-3-0020) during single-layer imaging. Repeat 28-layer structural imaging was performed at 12-h and 24-h post-injection.

### Blood Flow Velocity Detection

Blood flow velocity was measured using an established method^78^. Fluorescent dye injection rendered red blood cells (RBCs) as dark particles. RBC displacement was tracked by 1k Hz high-speed line-scanning within ROIs. Velocity was calculated from pixel-wise displacement over 10-s line-scanning recordings. Given MNP occlusion-prone capillaries, we analyzed single-RBC-perfused vessels. Measurements occurred at: 12/24-h post-injection. Blood flow was not assessed during tail vein injections to eliminate artifacts.

### Blood-Brain Barrier Integrity Assay

Male C57BL/6J mice were divided into: sham-operated, 5-day post-surgery, and 14-day post-surgery groups. Each received 125-μL tail vein injections of 2% Evans Blue (Beyotime Biotechnology #ST3273) 1.5 h before brain collection. Mice were anesthetized with 1% pentobarbital sodium (i.p.), then transcardially perfused with heparinized saline (20 U/mL) and 4% PFA. Evans Blue extravasation was visually assessed in whole brains to qualitatively evaluate BBB integrity.

## Declaration

### Availability of data and materials

Meso2P analysis code is available at https://github.com/Yourswear/Meso2P, with representative air-puff and anesthesia datasets at https://doi.org/10.5281/zenodo.20956973. NeuroPixelAI source code is available at https://github.com/Yourswear/NeuroPixelAI, with a representative example dataset at https://doi.org/10.5281/zenodo.16595850. Additional raw data supporting the study are available from the corresponding author upon reasonable request, and information needed to reanalyse the reported data is available from the lead contact. This study did not generate new unique reagents.

### Author Contributions

Conceptualization: B.L.

Methodology: B.L., J.H.H., S.P.L., C.Y.L., M.Z.

Biological experiments: Y.F.Z., J.H.H, X.Y.G, H.Y.Y.

Zemax optical simulation: J.H.H., F.X.;

Investigation: J.H.H., Y.F.Z.

Visualization: J.H.H., Y.F.Z., M.Z.

Funding acquisition: B.L., Y.M., L.C.

Project administration: B.L.

Writing: B.L., J.H.H., Y.F.Z., H.D.L., H.Y.Y., Y.M., L.C., J.C.W.

## Acknowledgements

We thank members of the Li research group, especially Yaoguang Zhao, Peng Deng, and Rui Liu, for their help and valuable discussions.

## Extended data

**Extended Data Fig. 1.**
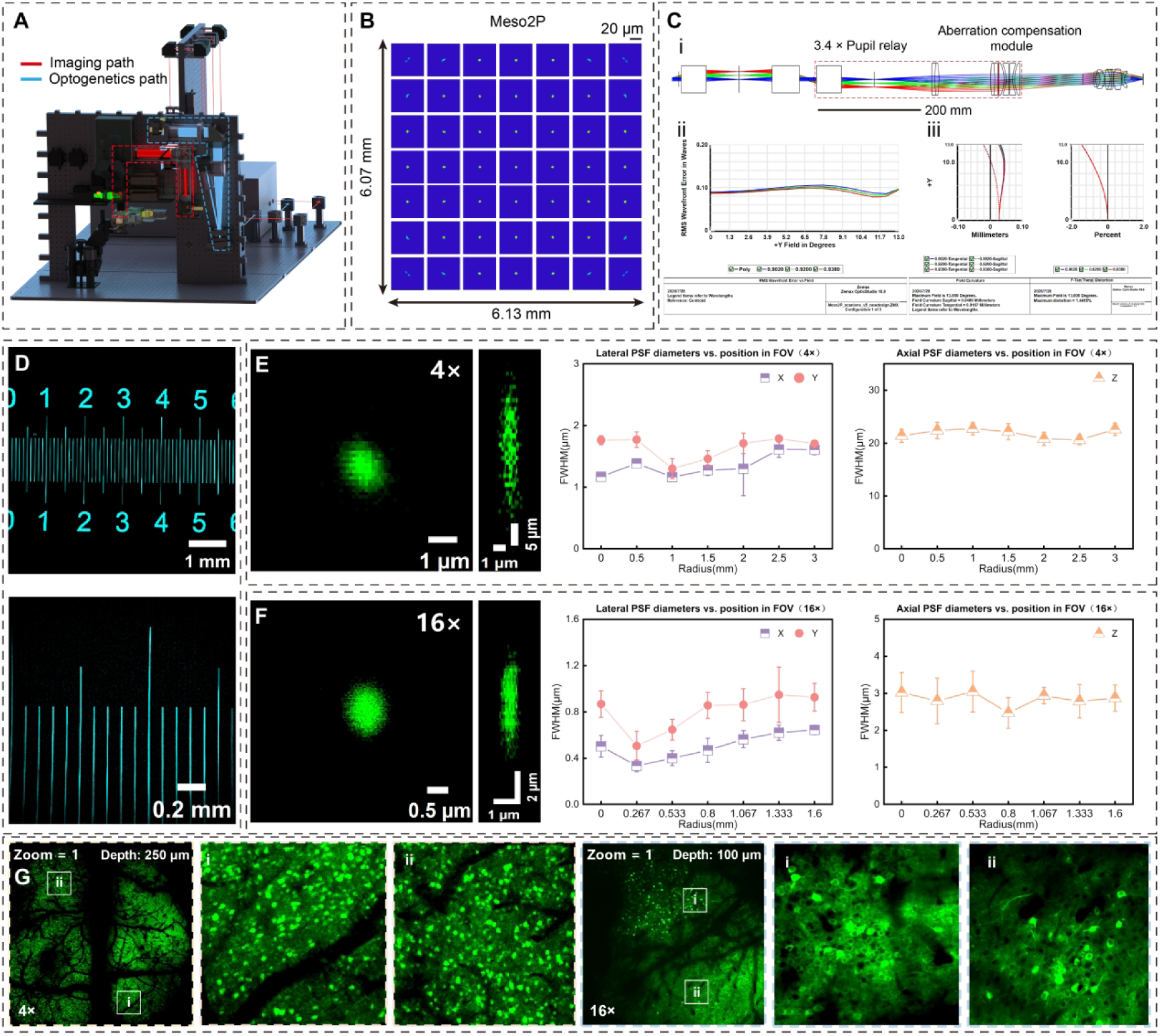
Modular design schematic and performance evaluation of Meso2P imaging, related to Figures 1 and 2 and Methods. **(A)** SolidWorks-rendered optical path geometry (excitation in blue, stimulation in orange). **(B)** Field-dependent aberration quantified by point spread function (PSF) profiles across the entire FOV. **(C)** Optical simulation performance of Meso2P transverse scanning module. (i) Zemax optical path diagram. (ii) Field-dependent RMS wavefront aberration (0°: FOV center; 13°: 3 mm off-axis). (iii) Field curvature and distortion across entire FOV. **(D)** FOV calibration using fluorescent-etched scale bars. **(E)** Acquired with a 4× objective. Left: representative point spread functions (PSFs) of the Meso2P system in the x-y and x-z planes, measured using 0.2 μm fluorescent beads. Right: lateral and axial full width at half maximum (FWHM) of the PSF as a function of radial position across the field of view (FOV). Each plotted point represents mean ± s.d. from n = 5 beads. **(F)** Same as (E), but for 16× objective. **(G)** Imaging demonstration with zoomed-in views of two regions (white boxes) obtained with 4× and 16× objectives, respectively.

**Extended Data Fig. 2.**
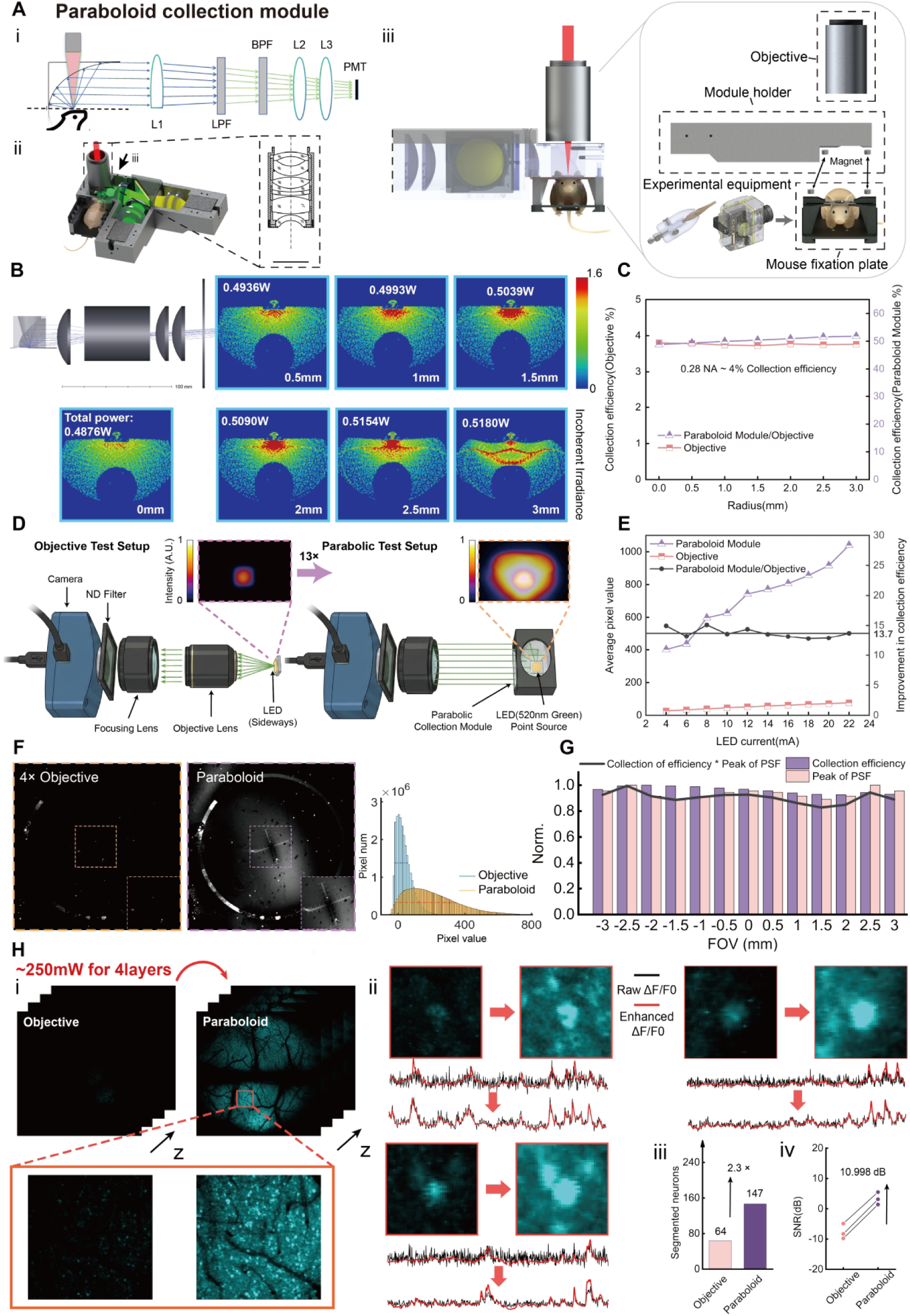
Parabolic collection module: optical layout, structural design, and performance evaluation. **(A)** Optical principle and 3D model of the parabolic collection module. (i) Optical design of the parabolic collection module. LPF, low-pass filter; BPF, band-pass filter. (ii) SolidWorks-rendered model (inset: custom 4× objective in dashed box). (iii) Exploded view of the model in (ii), showing three components: objective, module holder, and mouse fixation plate. The module holder and mouse fixation plate are connected via magnets. Experimental equipment is positioned either in front of or laterally to the mouse. **(B)** Collection efficiency simulation of the parabolic collection module in Zemax non-sequential mode. A standard point light source with a fixed power of 1 W is used, with a cone angle of 90° to restrict emission to the upper hemisphere. Heatmaps show ray tracing distribution on the PMT plane and the total collected optical power when the light source is placed at field radii from 0 to 3 mm. **(C)** Simulated collection efficiency comparison between a detector with NA = 0.28 and the parabolic collection module, as a function of point source position (0-3 mm field radius). The right y-axis scale corresponds to 13× the left y-axis scale. **(D)** Optical setup for relative radiometric measurement of collection gain. Left: Radiation measurement path through the objective. From left to right: camera, 2400 ND filter, focusing lens, objective, and point light source placed laterally at the focal point of the objective. Right: Same configuration but with the parabolic collection module; the point source is placed horizontally at the focal point of the parabolic reflector. **(E)** Relative radiometric comparison of the detector signals collected through the objective and paraboloid paths as a function of increasing source intensity (corresponding to panel D). Left y-axis: absolute mean pixel value of acquired images. Right y-axis: ratio of collection efficiency (parabolic module / objective). **(F)** Imaging comparison: objective-only versus parabolic module collection, using a fluorescent-etched scale bar sample. Pixel intensity distributions are shown for both conditions. **(G)** Field uniformity analysis (± 3 mm FOV). Plotted metrics include PSF peak intensities and collection efficiencies across the field of view. **(H)** Paraboloid collection module enhances neural imaging versus conventional objective. (i) Image quality comparison with a zoom-in region. (ii) Comparison of image quality and signal traces in single neurons (white arrows: example neurons; black/red traces, raw/NeuroPixelAI-enhanced signals). (iii) Segmentation improvement of neurons in (i). (iv) SNR enhancement of temporal signals from three marked neurons in (ii).

**Extended Data Fig. 3.**
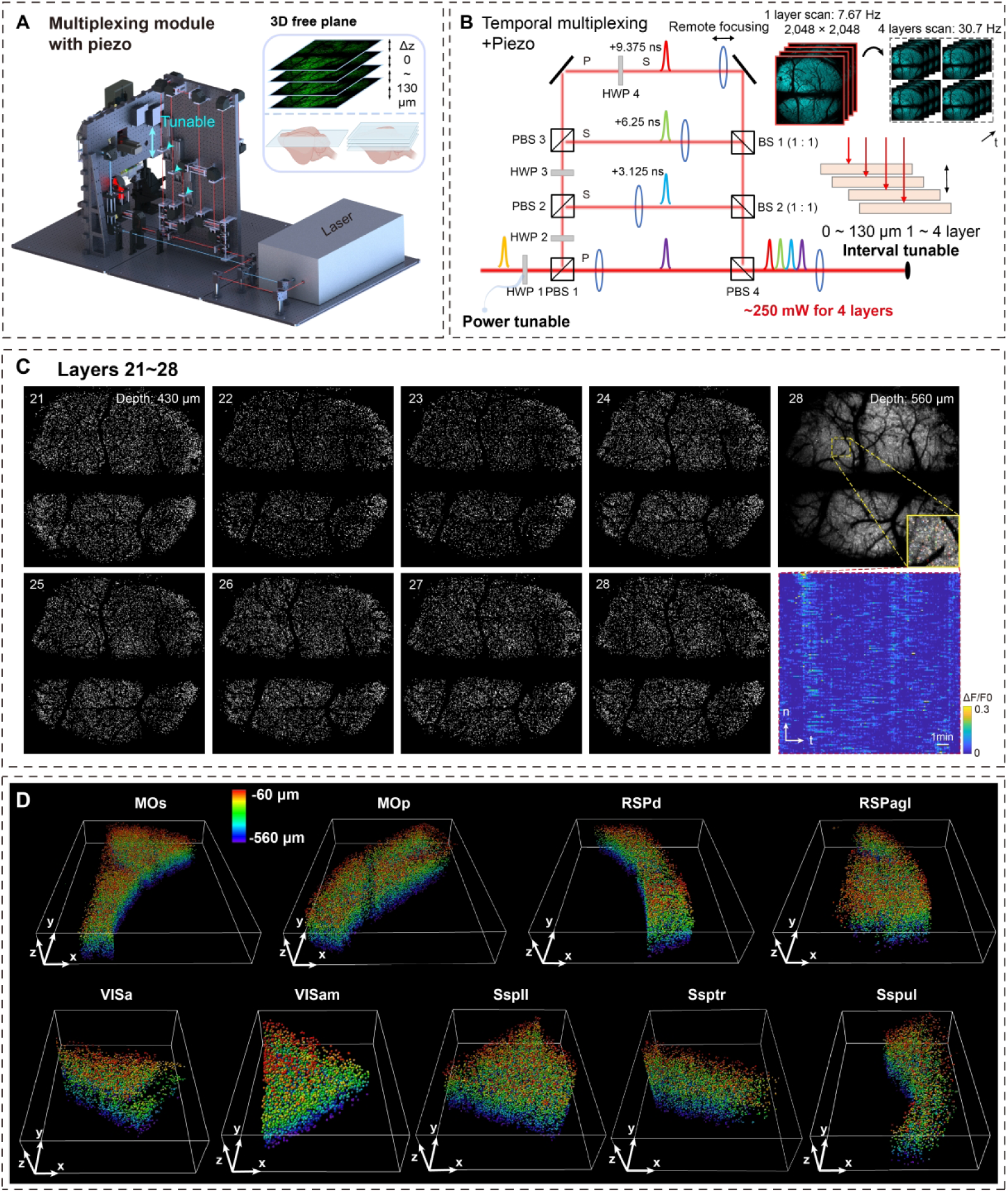
Schematic implementation and evaluation of diverse and flexible volumetric imaging using Meso2P, related to Figure 2 and Methods. **(A)** 3D SolidWorks schematic of the dual-extension module for volumetric imaging. **(B)** Dual-extension module for volumetric imaging. BS: 50/50 beam splitter; PBS: polarizing beam splitter; HWP: rotatable half-wave plate. All lenses are mounted on programmable stages for focal adjustment. **(C)** Example 3D segmentation masks across 28 imaging planes (layers 21-28; 430-560 μm below the surface). Right, example image at deepest layer (28) with zoomed-in neuronal regions (yellow boxes) and corresponding temporal signals. **(D)** 3D rendering of extracted neurons across nine representative brain regions (28 layers total). Neurons are colored by depth.

**Extended Data Fig. 4.**
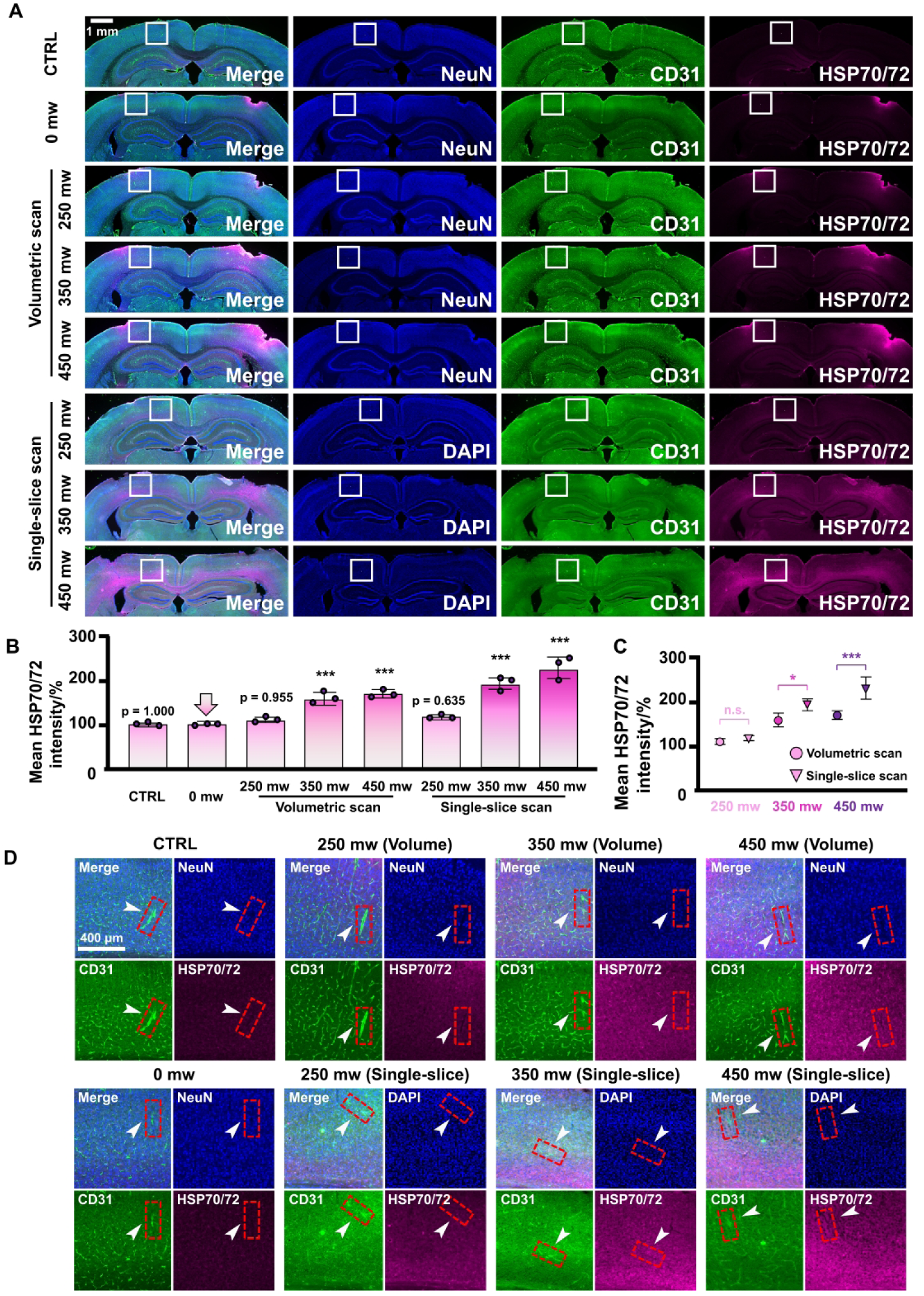
Verification of Meso2P light damage under different power and shooting modes. **(A)** Immunofluorescence comparison of light damage effects under different power and different shooting modes. **(B)** Comparison of mean HSP70/72 intensity under different power and imaging modes (n = 3 per group). \*\*\**p* < 0.001; Tukey test. **(C)** Comparison of mean HSP70/72 intensity under different imaging modes under the same power (n = 3 per group). n.s., no significance; \**p* < 0.05; \*\*\**p* < 0.001; Tukey test. **(D)** Magnified view of the white rectangular region in (A), showing HSP70/72 signal relative to CD31-positive vasculature and NeuN-positive cells. HSP70/72 signal was predominantly nonvascular and parenchymal; cellular subtype was not assigned because GFAP was not included and proximity does not establish colocalization.

**Extended Data Fig. 5.**
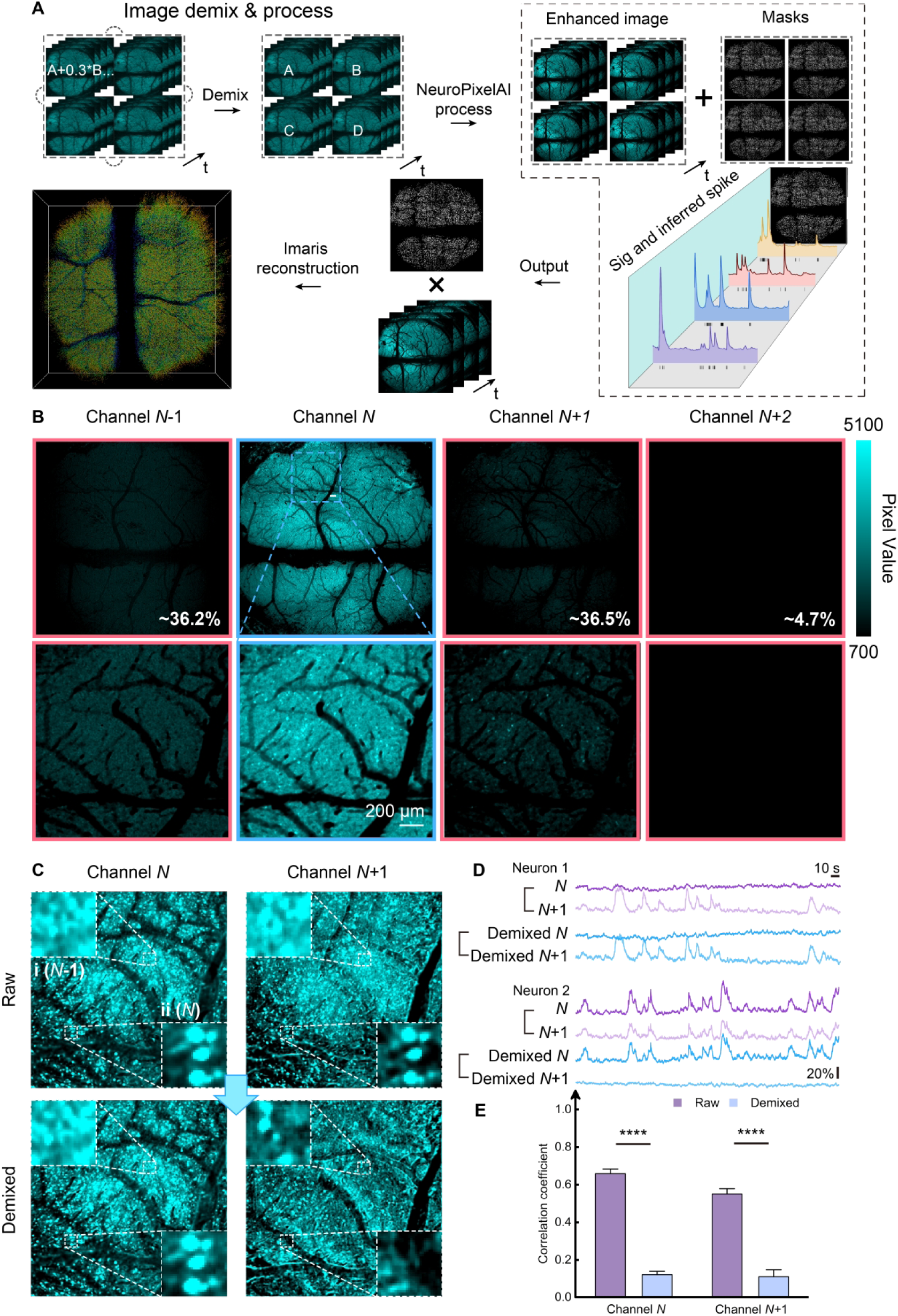
Temporal crosstalk and signal demixing across channels. **(A)** Data processing pipeline for large-volume 3D imaging on the basis of NeuroPixelAI. Key steps: Four-layer temporal crosstalk reduction by offline demixing, neuron information extraction via NeuroPixelAI, mask generation and volumetric reconstruction in Imaris. **(B)** Static crosstalk analysis for a 3.125-ns bin width. Example shows a single frame (Channel N) and bleed-through into the adjacent channels. Crosstalk ratios are calculated as percentages normalized to Channel N. **(C)** Static crosstalk between adjacent channels before and after demixing. Standard deviation projection frames are shown. Dashed boxes highlight example neuronal regions. **(D)** Neuronal signal crosstalk between adjacent channels before and after demixing, showing two example neuronal signals. **(E)** Statistical analysis of crosstalk correlation coefficients between neuronal signals before and after demixing, comparing signals from Channel N and Channel N+1 (n = 701 neurons for Channel N, n = 167 for Channel N+1; two-sided paired t-tests, \*\*\*\**p* < 0.0001). Neuronal masks were defined separately in Channel N (n = 701) and Channel N+1 (n = 167). For each mask, signals were extracted from both channels and the inter-channel correlation was calculated before and after demixing. Because the before- and after-demixing values were obtained from the same mask, comparisons were performed separately within each mask set using two-sided paired t-tests; the two mask sets were not compared with each other.

**Extended Data Fig. 6.**
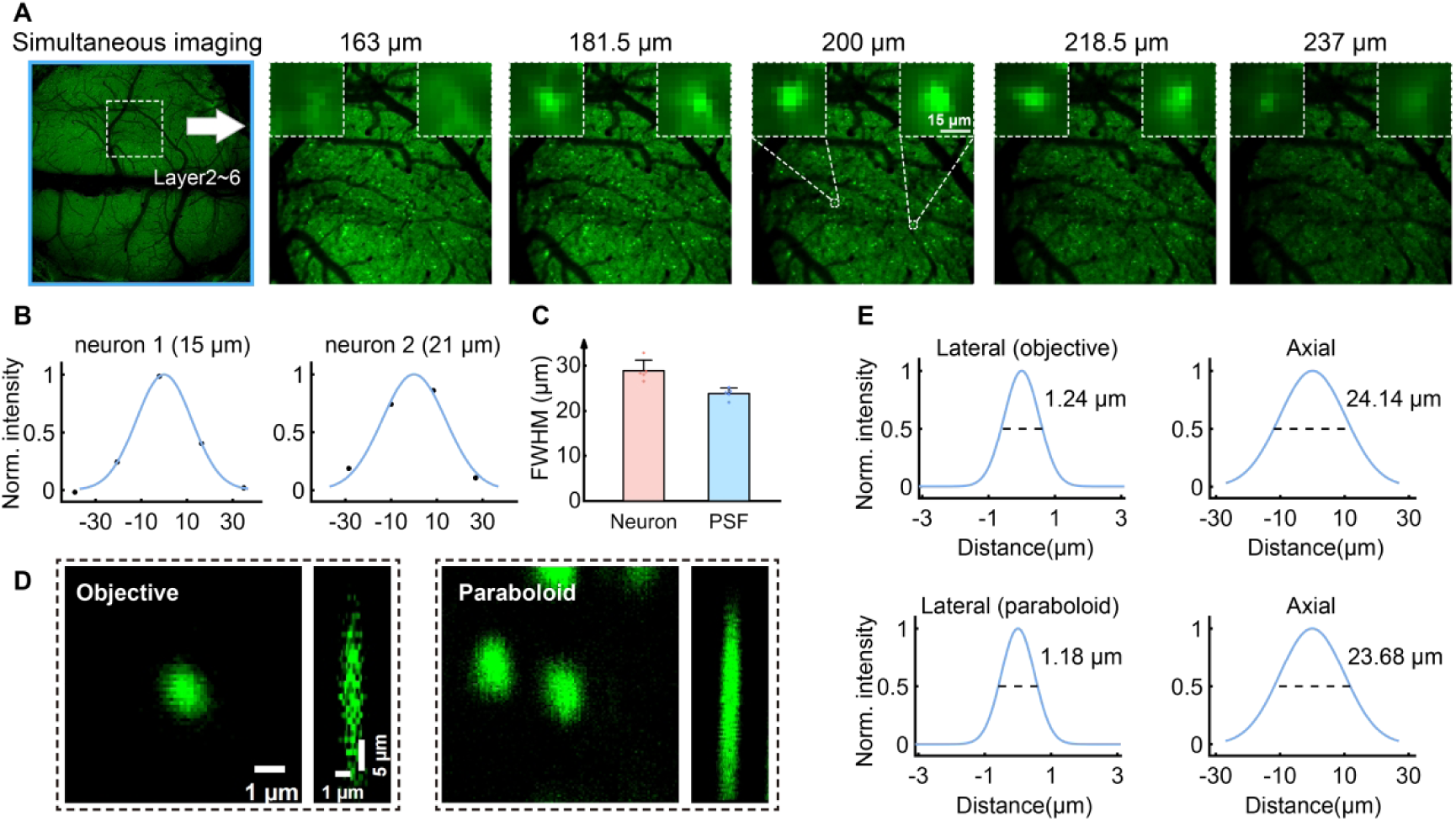
Side-by-side display of neurons in axially adjacent imaging planes. **(A)** Example single-frame images from simultaneous imaging of five axial planes spanning depths from 163 µm to 237 µm, with an inter-plane spacing of 18.5 µm. Dashed boxes indicate magnified views of the same neuronal region across these planes. **(B)** Gaussian fits to the axial fluorescence intensity profiles of example neurons with varying sizes. **(C)** Axial FWHM values of the imaged neurons (n = 5 neurons) and estimated optical PSF FWHM values after accounting for soma size. **(D)** Representative PSFs in the x-y and x-z planes, measured with 0.2-µm fluorescent beads under two collection configurations. Left, objective-only collection; right, paraboloid collection. **(E)** Gaussian fits to the lateral and axial intensity profiles of the PSFs shown in (D).

**Extended Data Fig. 7.**
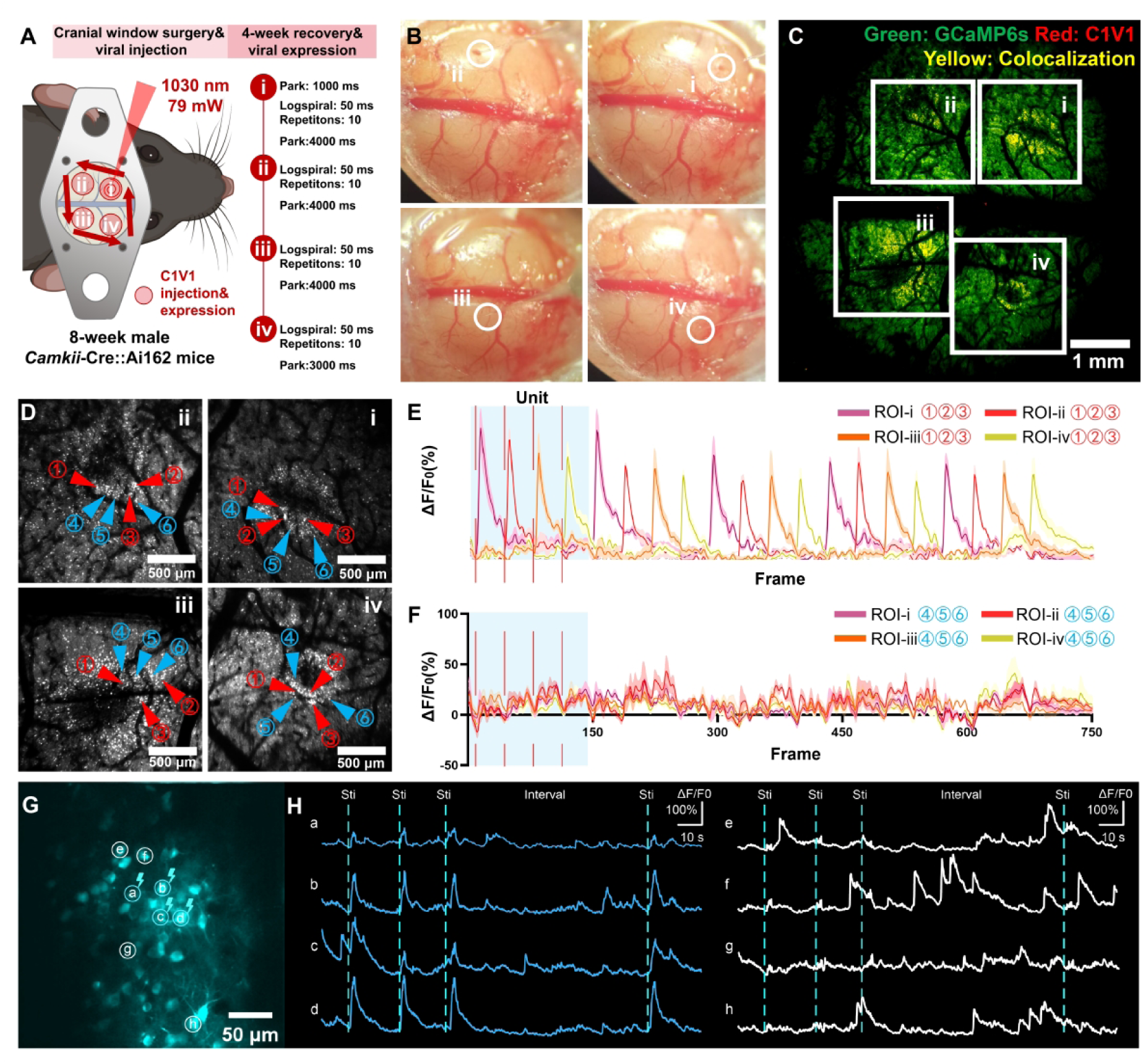
Validation of the optogenetic stimulation module. **(A)** Schematic of the optogenetic stimulation experiment. **(B)** C1V1 viral injection sites. **(C)** C1V1 expression. **(D)** Schematic of optogenetic stimulation. Red arrows indicate photostimulation target sites, and blue arrows indicate adjacent control cells. **(E)** Calcium signals from stimulated target sites during optogenetic stimulation. **(F)** Calcium signals from adjacent control cells during optogenetic stimulation. **(G)** Proof-of-concept two-photon photostimulation in layer 2/3 of the retrosplenial cortex. Four neurons (white circles) were simultaneously targeted for stimulation. **(H)** Left, calcium transients from the four photostimulated neurons showing the responses on all individual trials. Right, simultaneously recorded activity of four non-stimulated neurons.

**Extended Data Fig. 8.**
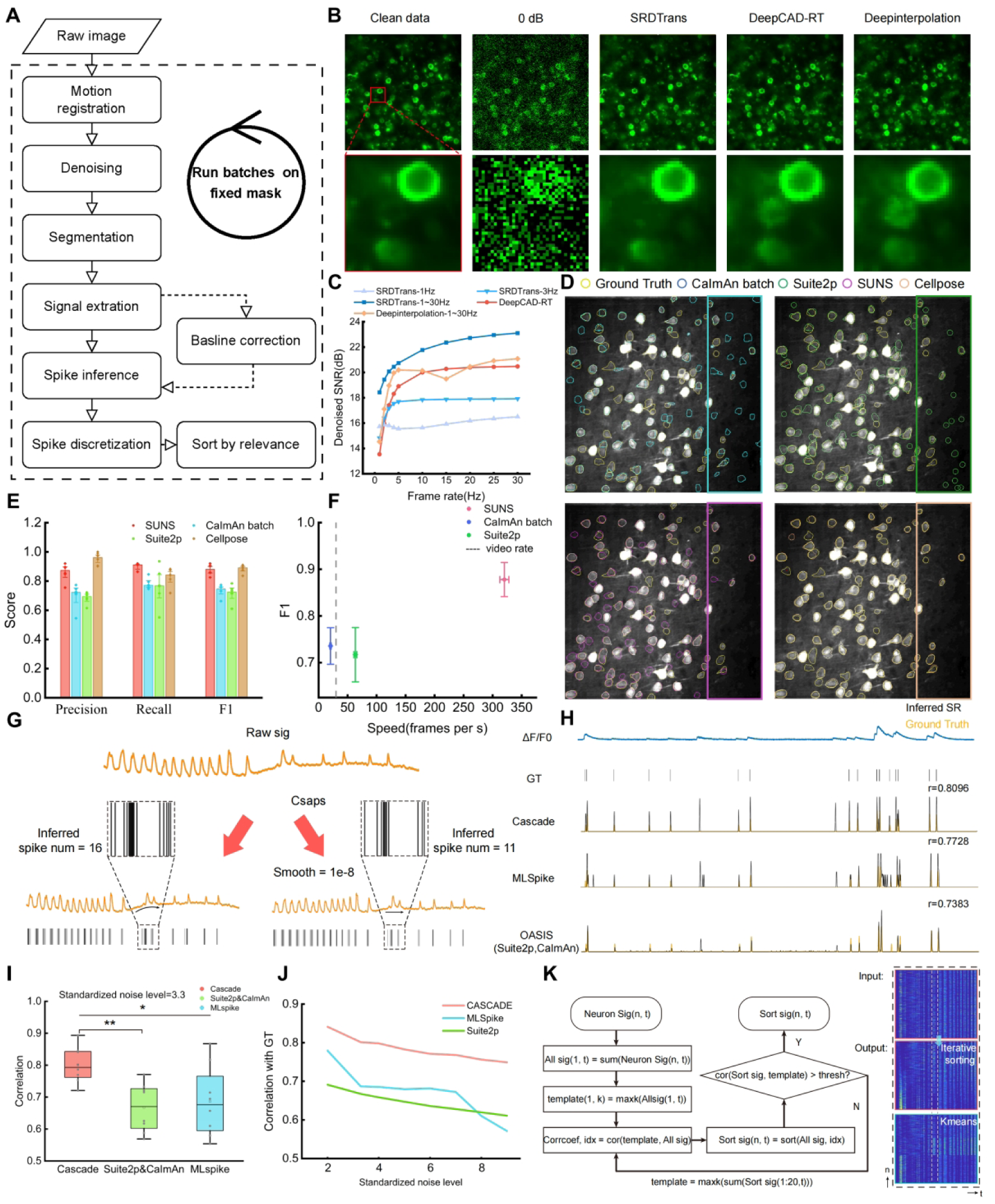
Algorithm benchmarking and implementation schematics for NeuroPixelAI, related to Figure 3 and Methods. **(A)** Schematic workflow of the standard NeuroPixelAI processing pipeline. Key steps include motion registration, denoising, segmentation, signal extraction, baseline correction, spike inference, spike discretization, and iterative sorting. Batch processing of selected steps is enabled using predefined processing masks. **(B)** Performance comparison of denoising algorithms on noisy data (0 dB SNR). Bottom panels show zoomed-in views of example neurons. **(C)** Denoising algorithm performance across varying imaging frame rates (1–30 Hz). SRDTrans was trained on datasets at each specific frame rate; DeepCAD-RT utilized its published large pretrained model. **(D)** Comparison of segmentation algorithm performance. Boxed regions highlight areas exhibiting significant differences. **(E)** Bar plot showing F1 scores of segmentation algorithms. Data derived from two-photon and three-photon imaging at depths of 100–1000 µm (n = 5 mice). **(F)** Comparison of processing speed versus F1 score for segmentation algorithms. **(G)** Schematic and efficacy of baseline correction using a cubic spline interpolation algorithm. Correction (right) reduces erroneous spike inference caused by baseline drift. **(H)** Performance comparison of spike inference algorithms. Predicted spikes (black) versus ground truth (yellow). **(I)** Correlation between inferred and ground-truth action potentials for spike inference algorithms across standardized noise levels. Datasets: Simultaneous electrophysiology and calcium imaging recordings from single neurons (GENIE datasets; n = 8 neurons). **(J)** Box plot statistics of inference correlation coefficients for all algorithms at a standardized noise level of 3.3. **(K)** Workflow and output of the label-free iterative sorting algorithm. Gray dashed lines indicate clear sorting results.

**Extended Data Fig. 9.**
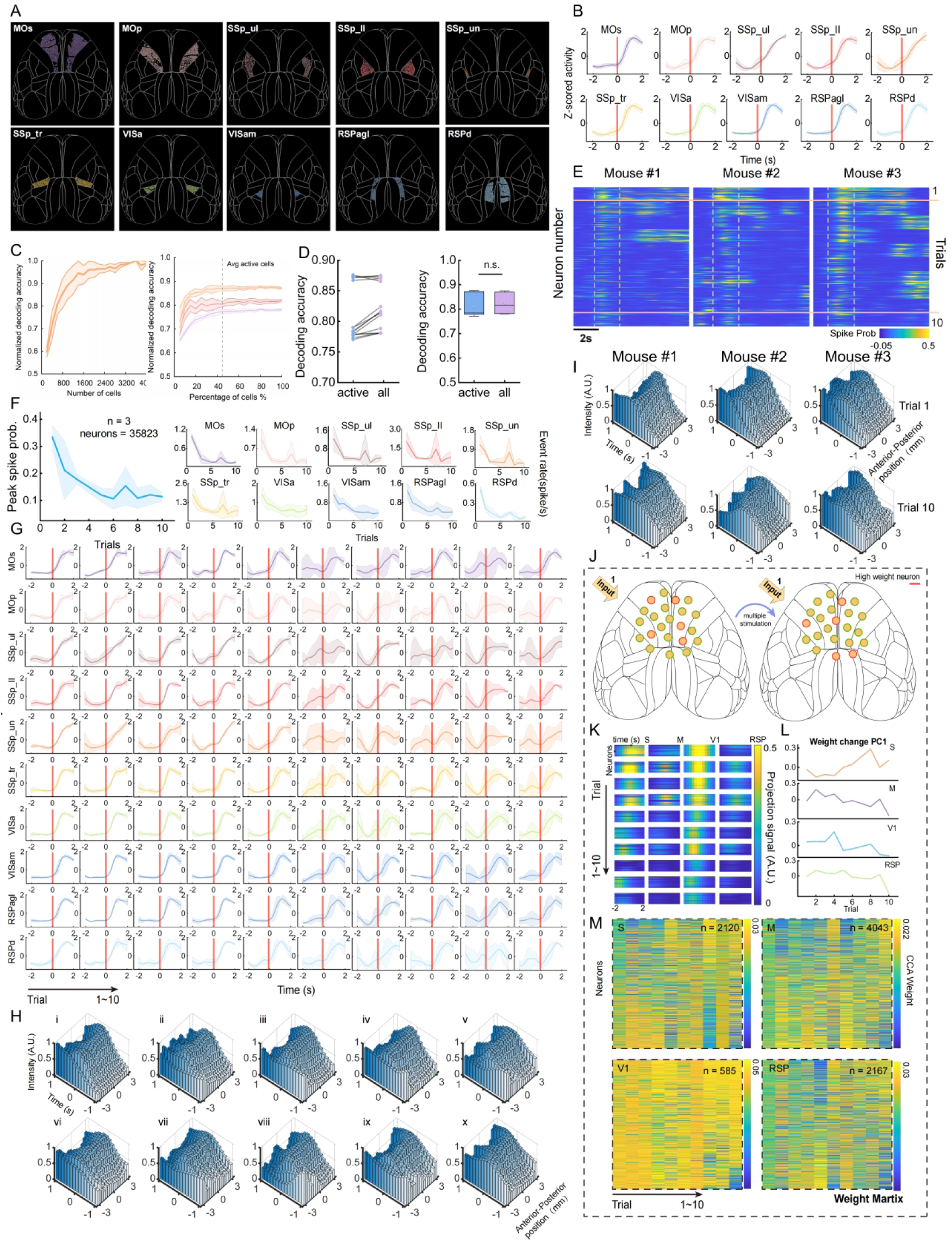
Large-scale population neural response dynamics during repetitive whisker stimulation, related to Figure 4 and Methods. **(A)** Cortical distribution of segmented stimulus-responsive neurons. Colors denote corresponding brain regions. **(B)** Average neural activity Z-scores across 10 trials for each brain region. Stimulus onset is indicated by the red vertical line. **(C)** Decoding performance as a function of neuronal population size. Left, performance versus absolute number of neurons. Right, performance versus the percentage of extracted neurons. Data are shown for three mice. The dashed line indicates the mean proportion of active neurons across mice, mean ± SD. **(D)** Comparisons of decoding accuracy using active neurons versus all neurons (n = 3 mice), 5 trials per mouse. **(E)** Calcium signal heatmaps across ten trials for three mice. Gray dashed lines demarcate the 4 s peri-stimulus period. Pink solid lines separate signals from the first and last trials. The heatmaps reveal concurrent changes in response intensity and neuronal loading weights across trials. **(F)** Left, trial-by-trial changes of peak spiking probability across 10 trials in three mice. Right, changes in calcium event rate (mean ± SD) within a 2-s window surrounding stimulus onset for individual recorded brain regions. **(G)** Trial-dependent dynamics of regional brain activity. Individual panels display neural activity Z-scores per brain region in single trials. Stimulus onset is marked by red vertical lines. **(H)** Bar plots of normalized spike probability across 10 trials, averaged across three mice. **(I)** Normalized neuronal spike probabilities over time and position for trials 1 and 10. Bar plots show the normalized spike probability at each time point and anterior-posterior bin for the first (Trial 1) and last (Trial 10) trials. Small neuronal populations in the posterior cortical areas of all three mice exhibited a significant temporal shift in peak activity in Trial 10 compared to Trial 1. **(J)** Schematic of structural reorganization in neural networks. Illustrates trial-dependent changes in single-neuron loading weights despite identical sensory inputs. **(K)** The projected activity of each area in the CCA dimensions identified across 10 trials. Population activity in S1 and V1 cortices exhibits dominant contributions to the stimulus-evoked cofluctuation mode, with trial-dependent modulation in response magnitude and phase shifts. **(L)** PC1 derived from PCA of neuronal loading weight matrices. **(M)** Heatmap of neuronal loading weights across trials. Depicts trial-dependent reorganization of loading weights within the network. **(K-M)** Representative data from one mouse.

**Extended Data Fig. 10.**
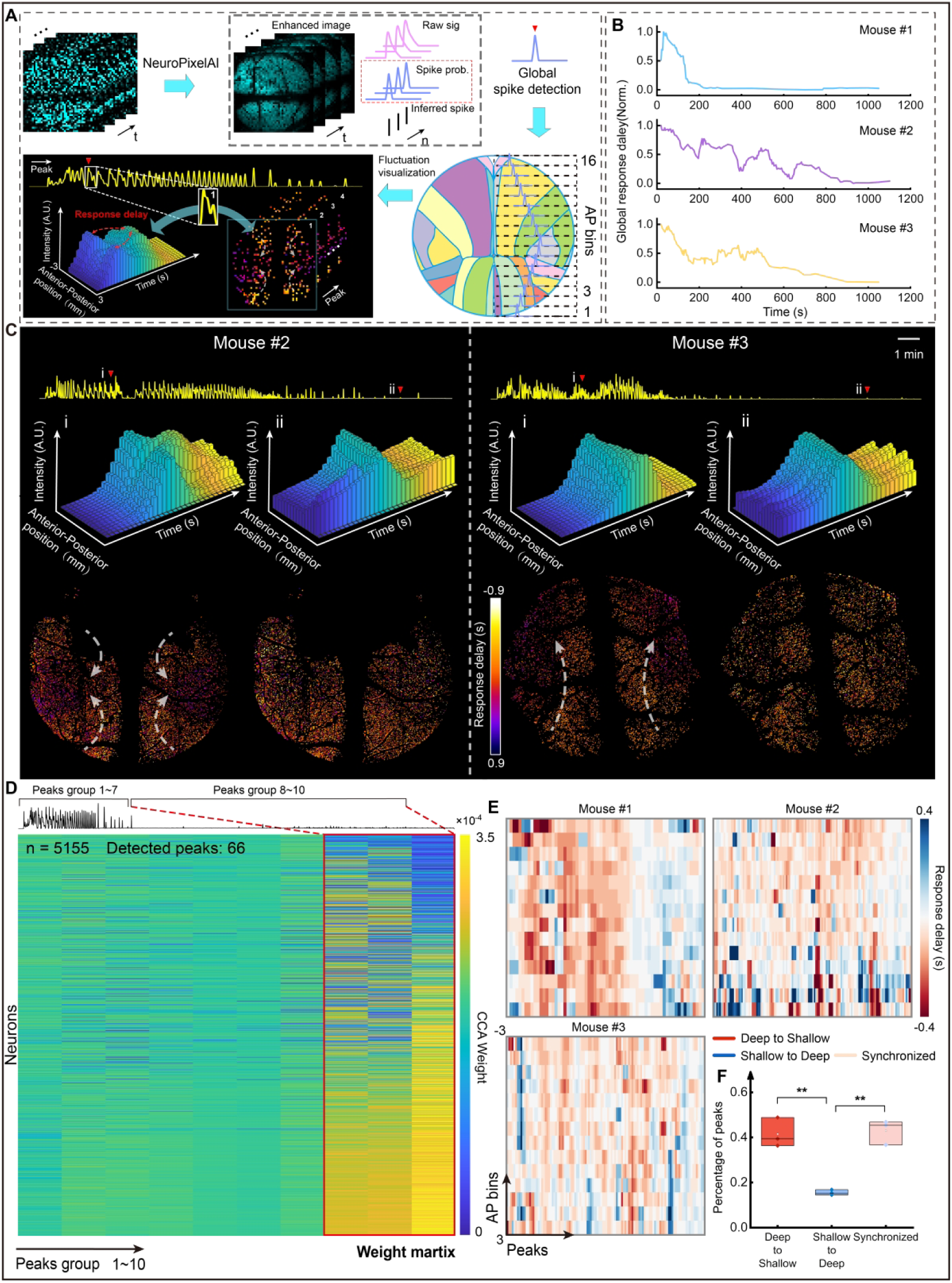
Schematic representation and assessment of intra- and interlaminar signaling dynamics during anesthesia induction using Meso2P microscopy, related to Figure 5 and Methods. **(A)** Data processing pipeline for anesthetized imaging experiments. Raw image data were processed by NeuroPixelAI to extract single-neuron spike probabilities. Global spike probability traces underwent peak detection to identify multiple “synchronous” event timings. For each synchronized event, fluctuation delays were quantified at both individual anterior-posterior (AP) bins and single-neuron levels, followed by spatiotemporal visualization. **(B)** Normalized global response delays per mouse. Standard deviation (SD) of peak times in spike probabilities across 16 anterior-posterior (AP) bins was computed during anesthesia progression. This reveals a gradual transition from spatially propagating delays (indicating asynchronous activity) to uniform synchrony across cortical regions within individual synchronous events. **(C)** Schematics of synchronized signal propagation direction in two additional mice. **(D)** Heatmap of neuronal CCA loading weights across synchronous events in an example mouse. Synchronous events were divided into 10 consecutive time bins. For each bin, cross-bin CCA identified stable activity modes during anesthesia induction. Late-phase bins exhibited significant reorganization of loading weights, concomitant with gradual uniform synchronization of cortical activity. **(E)** Heatmap of synchronized signal propagation direction between deep and shallow cortical layers during anesthesia induction. This indicates the predominant direction of signal flow during each synchronous event within individual AP bins. Red, Deep-to-shallow propagation; Blue, Shallow-to-deep propagation. **(F)** Boxplot of signal propagation direction proportions between deep and shallow cortical layers during anesthesia induction (n = 3 mice), *\*\*p* < 0.01, Tukey test.

**Extended Data Fig. 11.**
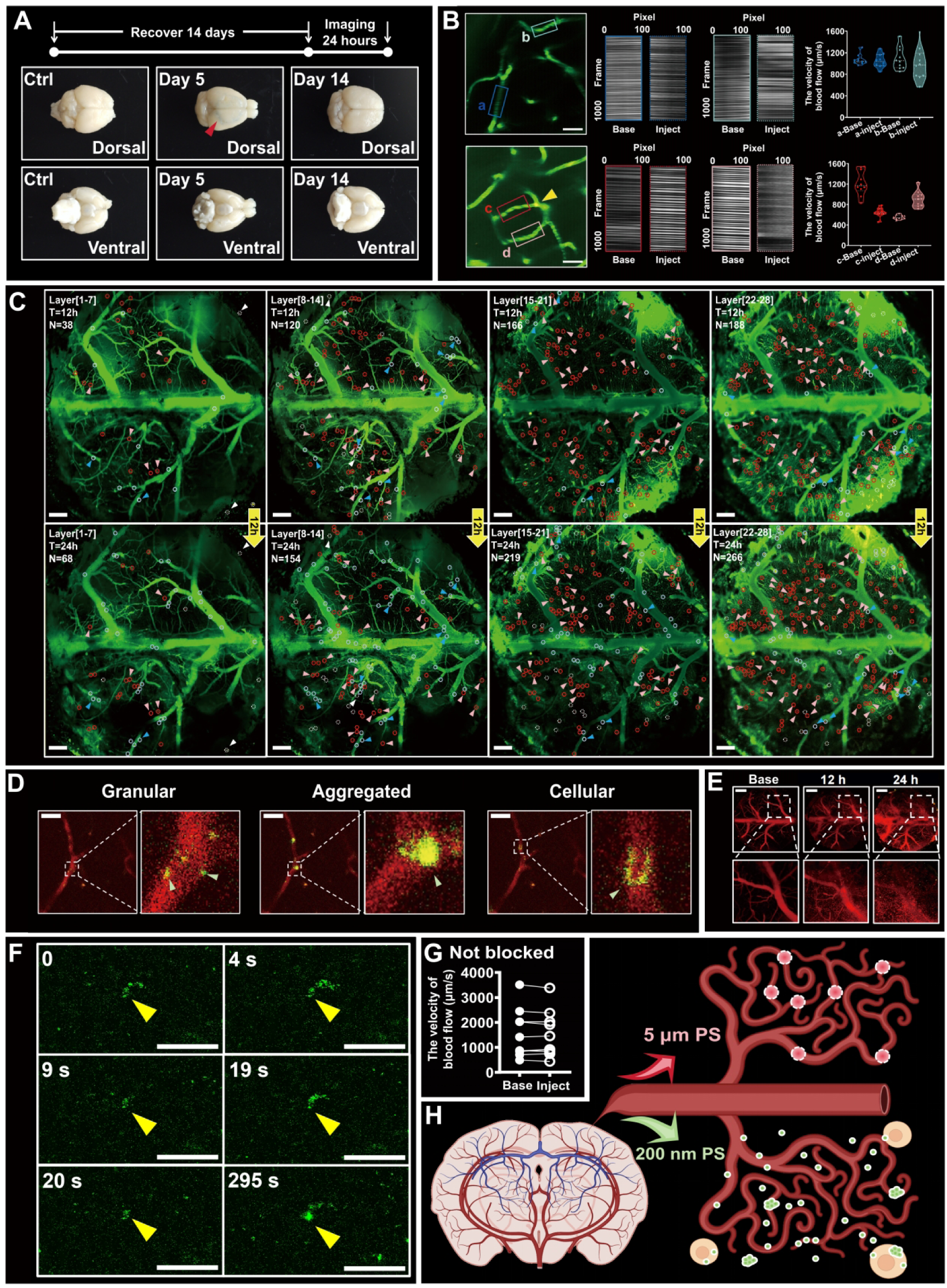
Supplementary data for in vivo MNP imaging. **(A)** Blood-brain barrier (BBB) integrity assay to determine optimal MNP imaging window; compromised BBB (red arrows). **(B)** Comparison of blood flow velocity before and 12–24 h after MP injection: vessels (green), MPs (yellow), unoccluded vessels (a, b), occluded vessel (c), adjacent vessel (d). **(C)** Cumulative distribution of MPs across vascular layers 12 h post-injection: vessels (green), MPs (red), co-localization (yellow). MPs occluding vessels < 30 µm (red circles), > 30 µm (blue circles), extravascular MPs (white dashed circles), unresolved MPs in vessels < 30 µm (pink arrows) and > 30 µm (blue arrows). **(D)** Structural morphology of NPs in blood vessels: vessels (red), NPs (green), co-localization (yellow). **(E)** Time-dependent accumulation of NP-laden cells in the uppermost cortical layer following NP injection: vessels (red). **(F)** NP aggregation and clearance dynamics in blood vessels: NPs (green), aggregation foci (yellow arrows). **(G)** Comparison of blood flow velocity in vessels treated with NPs. **(H)** Schematic illustration of differential brain distribution patterns between 5-µm MPs and 200-nm NPs. *Scale bars*: 500 µm (C, E); 100 µm (D, F). ***See also* Supplementary Videos 7-11.**

**Extended Data Fig. 12.**
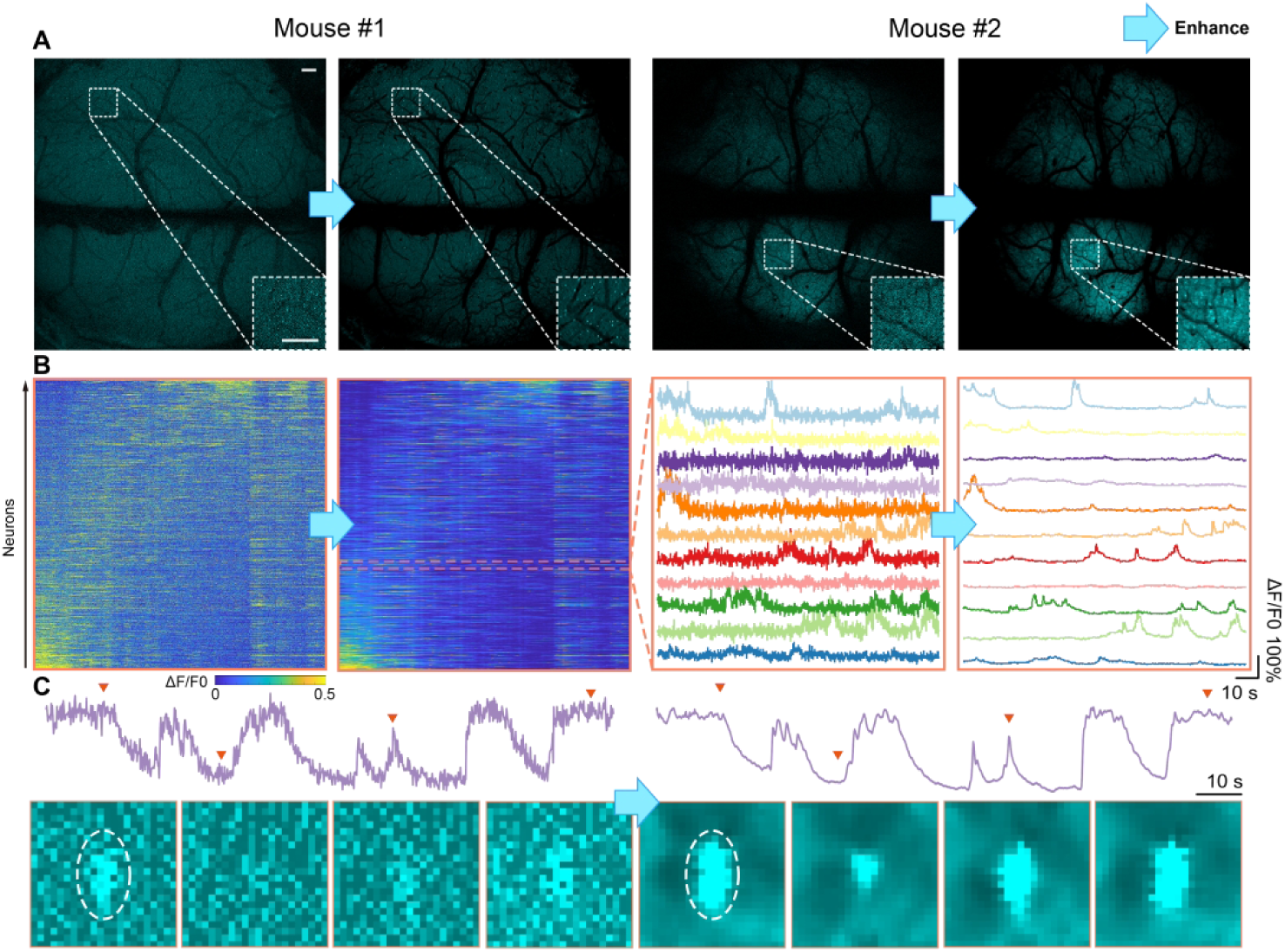
Comparison of data before and after NeuroPixelAI enhancement. **(A)** Comparison of raw acquired images and NeuroPixelAI-enhanced images from two example mice. Blue arrows indicate the enhancement step, and dashed boxes outline magnified regions. Scale bar: 300 μm. **(B)** Comparison of neuronal calcium traces extracted from the image data in (A) before and after enhancement. **(C)** Comparison of an example neuron before and after enhancement. Orange triangles mark representative frames, which are displayed below the calcium trace.

## Supplementary Tables

**Supplementary Table 1:** Summary of Meso2P operating specifications and demonstrated capabilities.

| Parameters | Meso2P |
| --- | --- |
| Maximum field of view | 6 × 6 mm<sup>2</sup> |
| Excitation objective | 4× objective, NA = 0.28 |
| Equivalent collection NA | 0.87 (solid-angle-equivalent collection metric) |
| Lateral resolution (FWHM) | 1.2-1.8 μm |
| Axial resolution (FWHM) | 21-24 μm |
| Maximum frame size | 4096 × 4096 pixels at 3.83 Hz |
| High-speed imaging | 2048 × 2048 pixels at 7.67 Hz |
| Volumetric imaging | 6 × 6 × 0.5 mm<sup>3</sup> at 1.1 Hz |
| Two-photon photostimulation | Yes |

**Supplementary Table 2:** Comparison of reported volumetric imaging parameters for Meso2P and representative large-FOV two-photon platforms.

| System | Volume size<br>(mm<sup>3</sup>) | Sampling format<br>(voxel<sup>3</sup>) | Volume<br>rate (Hz) | Axial planes<br>per volume | Voxel rate<br>(Mvoxel/s) |
| --- | --- | --- | --- | --- | --- |
| Meso2P | 6×6×0.5 | 2,048×2,048×28 | 1.1 | 28 | 129.18 |
| Light beads | 5.4×6×0.5 | 1,080×1,200×30 | 2.2 | 30 | 85.54 |
| DEEPscope | 3.23×3.23×0.08 | 1,030×515×5 | 2.2 | 5 | 5.83 |

**Supplementary Table 3:**
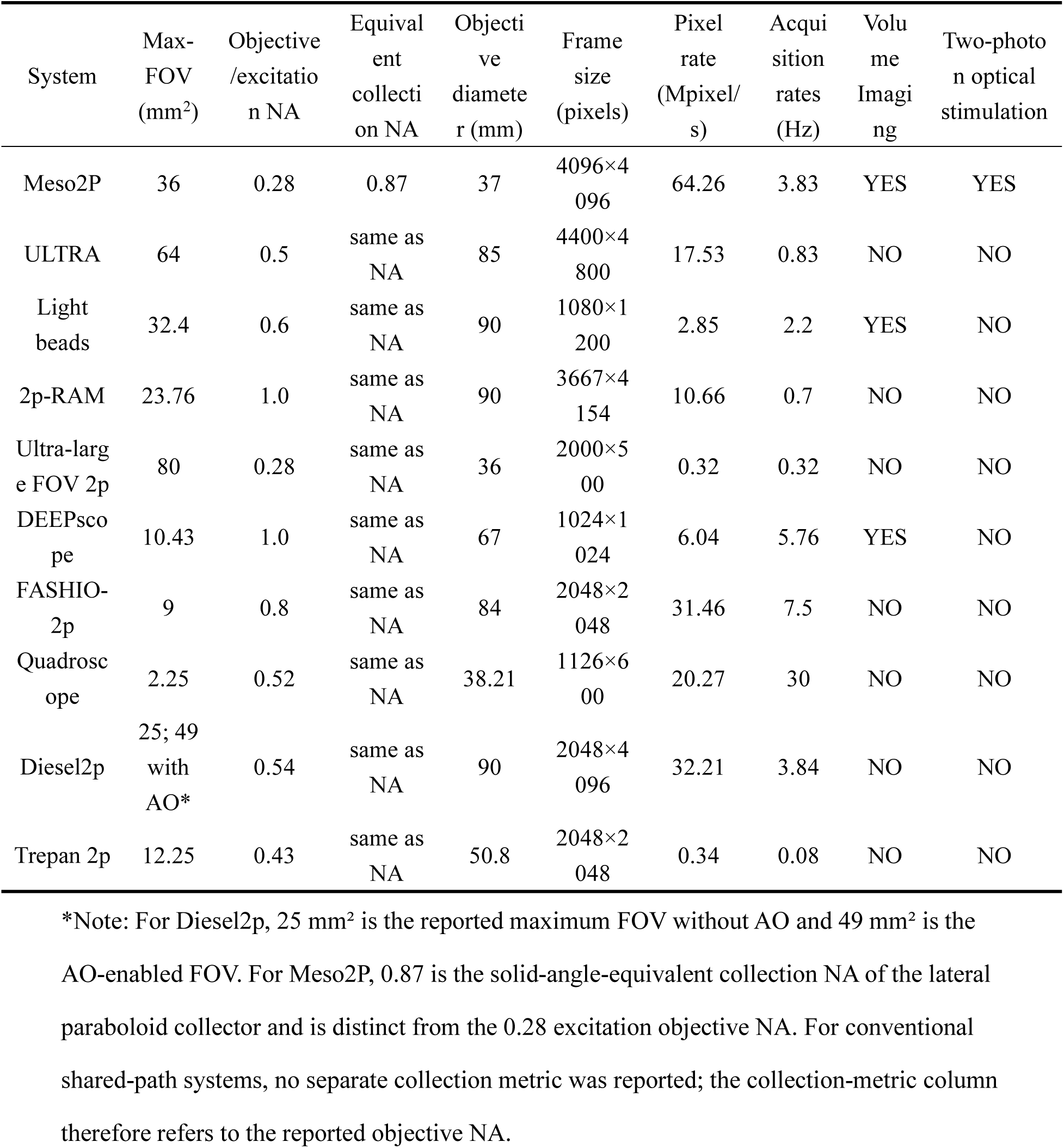
Comparison of optical and acquisition parameters and demonstrated capabilities for Meso2P and representative large-FOV two-photon platforms.

| System | Max-FOV (mm<sup>2</sup>) | Objective/excitation NA | Equivalent collection NA | Objective diameter (mm) | Frame size (pixels) | Pixel rate (Mpixel/s) | Acquisition rates (Hz) | Volume Imaging | Two-photon optical stimulation |
| --- | --- | --- | --- | --- | --- | --- | --- | --- | --- |
| Meso2P | 36 | 0.28 | 0.87 | 37 | 4096×4096 | 64.26 | 3.83 | YES | YES |
| ULTRA | 64 | 0.5 | same as NA | 85 | 4400×4800 | 17.53 | 0.83 | NO | NO |
| Light beads | 32.4 | 0.6 | same as NA | 90 | 1080×1200 | 2.85 | 2.2 | YES | NO |
| 2p-RAM | 23.76 | 1.0 | same as NA | 90 | 3667×4154 | 10.66 | 0.7 | NO | NO |
| Ultra-large FOV 2p | 80 | 0.28 | same as NA | 36 | 2000×500 | 0.32 | 0.32 | NO | NO |
| DEEPscope | 10.43 | 1.0 | same as NA | 67 | 1024×1024 | 6.04 | 5.76 | YES | NO |
| FASHIO-2p | 9 | 0.8 | same as NA | 84 | 2048×2048 | 31.46 | 7.5 | NO | NO |
| Quadroscope | 2.25 | 0.52 | same as NA | 38.21 | 1126×600 | 20.27 | 30 | NO | NO |
| Diesel2p | 25; 49 with AO* | 0.54 | same as NA | 90 | 2048×4096 | 32.21 | 3.84 | NO | NO |
| Trepan 2p | 12.25 | 0.43 | same as NA | 50.8 | 2048×2048 | 0.34 | 0.08 | NO | NO |

\*Note: For Diesel2p, 25 mm<sup>2</sup> is the reported maximum FOV without AO and 49 mm<sup>2</sup> is the AO-enabled FOV. For Meso2P, 0.87 is the solid-angle-equivalent collection NA of the lateral paraboloid collector and is distinct from the 0.28 excitation objective NA. For conventional shared-path systems, no separate collection metric was reported; the collection-metric column therefore refers to the reported objective NA.

**Supplementary Table 4:** Dispersion benchmark evaluation for each material in Meso2P (Main optical path).

|  | Components | Reference | Glass | Thickness | GVD (fs<sup>2</sup>/mm) | GDD (fs<sup>2</sup>) |
| --- | --- | --- | --- | --- | --- | --- |
| Main optical path | Relaylens 1 | 1 | N-SF6HT | 5 | 157.73 | 788.65 |
|  |  |  | N-LAK22 | 8.2 | 55.14 | 452.15 |
|  |  | 2 | N-LAK22 | 8.2 | 55.14 | 452.15 |
|  |  |  | N-SF6HT | 5 | 157.73 | 788.65 |
|  | Relaylens 2 | 3 | N-SF6HT | 5 | 157.73 | 788.65 |
|  |  |  | N-LAK22 | 8.2 | 55.14 | 452.15 |
|  |  | 4 | N-LAK22 | 8.2 | 55.14 | 452.15 |
|  |  |  | N-SF6HT | 5 | 157.73 | 788.65 |
|  | Relaylens 3 | 5 | N-F2 | 37.2 | 81.10 | 3017.07 |
|  | Relaylens 4 | 6 | N-F2 | 37.2 | 81.10 | 3017.07 |
|  | Scanlens | 7 | N-F2 | 37.2 | 81.10 | 3017.07 |
|  | Tubelens | 8 | N-SF11 | 6 | 150.80 | 904.80 |
|  |  |  | N-SK2 | 8 | 51.40 | 411.23 |
|  |  | 9 | N-SK2 | 8 | 51.40 | 411.23 |
|  |  |  | N-SF11 | 6 | 150.80 | 904.80 |
|  | Compensation lens | 10 | N-BK7 | 10 | 34.09 | 340.93 |
|  |  | 11 | N-BK7 | 5.43 | 34.09 | 185.12 |
|  |  | 12 | N-BK7 | 5.43 | 34.09 | 185.12 |
|  |  | 13 | N-BK7 | 10 | 34.09 | 340.93 |
|  | Objective | 14-20 | N-BK7 | 4 | 34.09 | 136.37 |
|  |  |  | N-BK7 | 8.645 | 34.09 | 294.73 |
|  |  |  | H-FK61 | 10 | 28.70 | 287.02 |
|  |  |  | H-ZF1 | 3 | 93.83 | 281.50 |
|  |  |  | H-ZF62 | 3 | 214.96 | 644.88 |
|  |  |  | H-FK61 | 15.03 | 28.70 | 431.39 |
|  |  |  | H-ZF62 | 4.746 | 214.96 | 1020.20 |
| Total |  |  |  |  |  | 20795 |

**Supplementary Table 5:**
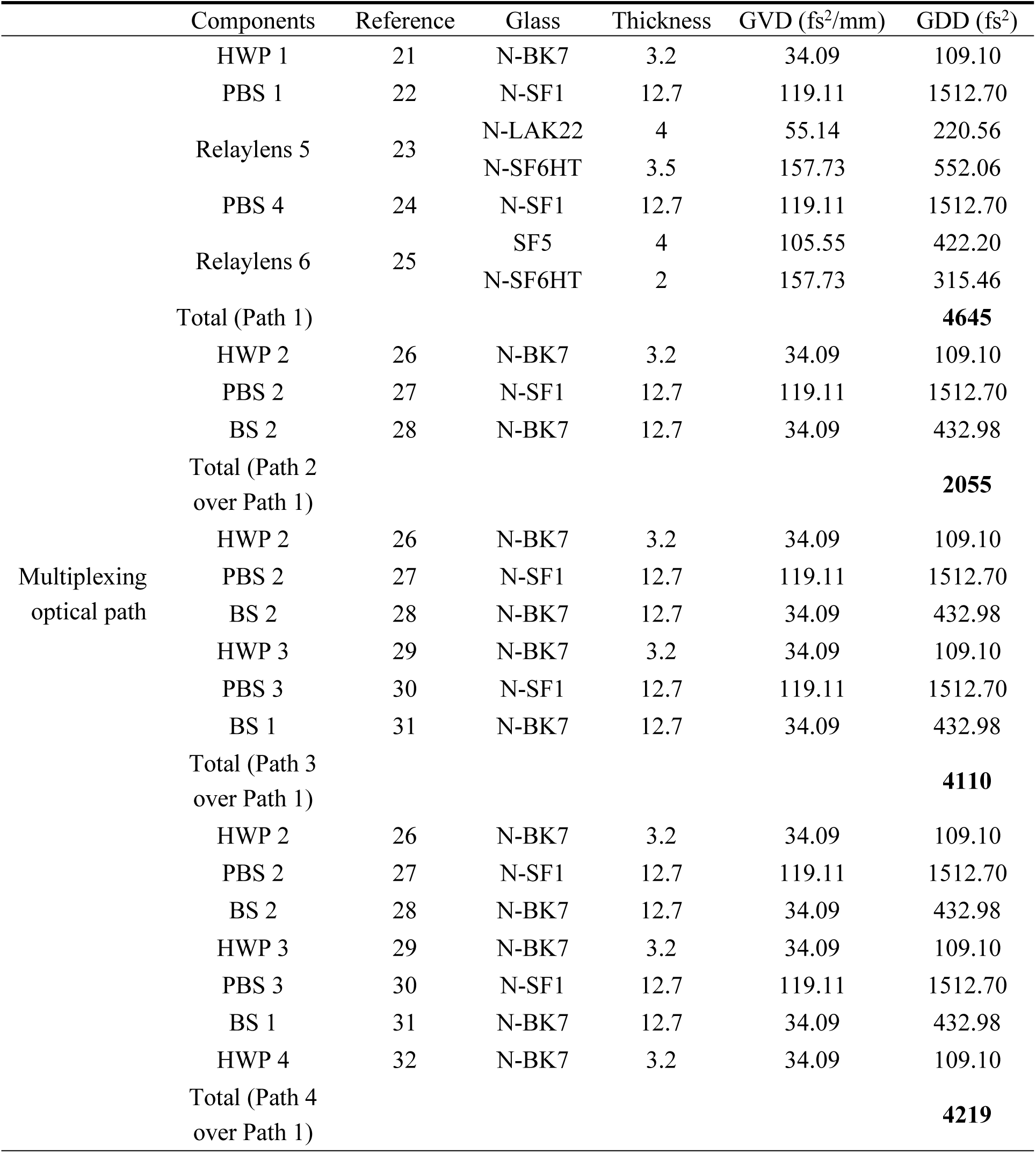
Dispersion benchmark evaluation for each material in Meso2P (Multiplexing optical path).

**Supplementary Table 6.**
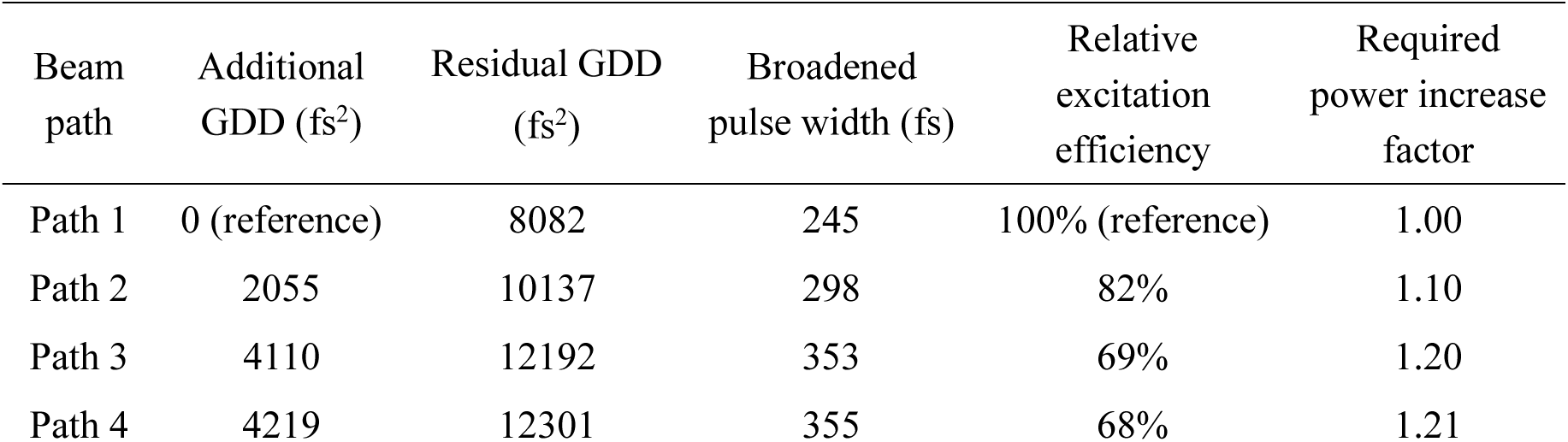
Arm-dependent GDD and estimated pulse broadening.

## Supplementary Notes

**Supplementary Note 1: Meso2P specifications and descriptive comparison with large-FOV two-photon microscopes**

Supplementary Table 1 summarizes the operating specifications and demonstrated capabilities of Meso2P. Supplementary Tables 2 and 3 provide descriptive cross-platform comparisons of volumetric imaging and optical/acquisition parameters, respectively. Because Meso2P employs separate excitation and collection optical paths, we report the collection metric separately from the objective/excitation NA, along with the field of view, frame dimensions, acquisition rate, volumetric capability, and two-photon photostimulation functionality as independent parameters, each listed on its own.

For single plane imaging, Meso2P operates at 3.83 Hz with a frame size of 4,096 × 4,096 pixels. In the volumetric configuration, it acquires 28 axial planes over a 6 × 6 × 0.5 mm³ volume at 1.1 Hz with a frame size of 2,048 × 2,048 pixels. The spatial sampling was selected to balance optical resolution and acquisition throughput. The 4,096×4,096 mode provides approximately 1.46 µm lateral sampling, comparable to the measured lateral FWHM of 1.2-1.8 µm, whereas the faster 2,048×2,048 mode provides approximately 2.93 µm sampling for soma-level imaging. The approximately 18.5 µm axial sampling interval is comparable to the measured axial FWHM of 21-24 µm. All throughput values depend on the specific operating mode and are provided as relative references for cross-comparison.

In addition, Meso2P also integrates two-photon optogenetic stimulation in the same optical core. The corresponding entries in Supplementary Tables 1 and 3 indicate demonstrated availability in the present system and are not used to construct a composite performance score.

**Supplementary Note 2: Design and beam propagation of the polarization-based spatiotemporal multiplexing module.**

The working principle of the polarization-based spatiotemporal multiplexing module and the propagation paths of the four beams are described as follows (see Extended Data Fig. 3B).

The input beam is first split by PBS 1 into P- and S-polarized components. The P-polarized component is transmitted and forms the first beam path (purple). This beam remains P-polarized and is transmitted through PBS 4 into the common output path as Channel 1.

The S-polarized component reflected by PBS 1 passes through HWP 2, which adjusts its polarization angle, and then reaches PBS 2. At PBS 2, the P-polarized component is transmitted, while the S-polarized component is reflected and forms the second beam path (blue). This blue beam remains S-polarized at the final recombination stage, passes through BS 2, and is reflected by PBS 4 into the common output path as Channel 2.

The transmitted P-polarized component then passes through HWP 3 to adjust its polarization angle and reaches PBS 3. At PBS 3, the S-polarized component is reflected and forms the third beam path (green). This green beam remains S-polarized at the final recombination stage, passes through BS 1 and BS 2, and is reflected by PBS 4 into the common output path as Channel 3.

The P-polarized component transmitted through PBS 3 constitutes the fourth beam path (red). It is then converted to S-polarization by HWP 4, passes through BS 1 and BS 2, and is reflected by PBS 4 into the common output path as Channel 4.

The optical path lengths of the four channels were set to produce relative delays of 0, 3.125, 6.250 and 9.375 ns at the focal plane. The axial separation between the corresponding focal planes can be continuously adjusted from 0 to 130 μm by translating the corresponding relay lens in the multiplexing module.

**Supplementary Note 3: Comparison of photodamage under varied laser powers and imaging modalities.**

To systematically characterize photodamage induced by the Meso2P system across different laser powers and imaging paradigms, we performed quantitative immunofluorescence assays, with the fluorescence intensity of HSP70/72 (Heat shock protein70/72) serving as the quantitative readout for photodamage. To exclude confounding effects derived from cranial window implantation surgery, two control groups were established: a naive control group (Ctrl) and a 0 mW sham surgery group that underwent identical cranial window procedures without laser irradiation.

For both multi-plane imaging enabled by the spatiotemporal multiplexing module and conventional single-plane two-photon excitation imaging, photodamage increased progressively with elevated laser power. Specifically, irradiation at 350 mW and 450 mW elicited significantly aggravated photodamage relative to the 0 mW sham group, whereas no statistically significant difference was detected at 250 mW (Extended Data Fig. 4A,B).

We further compared the power dependence of photodamage susceptibility between multi-plane and single-plane imaging modes. Under identical laser power settings, single-plane imaging induced more severe photodamage, and this discrepancy became more pronounced at higher laser powers. This phenomenon is likely attributed to the distributed allocation of total laser power across distinct focal planes during multi-plane image acquisition (Extended Data Fig. 4C).

To clarify the cellular origin of upregulated HSP70/72 upon laser exposure, we co-stained tissue sections for platelet endothelial cell adhesion molecule-1 (CD31) and neuron-specific nuclear protein (NeuN). HSP70 signals were predominantly distributed in regions surrounding NeuN and DAPI signals, with no detectable colocalization with CD31. These observations support a predominantly nonvascular, parenchymal distribution of the HSP70/72 signal. Because GFAP was not included and proximity to NeuN does not establish cellular colocalization, we do not assign the signal to a specific neuronal or glial cell type.

Collectively, these quantitative immunofluorescence analyses characterize HSP70/72-associated tissue stress across the tested laser powers and imaging modes. Under the specific acquisition and histological conditions tested, neither multi-plane nor single-plane imaging at 250 mW produced a statistically detectable increase relative to the sham control. This result does not establish universal safety or exclude other forms of phototoxicity outside the tested conditions.

**Supplementary Note 4: Characterization and correction of temporal crosstalk.**

In the Meso2P system, to expand imaging throughput and achieve high-speed multi-plane volumetric imaging, we incorporated a four-channel spatiotemporal multiplexing module. This module splits a single pulsed beam into four beamlets with a fixed temporal separation of 3.125 ns and an adjustable axial offset between their focal planes (Fig. 1C, Extended Data Fig. 3A,B). However, this 3.125-ns interval typically introduces measurable crosstalk between adjacent channels. The crosstalk arises from the combined influence of the PMT response time, the detection electronics, and the fluorescence lifetime of GCaMP6s (2.6-2.8 ns)^51^: the signal from one plane does not fully decay before the arrival of the subsequent excitation pulse. This incomplete decay leads to signal crosstalk between signals originating from different planes.

To quantitatively characterize the temporal crosstalk across beam channels, we performed a crosstalk ratio measurement informed by recent approaches to crosstalk characterization and correction in multi-beam imaging studies operating near the sampling limit^24, 79, 80^. The four channels were designated as *N-*1, *N*, *N*+1 and *N*+2, where *N*-1 and *N*+1 are adjacent to *N* and *N*+2 is separated by one plane. We selectively excited only channel *N* while simultaneously recording images from all four channels, and then calculated the crosstalk signal intensity in channels *N-*1, *N+*1 and *N*+2 as a percentage of the signal intensity in channel *N* (Extended Data Fig. 5B). Theoretically, at a 3.125-ns inter-pulse interval and given the fluorescence lifetime of GCaMP6s together with the aggregate bandwidth limit of the acquisition electronics, the temporal aliasing between adjacent channels is expected to exceed 30%. Our experimental measurements yielded contamination ratios of 36.2% and 36.5% for the two adjacent channels, whereas crosstalk into the non-adjacent channel (*N*+2) was substantially lower (∼ 4.7%).

Based on these empirically measured crosstalk coefficients, we constructed a channel unmixing matrix and applied a demixing pipeline to the volumetric imaging data (Extended Data Fig. 5C). Specifically, after equalizing the offsets across the four channels, we convolved the four raw pulse-train recordings with the unmixing matrix to estimate the demixed single-channel signals. The demixed images were subsequently processed with NeuroPixelAI to extract neuronal masks and their associated signals, which were then imported into Imaris for three-dimensional reconstruction.

To validate the effectiveness of this demixing strategy, we quantitatively assessed signal independence before and after correction. Extended Data Fig. 5C presents side-by-side neurons residing in distinct z-planes together with the degree of signal contamination before and after demixing. Following demixing, temporal crosstalk from adjacent channels was reduced. Furthermore, we computed the cross-correlation coefficient between neuronal signals extracted from channel *N* and those from channel *N*+1 (Extended Data Fig. 5D,E; n = 701 neurons for channel *N*, n = 167 neurons for channel *N*+1; two-sided paired t-tests, \*\*\*\**p* < 0.0001). The cross-correlation coefficient decreased significantly after demixing, demonstrating a substantial improvement in signal independence. The demixing procedure reduces temporal bleed-through between multiplexed channels but does not remove spatial overlap caused by the finite axial PSF. Neuronal masks were defined separately in Channel N (n = 701) and Channel N+1 (n = 167). For each mask, signals were extracted from both channels and the inter-channel correlation was calculated before and after demixing. Because the before- and after-demixing values were obtained from the same mask, comparisons were performed separately within each mask set using two-sided paired t-tests; the two mask sets were not compared with each other.

**Supplementary Note 5: Validation of the optogenetic stimulation module.**

As a wide-field two-photon microscopy system, Meso2P integrated with an optogenetic module enables cross-regional optogenetic stimulation in the mouse brain, providing a powerful tool for investigating interregional functional interactions across cortical areas.

For optogenetic validation experiments, 8-week-old male *Camk2a*-Cre::Ai162D mice were used. The cortical brain region was divided into four quadrants, and each quadrant was injected with 250 nL of C1V1 viral vector. Cranial window implantation was subsequently performed in all mice. After a 4-week recovery period to allow sufficient viral expression, viral transfection efficiency and optogenetic stimulation performance were systematically validated (Extended Data Fig. 7A-C).

The standardized optogenetic stimulation paradigm was implemented under 4× objective imaging. Briefly, laser stimulation was performed at a wavelength of 1030 nm and a power of 79 mW, delivered sequentially across the four brain quadrants in a cyclic order (quadrant i → ii → iii → iv → i). Each experimental session consisted of five stimulation cycles. A single quadrant stimulation lasted for 500 ms, composed of ten consecutive 50 ms log-spiral scans, with an inter-stimulus interval of 4000 ms between adjacent quadrants (Extended Data Fig. 7A).

Three stimulation targets were selected within each quadrant, with nearby cells were selected as internal controls, to verify the capability of the Meso2P system for spatially distributed multi-region optogenetic activation. The stimulated target sites across the four quadrants exhibited robust stimulation-locked calcium responses. In contrast, adjacent control cells displayed no periodic calcium fluctuations corresponding to the stimulation protocol (Extended Data Fig. 7D-F).

We further validated the compatibility and performance of the optogenetic module with the Meso2P system under 16× objective magnification. Consistent with the 4× imaging results, neurons receiving optogenetic stimulation exhibited stimulus-locked calcium transients, while neighboring control cells maintained spontaneous calcium dynamics without periodic activation responses (Extended Data Fig. 7G-H).

Collectively, these results validate spatially distributed optogenetic stimulation under the tested 4× configuration and stimulus-locked single-cell responses under the tested 16× configuration. They establish compatibility of the optogenetic module with both objectives under the reported conditions, without implying identical spatial precision or efficacy across the full mesoscopic FOV.

**Supplementary Note 6: Evaluation of system group delay dispersion**

This note provides a detailed evaluation of the group delay dispersion (GDD) of the Meso2P system, including the calculated material GDD budget, the effect of uncompensated residual dispersion on pulse width and two-photon excitation efficiency, arm-dependent dispersion in the spatiotemporal multiplexing module, and future optimization strategies.

## 1. Calculation of system material GDD

First, we listed all transmissive optical elements in the main optical path (Supplementary Fig. 1), including their material, thickness, and quantity. Using the Sellmeier equation, we calculated the group refractive index and group velocity dispersion (GVD) for each element at 920nm, and subsequently computed the total material GDD of the entire optical path (Supplementary Table 4).

**Supplementary Fig. 1.**
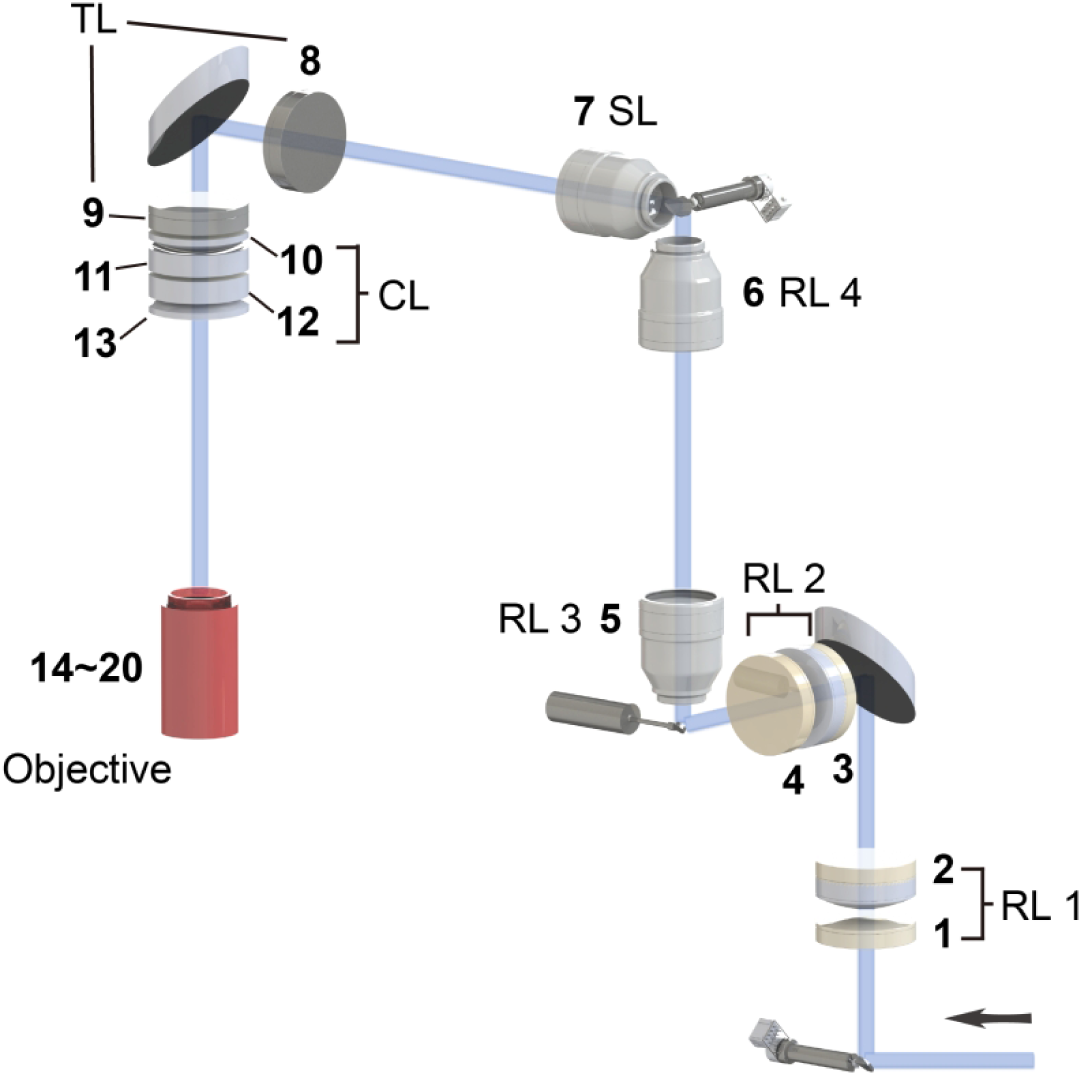
Arrangement of transmissive optical elements in the main optical path. (excluding the spatiotemporal multiplexing module)

The main optical path, without the spatiotemporal multiplexing module, introduces approximately 20,795 fs^2^ of material GDD. The first path in the temporal multiplexing module introduces an additional approximately 4,645 fs^2^ of GDD (Supplementary Table 5). Therefore, the total material GDD for the first multiplexed beam path is approximately 25,440 fs^2^.

The laser source used in our system is a Chameleon Discovery NX. In our instrument, the laser control software was set to a pre-chirp value of –17,358 fs^2^ at 920 nm (Supplementary Fig. 2). The available pre-compensation is not sufficient to fully compensate the total material GDD of our system.

**Supplementary Fig. 2.**
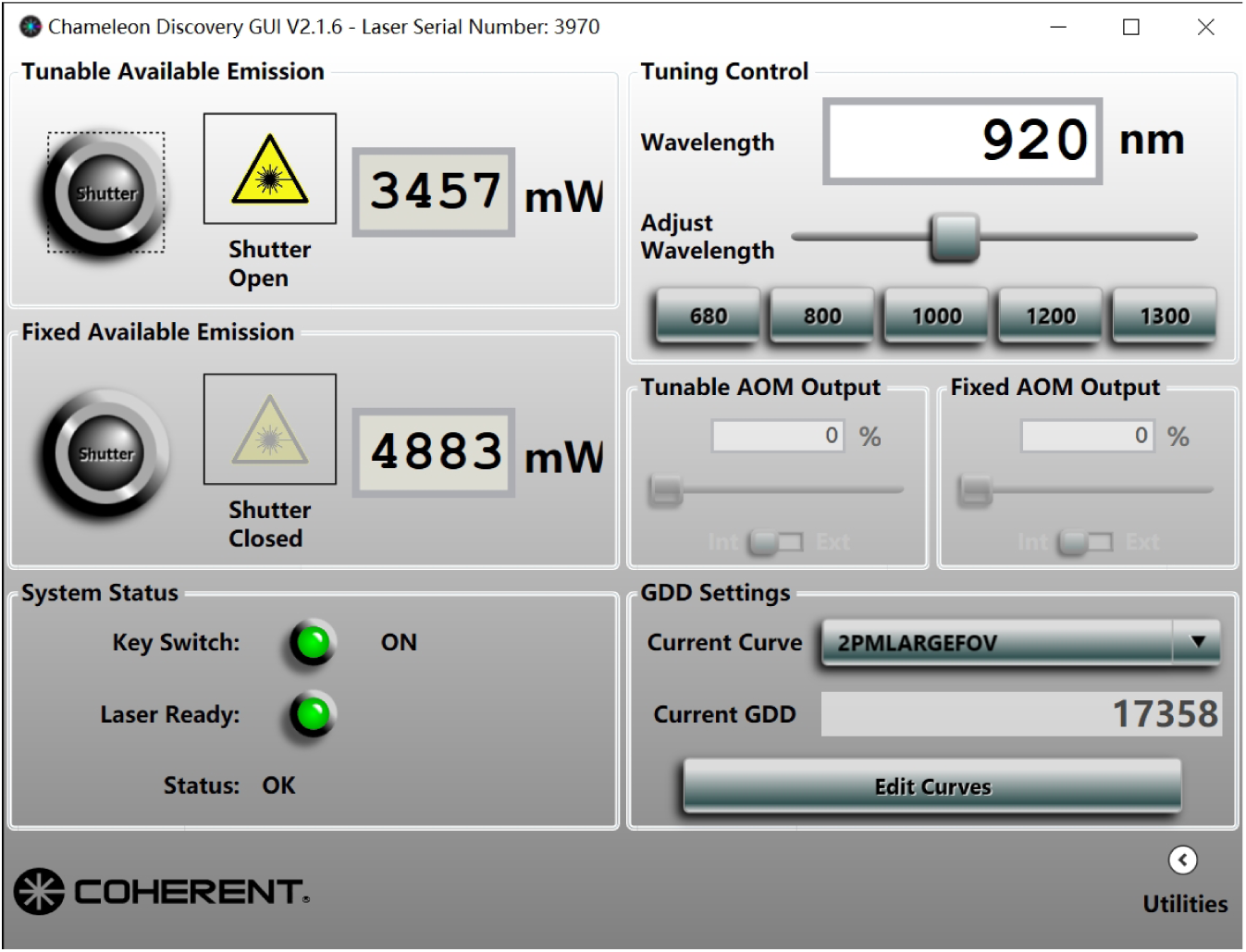
Laser setup parameters.

## 2. Effect of uncompensated residual GDD on pulse width and excitation efficiency

After applying the internal pre-chirp, the first multiplexed beam path still has an uncompensated GDD of approximately:

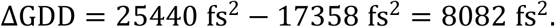

For a Gaussian pulse with an initial pulse width of 100 fs (FWHM), the broadened pulse width τ_out_ after propagating through the uncompensated GDD is given by:

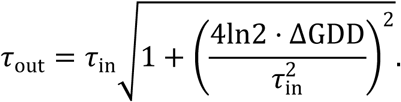

Substituting the values:

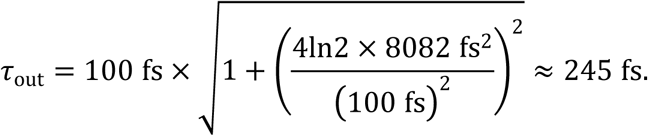

When the average power and repetition rate are fixed, the two-photon fluorescence excitation efficiency *F* is approximately inversely proportional to the pulse width τ^81^:

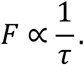

Thus, after pulse broadening from 100 fs to 245 fs, the two-photon excitation efficiency is reduced to:

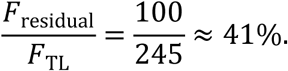

This corresponds to an approximately 2.4-fold reduction in excitation efficiency relative to the transform-limited case.

For two-photon excitation, the fluorescence intensity scales with the square of the average excitation power :

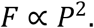

To achieve the same fluorescence yield, the required excitation power must be increased by a factor of:

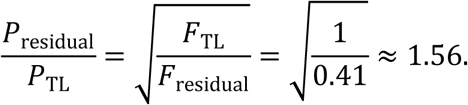

That is, the excitation power needs to be increased to 1.56 times that of the transform-limited case.

## 3. Arm-dependent dispersion in the spatiotemporal multiplexing module

In the multiplexing optical path, the four beam paths pass through different thicknesses of glass. Consequently, each beam path experiences a different amount of material GDD. We then listed all transmissive optical elements in the multiplexing optical path (Supplementary Fig. 3).

**Supplementary Fig. 3.**
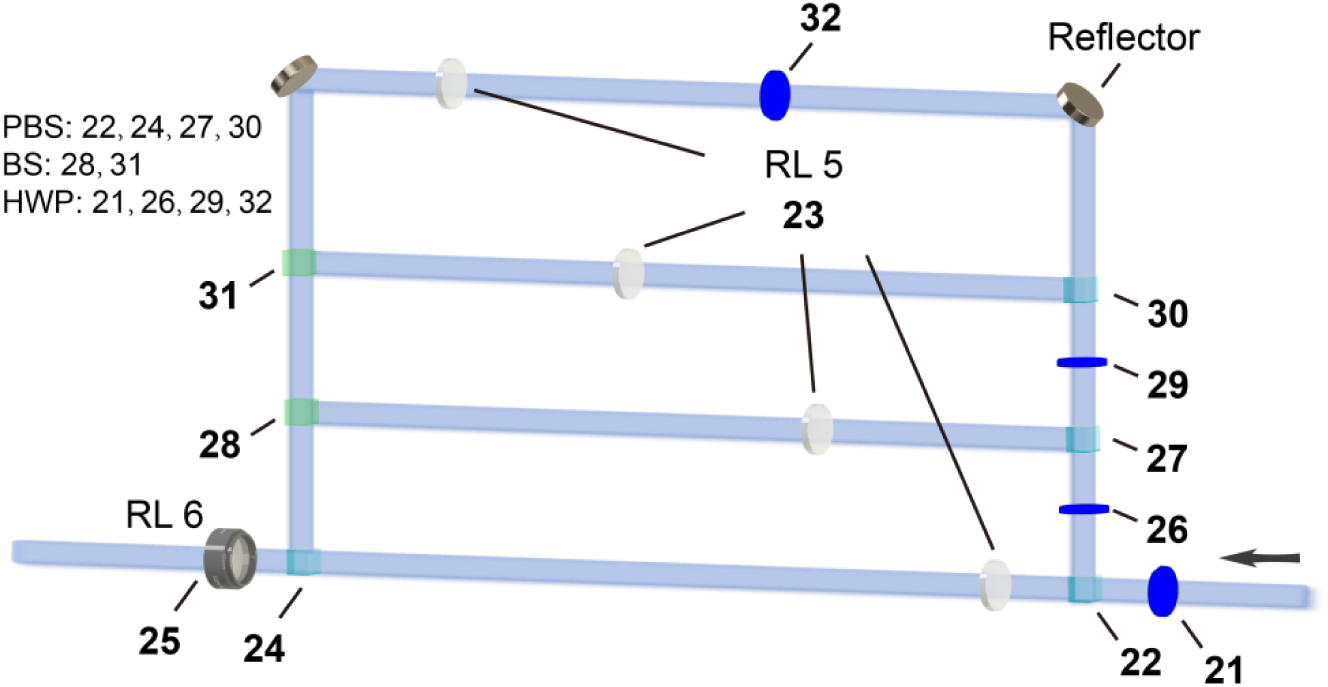
Arrangement of transmissive optical elements in the multiplexing optical path.

Relative to the first beam path, the other three beam paths introduce additional material GDD as Supplementary Table 6.

### Arm-dependent GDD and estimated pulse broadening

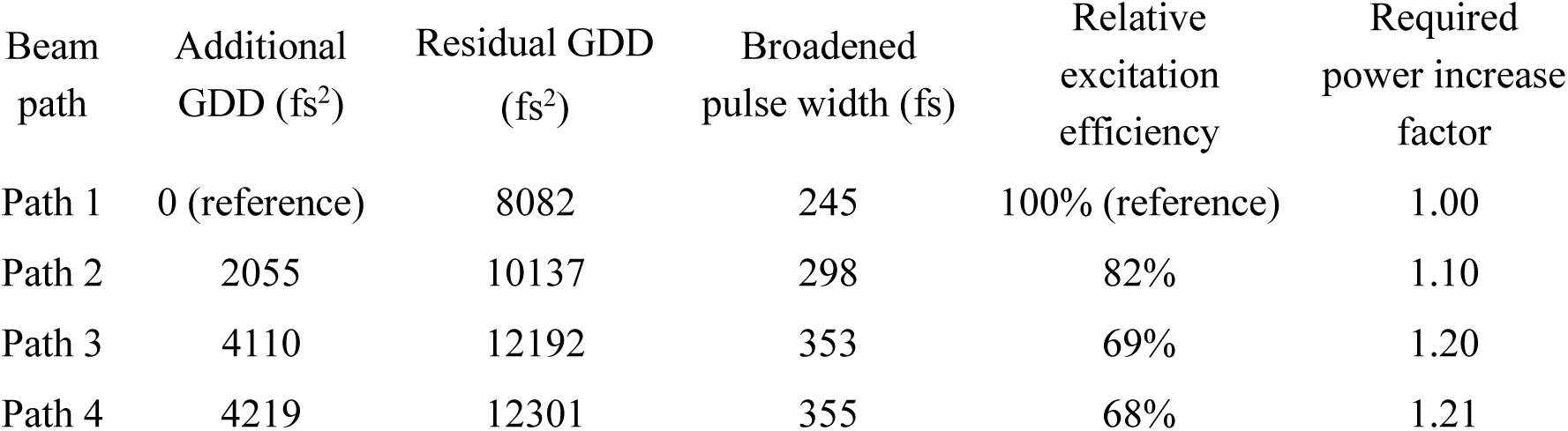

The broadened pulse widths were calculated using the same formula as in Section 2. The relative excitation efficiencies and the required power increase factors were calculated using the same relationships 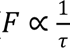 and *F* ⧜ *P*^2^ as in Section 2.

In the current system, we did not independently compensate the dispersion of each beam path. Instead, we used a single global pre-compensation from the laser and adjusted the excitation power of each beam path using the corresponding half-wave plates (HWPs). This power adjustment can balance the fluorescence levels between different imaging planes, but it does not remove the pulse broadening itself.

Nevertheless, the arm-dependent GDD should be explicitly quantified and reported.

## 4. Future optimization strategies

We identify this arm-dependent dispersion as a current limitation of Meso2P. In principle, additional glass elements of the appropriate thickness could be inserted into the shorter beam paths to equalize the total material GDD across all four paths. With a larger common pre-compensation range, all four paths could then be compensated together. Feasible optimization approaches include:

- Using a laser source with a larger pre-chirp range.
- Adding an external dispersion-compensation unit (e.g., a tunable prism pair or grating pair).

We plan to implement these strategies in future instrumental upgrades to reduce residual dispersion and arm-dependent GDD variations, thereby maximizing two-photon excitation efficiency across all beam paths.

### Associated display items

Supplementary Table 4, Supplementary Table 5.

